## Supplementary material for "Single molecule multi-omics reveals context-dependent regulation of enhancers by DNA methylation": LaTeX files: 5mC_citation_library.bib

@article{stadler_dna-binding_2011,
	title = {{DNA}-binding factors shape the mouse methylome at distal regulatory regions},
	year = {2011},
	volume = {480},
	issn = {1476-4687},
	doi = {10.1038/nature10716},
	abstract = {Methylation of cytosines is an essential epigenetic modification in mammalian genomes, yet the rules that govern methylation patterns remain largely elusive. To gain insights into this process, we generated base-pair-resolution mouse methylomes in stem cells and neuronal progenitors. Advanced quantitative analysis identified low-methylated regions ({LMRs}) with an average methylation of 30\%. These represent {CpG}-poor distal regulatory regions as evidenced by location, {DNase} I hypersensitivity, presence of enhancer chromatin marks and enhancer activity in reporter assays. {LMRs} are occupied by {DNA}-binding factors and their binding is necessary and sufficient to create {LMRs}. A comparison of neuronal and stem-cell methylomes confirms this dependency, as cell-type-specific {LMRs} are occupied by cell-type-specific transcription factors. This study provides methylome references for the mouse and shows that {DNA}-binding factors locally influence {DNA} methylation, enabling the identification of active regulatory regions.},
	pages = {490--495},
	number = {7378},
	journal = {Nature},
	shortjournal = {Nature},
	author = {Stadler, Michael B. and Murr, Rabih and Burger, Lukas and Ivanek, Robert and Lienert, Florian and Schöler, Anne and Nimwegen, Erik and Wirbelauer, Christiane and Oakeley, Edward J. and Gaidatzis, Dimos and Tiwari, Vijay K. and Schübeler, Dirk},
	date = {2011-12-14},
	pmid = {22170606},
}

@article{ziller_charting_2013,
	title = {Charting a dynamic {DNA} methylation landscape of the human genome},
	year = {2013},
	volume = {500},
	issn = {1476-4687},
	doi = {10.1038/nature12433},
	abstract = {Whole-genome bisulphite sequencing data from diverse human cell and tissue types shows that only about 22\% of {CpGs} change their methylation state across these cell types; most of these {CpGs} are located at gene regulatory elements, particularly enhancers and transcription-factor-binding sites, and these selected regions with dynamic {DNA} methylation patterns could help to define putative regulatory elements further.},
	pages = {477--481},
	number = {7463},
	journal = {Nature},
	author = {Ziller, Michael J. and Gu, Hongcang and Müller, Fabian and Donaghey, Julie and Tsai, Linus T.-Y. and Kohlbacher, Oliver and De Jager, Philip L. and Rosen, Evan D. and Bennett, David A. and Bernstein, Bradley E. and Gnirke, Andreas and Meissner, Alexander},
	date = {2013-08},
	langid = {english},
	note = {number: 7463
publisher: Nature Publishing Group},
}

@article{hodges_directional_2011,
	title = {Directional {DNA} methylation changes and complex intermediate states accompany lineage specificity in the adult hematopoietic compartment},
	year = {2011},
	volume = {44},
	issn = {1097-4164},
	doi = {10.1016/j.molcel.2011.08.026},
	abstract = {{DNA} methylation has been implicated as an epigenetic component of mechanisms that stabilize cell-fate decisions. Here, we have characterized the methylomes of human female hematopoietic stem/progenitor cells ({HSPCs}) and mature cells from the myeloid and lymphoid lineages. Hypomethylated regions ({HMRs}) associated with lineage-specific genes were often methylated in the opposing lineage. In {HSPCs}, these sites tended to show intermediate, complex patterns that resolve to uniformity upon differentiation, by increased or decreased methylation. Promoter {HMRs} shared across diverse cell types typically display a constitutive core that expands and contracts in a lineage-specific manner to fine-tune the expression of associated genes. Many newly identified intergenic {HMRs}, both constitutive and lineage specific, were enriched for factor binding sites with an implied role in genome organization and regulation of gene expression, respectively. Overall, our studies represent an important reference data set and provide insights into directional changes in {DNA} methylation as cells adopt terminal fates.},
	pages = {17--28},
	number = {1},
	journal = {Molecular Cell},
	shortjournal = {Mol Cell},
	author = {Hodges, Emily and Molaro, Antoine and Dos Santos, Camila O. and Thekkat, Pramod and Song, Qiang and Uren, Philip J. and Park, Jin and Butler, Jason and Rafii, Shahin and {McCombie}, W. Richard and Smith, Andrew D. and Hannon, Gregory J.},
	date = {2011-10-07},
	pmid = {21924933},
	pmcid = {PMC3412369},
}

@article{luo_dynamic_2018,
	title = {Dynamic {DNA} methylation: In the right place at the right time},
	year = {2018},
	volume = {361},
	issn = {0036-8075, 1095-9203},
	doi = {10.1126/science.aat6806},
	shorttitle = {Dynamic {DNA} methylation},
	abstract = {The classical model of cytosine {DNA} methylation (the presence of 5-methylcytosine, 5mC) regulation depicts this covalent modification as a stable repressive regulator of promoter activity. However, whole-genome analysis of 5mC reveals widespread tissue- and cell type–specific patterns and pervasive dynamics during mammalian development. Here we review recent findings that delineate 5mC functions in developmental stages and diverse genomic compartments as well as discuss the molecular mechanisms that connect transcriptional regulation and 5mC. Beyond the newly appreciated dynamics, regulatory roles for 5mC have been suggested in new biological contexts, such as learning and memory or aging. The use of new single-cell measurement techniques and precise editing tools will enable functional analyses of 5mC in gene expression, clarifying its role in various biological processes.},
	pages = {1336--1340},
	number = {6409},
	journal = {Science},
	shortjournal = {Science},
	author = {Luo, Chongyuan and Hajkova, Petra and Ecker, Joseph R.},
	date = {2018-09-28},
	langid = {english},
}

@article{iurlaro_dna_2017,
	title = {{DNA} methylation homeostasis in human and mouse development},
	year = {2017},
	volume = {43},
	issn = {1879-0380},
	doi = {10.1016/j.gde.2017.02.003},
	abstract = {The molecular pathways that regulate gain and loss of {DNA} methylation during mammalian development need to be tightly balanced to maintain a physiological equilibrium. Here we explore the relative contributions of the different pathways and enzymatic activities involved in methylation homeostasis in the context of genome-wide and locus-specific epigenetic reprogramming in mammals. An adaptable epigenetic machinery allows global epigenetic reprogramming to concur with local maintenance of critical epigenetic memory in the genome, and appears to regulate the tempo of global reprogramming in different cell lineages and species.},
	pages = {101--109},
	journal = {Current Opinion in Genetics \& Development},
	shortjournal = {Curr Opin Genet Dev},
	author = {Iurlaro, Mario and Meyenn, Ferdinand and Reik, Wolf},
	date = {2017-04},
	pmid = {28260631},
}

@article{ehrlich_amount_1982,
	title = {Amount and distribution of 5-methylcytosine in human {DNA} from different types of tissues of cells},
	year = {1982},
	volume = {10},
	issn = {0305-1048},
	doi = {10.1093/nar/10.8.2709},
	abstract = {Analysis of the total base composition of {DNA} from seven different normal human tissues and eight different types of homogeneous human cell populations revealed considerable tissue-specific and cell-specific differences in the extent of methylation of cytosine residues. The two most highly methylated {DNAs} were from thymus and brain with 1.00 and 0.98 mole percent 5-methylcytosine (m5C), respectively. The two least methylated {DNAs} from in vivo sources were placental {DNA} and sperm {DNA}, which had 0.76 and 0.84 mole percent m5C, respectively. The differences between these two groups of samples were significant with p less than 0.01. The m5C content of {DNA} from six human cell lines or strains ranged from 0.57 to 0.85 mole percent. The major and minor base composition of {DNA} fractionated by reassociation kinetics was also determined. The distribution of m5C among these fractions showed little or no variation with tissue or cell type with the possible exception of sperm {DNA}. In each case, nonrepetitive {DNA} sequences were hypomethylated compared to unfractionated {DNA}.},
	pages = {2709--2721},
	number = {8},
	journal = {Nucleic Acids Research},
	shortjournal = {Nucleic Acids Res},
	author = {Ehrlich, M. and Gama-Sosa, M. A. and Huang, L. H. and Midgett, R. M. and Kuo, K. C. and {McCune}, R. A. and Gehrke, C.},
	date = {1982-04-24},
	pmid = {7079182},
	pmcid = {PMC320645},
}

@article{lister_human_2009,
	title = {Human {DNA} methylomes at base resolution show widespread epigenomic differences},
	year = {2009},
	volume = {462},
	issn = {1476-4687},
	doi = {10.1038/nature08514},
	abstract = {{DNA} cytosine methylation is a central epigenetic modification that has essential roles in cellular processes including genome regulation, development and disease. Here we present the first genome-wide, single-base-resolution maps of methylated cytosines in a mammalian genome, from both human embryonic stem cells and fetal fibroblasts, along with comparative analysis of messenger {RNA} and small {RNA} components of the transcriptome, several histone modifications, and sites of {DNA}–protein interaction for several key regulatory factors. Widespread differences were identified in the composition and patterning of cytosine methylation between the two genomes. Nearly one-quarter of all methylation identified in embryonic stem cells was in a non-{CG} context, suggesting that embryonic stem cells may use different methylation mechanisms to affect gene regulation. Methylation in non-{CG} contexts showed enrichment in gene bodies and depletion in protein binding sites and enhancers. Non-{CG} methylation disappeared upon induced differentiation of the embryonic stem cells, and was restored in induced pluripotent stem cells. We identified hundreds of differentially methylated regions proximal to genes involved in pluripotency and differentiation, and widespread reduced methylation levels in fibroblasts associated with lower transcriptional activity. These reference epigenomes provide a foundation for future studies exploring this key epigenetic modification in human disease and development.},
	pages = {315--322},
	number = {7271},
	journal = {Nature},
	author = {Lister, Ryan and Pelizzola, Mattia and Dowen, Robert H. and Hawkins, R. David and Hon, Gary and Tonti-Filippini, Julian and Nery, Joseph R. and Lee, Leonard and Ye, Zhen and Ngo, Que-Minh and Edsall, Lee and Antosiewicz-Bourget, Jessica and Stewart, Ron and Ruotti, Victor and Millar, A. Harvey and Thomson, James A. and Ren, Bing and Ecker, Joseph R.},
	date = {2009-11},
	langid = {english},
	note = {number: 7271
publisher: Nature Publishing Group},
}

@article{meissner_genome-scale_2008,
	title = {Genome-scale {DNA} methylation maps of pluripotent and differentiated cells},
	year = {2008},
	volume = {454},
	issn = {1476-4687},
	doi = {10.1038/nature07107},
	abstract = {{DNA} methylation, an important mechanism of epigenetic modification that produces different patterns of gene expression from a single {DNA} sequence, is vital to normal development and its malfunction can cause cancer and other abnormalities. A map of {DNA} methylation in embryonic stem cells, and in various cell types derived from them, has now been produced at nucleotide resolution using high-throughput bisulphite sequencing combined with single molecule-based sequencing. The map reveals specific sites in the genome where methylation changes as cells develop, for instance when embryonic stem cells mature into nerve cells. More generally, the methodology will be of value for the epigenetic profiling of cell populations relevant to developmental biology, cancer and regenerative medicine.},
	pages = {766--770},
	number = {7205},
	journal = {Nature},
	author = {Meissner, Alexander and Mikkelsen, Tarjei S. and Gu, Hongcang and Wernig, Marius and Hanna, Jacob and Sivachenko, Andrey and Zhang, Xiaolan and Bernstein, Bradley E. and Nusbaum, Chad and Jaffe, David B. and Gnirke, Andreas and Jaenisch, Rudolf and Lander, Eric S.},
	date = {2008-08},
	langid = {english},
	note = {number: 7205
publisher: Nature Publishing Group},
}

@article{greenberg_diverse_2019,
	title = {The diverse roles of {DNA} methylation in mammalian development and disease},
	year = {2019},
	volume = {20},
	issn = {1471-0072, 1471-0080},
	doi = {10.1038/s41580-019-0159-6},
	abstract = {{DNA} methylation is of paramount importance for mammalian embryonic development. {DNA} methylation has numerous functions: it is implicated in the repression of transposons and genes, but is also associated with actively transcribed gene bodies and, in some cases, with gene activation per se. In recent years, sensitive technologies have been developed that allow the interrogation of {DNA} methylation patterns from a small number of cells. The use of these technol­ ogies has greatly improved our knowledge of {DNA} methylation dynamics and heterogeneity in embryos and in specific tissues. Combined with genetic analyses, it is increasingly apparent that regulation of {DNA} methylation erasure and (re-)establishment varies considerably between different developmental stages. In this Review , we discuss the mechanisms and functions of {DNA} methylation and demethylation in both mice and humans at {CpG}-rich promoters, gene bodies and transposable elements. We highlight the dynamic erasure and re-establishment of {DNA} methyl­ation in embryonic, germline and somatic cell development. Finally , we provide insights into {DNA} methylation gained from studying genetic diseases.},
	pages = {590--607},
	number = {10},
	journal = {Nature Reviews Molecular Cell Biology},
	shortjournal = {Nat Rev Mol Cell Biol},
	author = {Greenberg, Maxim V. C. and Bourc’his, Deborah},
	date = {2019-10},
	langid = {english},
}

@article{dahlet_genome-wide_2020,
	title = {Genome-wide analysis in the mouse embryo reveals the importance of {DNA} methylation for transcription integrity},
	year = {2020},
	volume = {11},
	issn = {2041-1723},
	doi = {10.1038/s41467-020-16919-w},
	abstract = {Abstract
            
              Mouse embryos acquire global {DNA} methylation of their genome during implantation. However the exact roles of {DNA} methyltransferases ({DNMTs}) in embryos have not been studied comprehensively. Here we systematically analyze the consequences of genetic inactivation of
              Dnmt1
              ,
              Dnmt3a
              and
              Dnmt3b
              on the methylome and transcriptome of mouse embryos. We find a strict division of function between {DNMT}1, responsible for maintenance methylation, and {DNMT}3A/B, solely responsible for methylation acquisition in development. By analyzing severely hypomethylated embryos, we uncover multiple functions of {DNA} methylation that is used as a mechanism of repression for a panel of genes including not only imprinted and germline genes, but also lineage-committed genes and 2-cell genes. {DNA} methylation also suppresses multiple retrotransposons and illegitimate transcripts from cryptic promoters in transposons and gene bodies. Our work provides a thorough analysis of the roles of {DNA} methyltransferases and the importance of {DNA} methylation for transcriptome integrity in mammalian embryos.},
	pages = {3153},
	number = {1},
	journal = {Nature Communications},
	shortjournal = {Nat Commun},
	author = {Dahlet, Thomas and Argüeso Lleida, Andrea and Al Adhami, Hala and Dumas, Michael and Bender, Ambre and Ngondo, Richard P. and Tanguy, Manon and Vallet, Judith and Auclair, Ghislain and Bardet, Anaïs F. and Weber, Michael},
	date = {2020-12},
	langid = {english},
}

@article{li_targeted_1992,
	title = {Targeted mutation of the {DNA} methyltransferase gene results in embryonic lethality},
	year = {1992},
	volume = {69},
	issn = {00928674},
	doi = {10.1016/0092-8674(92)90611-F},
	pages = {915--926},
	number = {6},
	journal = {Cell},
	shortjournal = {Cell},
	author = {Li, En and Bestor, Timothy H. and Jaenisch, Rudolf},
	date = {1992-06},
	langid = {english},
}

@article{sakaue_dna_2010,
	title = {{DNA} Methylation Is Dispensable for the Growth and Survival of the Extraembryonic Lineages},
	year = {2010},
	volume = {20},
	issn = {0960-9822},
	doi = {10.1016/j.cub.2010.06.050},
	abstract = {{DNA} methylation regulates development and many epigenetic processes in mammals [1], and it is required for somatic cell growth and survival [2, 3]. In contrast, embryonic stem ({ES}) cells can self-renew without {DNA} methylation [4, 5, 6]. It remains unclear whether any lineage-committed cells can survive without {DNA}-methylation machineries. Unlike in somatic cells, {DNA} methylation is dispensable for imprinting and X-inactivation in the extraembryonic lineages [7, 8, 9, 10, 11, 12]. In {ES} cells, {DNA} methylation prevents differentiation into the trophectodermal fate [13]. Here, we created triple-knockout ({TKO}) mouse embryos deficient for the active {DNA} methyltransferases Dnmt1, Dnmt3a, and Dnmt3b ({TKO}) by nuclear transfer ({NT}), and we examined their development. In chimeric {TKO}-{NT} and {WT} embryos, few {TKO} cells were found in the embryo proper, but they contributed to extraembryonic tissues. {TKO} {ES} cells showed increasing cell death during their differentiation into epiblast lineages, but not during differentiation into extraembryonic lineages. Furthermore, we successfully established trophoblastic stem cells ({ntTS} cells) from {TKO}-{NT} blastocysts. These {TKO} {ntTS} cells could self-renew, and they retained the fundamental gene expression patterns of stem cells. Our findings indicated that extraembryonic-lineage cells can survive and proliferate in the absence of {DNA} methyltransferases and that a cell's response to the stress of epigenomic damage is cell type dependent.},
	pages = {1452--1457},
	number = {16},
	journal = {Current Biology},
	shortjournal = {Current Biology},
	author = {Sakaue, Morito and Ohta, Hiroshi and Kumaki, Yuichi and Oda, Masaaki and Sakaide, Yuko and Matsuoka, Chisa and Yamagiwa, Akiko and Niwa, Hitoshi and Wakayama, Teruhiko and Okano, Masaki},
	date = {2010-08-24},
	langid = {english},
}

@article{baubec_genomic_2014,
	title = {Genomic patterns and context specific interpretation of {DNA} methylation},
	year = {2014},
	volume = {25},
	issn = {0959-437X},
	doi = {10.1016/j.gde.2013.11.015},
	abstract = {Methylation of {CpG} dinucleotides is a reversible modification of {DNA} that is highly prevalent throughout mammalian genomes. Recent advances generated genomic {DNA} methylation maps during cellular differentiation at unprecedented resolution. Combined with functional assays this revealed that dynamics in {DNA} methylation coincide with changes in regulatory activity and that transcription factors play an important role in shaping methylation patterns. This tightly links {DNA} methylation with underlying {DNA} sequence features and suggests that a substantial fraction of methylation changes occur downstream of gene regulation. Here we discuss our current understanding of the context-dependent readout of {DNA} methylation and criteria that need to be fulfilled for this modification to be instructive for gene regulation.},
	pages = {85--92},
	journal = {Current Opinion in Genetics \& Development},
	shortjournal = {Current Opinion in Genetics \& Development},
	author = {Baubec, Tuncay and Schübeler, Dirk},
	date = {2014-04-01},
	langid = {english},
}

@article{kribelbauer_toward_2020,
	title = {Toward a Mechanistic Understanding of {DNA} Methylation Readout by Transcription Factors},
	year = {2020},
	volume = {432},
	issn = {0022-2836},
	doi = {10.1016/j.jmb.2019.10.021},
	abstract = {Epigenetic {DNA} modification impacts gene expression, but the underlying molecular mechanisms are only partly understood. Adding a methyl group to a cytosine base locally modifies the structural features of {DNA} in multiple ways, which may change the interaction with {DNA}-binding transcription factors ({TFs}) and trigger a cascade of downstream molecular events. Cells can be probed using various functional genomics assays, but it is difficult to disentangle the confounded effects of {DNA} modification on {TF} binding, chromatin accessibility, intranuclear variation in local {TF} concentration, and rate of transcription. Here we discuss how high-throughput in vitro profiling of protein–{DNA} interactions has enabled comprehensive characterization and quantification of the methylation sensitivity of {TFs}. Despite the limited structural data for {DNA} containing methylated cytosine, automated analysis of structural information in the Protein Data Bank ({PDB}) shows how 5-methylcytosine (5mC) can be recognized in various ways by amino acid side chains. We discuss how a context-dependent effect of methylation on {DNA} groove geometry can affect {DNA} binding by homeodomain proteins and how principled modeling of {ChIP}-seq data can overcome the confounding that makes the interpretation of in vivo data challenging. The emerging picture is that epigenetic modifications affect {TF} binding in a highly context-specific manner, with a direction and effect size that depend critically on their position within the {TF} binding site and the amino acid sequence of the {TF}. With this improved mechanistic knowledge, we have come closer to understanding how cells use {DNA} modification to acquire, retain, and change their identity.},
	pages = {1801--1815},
	number = {6},
	journal = {Journal of Molecular Biology},
	shortjournal = {Journal of Molecular Biology},
	author = {Kribelbauer, Judith F. and Lu, Xiang-Jun and Rohs, Remo and Mann, Richard S. and Bussemaker, Harmen J.},
	date = {2020-03-13},
	langid = {english},
}

@article{kribelbauer_quantitative_2017,
	title = {Quantitative Analysis of the {DNA} Methylation Sensitivity of Transcription Factor Complexes},
	year = {2017},
	volume = {19},
	issn = {2211-1247},
	doi = {10.1016/j.celrep.2017.05.069},
	abstract = {Although {DNA} modifications play an important role in gene regulation, the underlying mechanisms remain elusive. We developed {EpiSELEX}-seq to probe the sensitivity of transcription factor binding to {DNA} modification in vitro using massively parallel sequencing. Feature-based modeling quantifies the effect of cytosine methylation (5mC) on binding free energy in a position-specific manner. Application to the human {bZIP} proteins {ATF}4 and C/{EBPβ} and three different Pbx-Hox complexes shows that 5mCpG can both increase and decrease affinity, depending on where the modification occurs within the protein-{DNA} interface. The {TF} paralogs tested vary in their methylation sensitivity, for which we provide a structural rationale. We show that 5mCpG can also enhance in vitro p53 binding and provide evidence for increased in vivo p53 occupancy at methylated binding sites, correlating with primed enhancer histone marks. Our results establish a powerful strategy for dissecting the epigenomic modulation of protein-{DNA} interactions and their role in gene regulation.},
	pages = {2383--2395},
	number = {11},
	journal = {Cell Reports},
	shortjournal = {Cell Rep},
	author = {Kribelbauer, Judith F. and Laptenko, Oleg and Chen, Siying and Martini, Gabriella D. and Freed-Pastor, William A. and Prives, Carol and Mann, Richard S. and Bussemaker, Harmen J.},
	date = {2017-06-13},
	pmid = {28614722},
	pmcid = {PMC5533174},
}

@article{yin_impact_2017,
	title = {Impact of cytosine methylation on {DNA} binding specificities of human transcription factors},
	year = {2017},
	volume = {356},
	issn = {0036-8075, 1095-9203},
	doi = {10.1126/science.aaj2239},
	pages = {eaaj2239},
	number = {6337},
	journal = {Science},
	shortjournal = {Science},
	author = {Yin, Yimeng and Morgunova, Ekaterina and Jolma, Arttu and Kaasinen, Eevi and Sahu, Biswajyoti and Khund-Sayeed, Syed and Das, Pratyush K. and Kivioja, Teemu and Dave, Kashyap and Zhong, Fan and Nitta, Kazuhiro R. and Taipale, Minna and Popov, Alexander and Ginno, Paul A. and Domcke, Silvia and Yan, Jian and Schübeler, Dirk and Vinson, Charles and Taipale, Jussi},
	date = {2017-05-05},
	langid = {english},
}

@article{domcke_competition_2015,
	title = {Competition between {DNA} methylation and transcription factors determines binding of {NRF}1},
	year = {2015},
	volume = {528},
	issn = {0028-0836, 1476-4687},
	doi = {10.1038/nature16462},
	pages = {575--579},
	number = {7583},
	journal = {Nature},
	shortjournal = {Nature},
	author = {Domcke, Silvia and Bardet, Anaïs Flore and Adrian Ginno, Paul and Hartl, Dominik and Burger, Lukas and Schübeler, Dirk},
	date = {2015-12},
	langid = {english},
}

@article{maurano_role_2015,
	title = {Role of {DNA} Methylation in Modulating Transcription Factor Occupancy},
	year = {2015},
	volume = {12},
	issn = {2211-1247},
	doi = {10.1016/j.celrep.2015.07.024},
	pages = {1184--1195},
	number = {7},
	journal = {Cell Reports},
	shortjournal = {Cell Reports},
	author = {Maurano, Matthew T. and Wang, Hao and John, Sam and Shafer, Anthony and Canfield, Theresa and Lee, Kristen and Stamatoyannopoulos, John A.},
	date = {2015-08-18},
	pmid = {26257180},
	note = {publisher: Elsevier},
}

@article{hartl_cis-regulatory_2017,
	title = {Cis-regulatory landscapes of four cell types of the retina},
	year = {2017},
	volume = {45},
	issn = {1362-4962},
	doi = {10.1093/nar/gkx923},
	abstract = {The retina is composed of ∼50 cell-types with specific functions for the process of vision. Identification of the cis-regulatory elements active in retinal cell-types is key to elucidate the networks controlling this diversity. Here, we combined transcriptome and epigenome profiling to map the regulatory landscape of four cell-types isolated from mouse retinas including rod and cone photoreceptors as well as rare inter-neuron populations such as horizontal and starburst amacrine cells. Integration of this information reveals sequence determinants and candidate transcription factors for controlling cellular specialization. Additionally, we refined parallel reporter assays to enable studying the transcriptional activity of large collection of sequences in individual cell-types isolated from a tissue. We provide proof of concept for this approach and its scalability by characterizing the transcriptional capacity of several hundred putative regulatory sequences within individual retinal cell-types. This generates a catalogue of cis-regulatory regions active in retinal cell types and we further demonstrate their utility as potential resource for cellular tagging and manipulation.},
	pages = {11607--11621},
	number = {20},
	journal = {Nucleic Acids Research},
	shortjournal = {Nucleic Acids Res},
	author = {Hartl, Dominik and Krebs, Arnaud R. and Jüttner, Josephine and Roska, Botond and Schübeler, Dirk},
	date = {2017-11-16},
	pmid = {29059322},
	pmcid = {PMC5714137},
}

@article{parry_active_2021,
	title = {Active turnover of {DNA} methylation during cell fate decisions},
	year = {2021},
	volume = {22},
	issn = {1471-0056, 1471-0064},
	doi = {10.1038/s41576-020-00287-8},
	abstract = {{DNA} methylation is a key layer of epigenetic regulation. The deposition of methylation marks relies on the catalytic activity of {DNA} methyltransferases ({DNMTs}), and their active removal relies on the activity of ten–eleven translocation ({TET}) enzymes. Paradoxically, in important biological contexts these antagonistic factors are co-expressed and target overlapping genomic regions. The ensuing cyclic biochemistry of cytosine modifications gives rise to a continuous, out-of-thermal equilibrium transition through different methylation states. But what is the purpose of this intriguing turnover of {DNA} methylation? Recent evidence demonstrates that methylation turnover is enriched at gene distal regulatory elements, including enhancers, and can give rise to large-scale oscillatory dynamics. We discuss this phenomenon and propose that {DNA} methylation turnover might facilitate key lineage decisions.},
	pages = {59--66},
	number = {1},
	journal = {Nature Reviews Genetics},
	shortjournal = {Nat Rev Genet},
	author = {Parry, Aled and Rulands, Steffen and Reik, Wolf},
	date = {2021-01},
	langid = {english},
}

@article{ginno_genome-scale_2020,
	title = {A genome-scale map of {DNA} methylation turnover identifies site-specific dependencies of {DNMT} and {TET} activity},
	year = {2020},
	volume = {11},
	issn = {2041-1723},
	doi = {10.1038/s41467-020-16354-x},
	abstract = {{DNA} methylation is considered a stable epigenetic mark, yet methylation patterns can vary during differentiation and in diseases such as cancer. Local levels of {DNA} methylation result from opposing enzymatic activities, the rates of which remain largely unknown. Here we developed a theoretical and experimental framework enabling us to infer methylation and demethylation rates at 860,404 {CpGs} in mouse embryonic stem cells. We find that enzymatic rates can vary as much as two orders of magnitude between {CpGs} with identical steady-state {DNA} methylation. Unexpectedly, de novo and maintenance methylation activity is reduced at transcription factor binding sites, while methylation turnover is elevated in transcribed gene bodies. Furthermore, we show that {TET} activity contributes substantially more than passive demethylation to establishing low methylation levels at distal enhancers. Taken together, our work unveils a genome-scale map of methylation kinetics, revealing highly variable and context-specific activity for the {DNA} methylation machinery.},
	pages = {2680},
	number = {1},
	journal = {Nature Communications},
	shortjournal = {Nat Commun},
	author = {Ginno, Paul Adrian and Gaidatzis, Dimos and Feldmann, Angelika and Hoerner, Leslie and Imanci, Dilek and Burger, Lukas and Zilbermann, Frederic and Peters, Antoine H. F. M. and Edenhofer, Frank and Smallwood, Sébastien A. and Krebs, Arnaud R. and Schübeler, Dirk},
	date = {2020-05-29},
	pmid = {32471981},
	pmcid = {PMC7260214},
}

@article{lu_role_2014,
	title = {Role of Tet proteins in enhancer activity and telomere elongation},
	year = {2014},
	volume = {28},
	issn = {0890-9369, 1549-5477},
	doi = {10.1101/gad.248005.114},
	pages = {2103--2119},
	number = {19},
	journal = {Genes \& Development},
	shortjournal = {Genes Dev.},
	author = {Lu, Falong and Liu, Yuting and Jiang, Lan and Yamaguchi, Shinpei and Zhang, Yi},
	date = {2014-10-01},
	langid = {english},
}

@article{landan_epigenetic_2012,
	title = {Epigenetic polymorphism and the stochastic formation of differentially methylated regions in normal and cancerous tissues},
	year = {2021},
	volume = {44},
	issn = {1061-4036, 1546-1718},
	doi = {10.1038/ng.2442},
	pages = {1207--1214},
	number = {11},
	journal = {Nature Genetics},
	shortjournal = {Nat Genet},
	author = {Landan, Gilad and Cohen, Netta Mendelson and Mukamel, Zohar and Bar, Amir and Molchadsky, Alina and Brosh, Ran and Horn-Saban, Shirley and Zalcenstein, Daniela Amann and Goldfinger, Naomi and Zundelevich, Adi and Gal-Yam, Einav Nili and Rotter, Varda and Tanay, Amos},
	date = {2012-11},
	langid = {english},
}

@article{shipony_dynamic_2014,
	title = {Dynamic and static maintenance of epigenetic memory in pluripotent and somatic cells},
	year = {2014},
	volume = {513},
	issn = {0028-0836, 1476-4687},
	doi = {10.1038/nature13458},
	pages = {115--119},
	number = {7516},
	journal = {Nature},
	shortjournal = {Nature},
	author = {Shipony, Zohar and Mukamel, Zohar and Cohen, Netta Mendelson and Landan, Gilad and Chomsky, Elad and Zeliger, Shlomit Reich and Fried, Yael Chagit and Ainbinder, Elena and Friedman, Nir and Tanay, Amos},
	date = {2014-09},
	langid = {english},
}

@article{farlik_single-cell_2015,
	title = {Single-Cell {DNA} Methylome Sequencing and Bioinformatic Inference of Epigenomic Cell-State Dynamics},
	year = {2015},
	volume = {10},
	issn = {2211-1247},
	doi = {10.1016/j.celrep.2015.02.001},
	abstract = {Methods for single-cell genome and transcriptome sequencing have contributed to our understanding of cellular heterogeneity, whereas methods for single-cell epigenomics are much less established. Here, we describe a whole-genome bisulfite sequencing ({WGBS}) assay that enables {DNA} methylation mapping in very small cell populations (μ{WGBS}) and single cells ({scWGBS}). Our assay is optimized for profiling many samples at low coverage, and we describe a bioinformatic method that analyzes collections of single-cell methylomes to infer cell-state dynamics. Using these technological advances, we studied epigenomic cell-state dynamics in three in vitro models of cellular differentiation and pluripotency, where we observed characteristic patterns of epigenome remodeling and cell-to-cell heterogeneity. The described method enables single-cell analysis of {DNA} methylation in a broad range of biological systems, including embryonic development, stem cell differentiation, and cancer. It can also be used to establish composite methylomes that account for cell-to-cell heterogeneity in complex tissue samples., 
          
            
              •
              High-throughput bisulfite sequencing assay for low-input and single-cell samples
            
            
              •
              Single-cell methylomes for in vitro models (K562 {AzaC}, {HL}60 {VitD}, and {mES} 2i/{ATRA}/{EB})
            
            
              •
              Bioinformatic method for inferring cell-state dynamics from sparse methylome data
            
            
              •
              Identification of genomic region types with consistent changes among single cells
            
          
        , Farlik et al. describe a method for {DNA} methylation sequencing in very small cell populations (μ{WGBS}) and single cells ({scWGBS}). Furthermore, they present a bioinformatic method for analyzing low-coverage methylome data and apply this technique to inferring epigenomic cell-state dynamics in pluripotent and differentiating cells.},
	pages = {1386--1397},
	number = {8},
	journal = {Cell Reports},
	shortjournal = {Cell Rep},
	author = {Farlik, Matthias and Sheffield, Nathan C. and Nuzzo, Angelo and Datlinger, Paul and Schönegger, Andreas and Klughammer, Johanna and Bock, Christoph},
	date = {2015-02-26},
	pmid = {25732828},
	pmcid = {PMC4542311},
}

@article{smallwood_single-cell_2014,
	title = {Single-cell genome-wide bisulfite sequencing for assessing epigenetic heterogeneity},
	year = {2014},
	volume = {11},
	issn = {1548-7105},
	doi = {10.1038/nmeth.3035},
	abstract = {Single-cell bisulfite sequencing ({scBS}-seq) allows robust {DNA} methylation analysis in rare cells and heterogeneous populations.},
	pages = {817--820},
	number = {8},
	journal = {Nature Methods},
	author = {Smallwood, Sébastien A. and Lee, Heather J. and Angermueller, Christof and Krueger, Felix and Saadeh, Heba and Peat, Julian and Andrews, Simon R. and Stegle, Oliver and Reik, Wolf and Kelsey, Gavin},
	date = {2014-08},
	langid = {english},
	note = {number: 8
publisher: Nature Publishing Group},
}

@article{song_dynamic_2019,
	title = {Dynamic Enhancer {DNA} Methylation as Basis for Transcriptional and Cellular Heterogeneity of {ESCs}},
	year = {2019},
	volume = {75},
	issn = {10972765},
	doi = {10.1016/j.molcel.2019.06.045},
	abstract = {Variable levels of {DNA} methylation have been reported at tissue-speciﬁc differential methylation regions ({DMRs}) overlapping enhancers, including super-enhancers ({SEs}) associated with key cell identity genes, but the mechanisms responsible for this intriguing behavior are not well understood. We used allele-speciﬁc reporters at the endogenous Sox2 and Mir290 {SEs} in embryonic stem cells and found that the allelic {DNA} methylation state is dynamically switching, resulting in cell-to-cell heterogeneity. Dynamic {DNA} methylation is driven by the balance between {DNA} methyltransferases and transcription factor binding on one side and co-regulated with the Mediator complex recruitment and H3K27ac level changes at regulatory elements on the other side. {DNA} methylation at the Sox2 and the Mir290 {SEs} is independently regulated and has distinct consequences on the cellular differentiation state. Dynamic allele-speciﬁc {DNA} methylation at the two {SEs} was also seen at different stages in preimplantation embryos, revealing that methylation heterogeneity occurs in vivo.},
	pages = {905--920.e6},
	number = {5},
	journal = {Molecular Cell},
	shortjournal = {Molecular Cell},
	author = {Song, Yuelin and Berg, Patrick R. and Markoulaki, Styliani and Soldner, Frank and Dall’Agnese, Alessandra and Henninger, Jonathan E. and Drotar, Jesse and Rosenau, Nicholas and Cohen, Malkiel A. and Young, Richard A. and Semrau, Stefan and Stelzer, Yonatan and Jaenisch, Rudolf},
	date = {2019-09},
	langid = {english},
}

@article{barnett_atac-me_2020,
	title = {{ATAC}-Me Captures Prolonged {DNA} Methylation of Dynamic Chromatin Accessibility Loci during Cell Fate Transitions},
	year = {2020},
	volume = {77},
	issn = {10972765},
	doi = {10.1016/j.molcel.2020.01.004},
	abstract = {{DNA} methylation of enhancers is dynamic, cell-type speciﬁc, and vital for cell fate progression. However, current models inadequately deﬁne its role within the hierarchy of gene regulation. Analysis of independent datasets shows an unanticipated overlap between {DNA} methylation and chromatin accessibility at enhancers of steady-state stem cells, suggesting that these two opposing features might exist concurrently. To deﬁne their temporal relationship, we developed {ATAC}-Me, which probes accessibility and methylation from single {DNA} library preparations. We identiﬁed waves of accessibility occurring rapidly across thousands of myeloid enhancers in a monocyte-to-macrophage cell fate model. Prolonged methylation states were observed at a majority of these sites, while transcription of nearby genes tracked closely with accessibility. {ATAC}-Me uncovers a signiﬁcant disconnect between chromatin accessibility, {DNA} methylation status, and gene activity. This unexpected observation highlights the value of {ATAC}-Me in constructing precise molecular timelines for understanding the role of {DNA} methylation in gene regulation.},
	pages = {1350--1364.e6},
	number = {6},
	journal = {Molecular Cell},
	shortjournal = {Molecular Cell},
	author = {Barnett, Kelly R. and Decato, Benjamin E. and Scott, Timothy J. and Hansen, Tyler J. and Chen, Bob and Attalla, Jonathan and Smith, Andrew D. and Hodges, Emily},
	date = {2020-03},
	langid = {english},
}

@article{sonmezer_molecular_2021,
	title = {Molecular Co-occupancy Identifies Transcription Factor Binding Cooperativity In Vivo},
	year = {2021},
	volume = {81},
	issn = {1097-4164},
	doi = {10.1016/j.molcel.2020.11.015},
	abstract = {Gene activation requires the cooperative activity of multiple transcription factors at cis-regulatory elements ({CREs}). Yet, most transcription factors have short residence time, questioning the requirement of their physical co-occupancy on {DNA} to achieve cooperativity. Here, we present a {DNA} footprinting method that detects individual molecular interactions of transcription factors and nucleosomes with {DNA} in vivo. We apply this strategy to quantify the simultaneous binding of multiple transcription factors on single {DNA} molecules at mouse {CREs}. Analysis of the binary occupancy patterns at thousands of motif combinations reveals that high {DNA} co-occupancy occurs for most types of transcription factors, in the absence of direct physical interaction, at sites of competition with nucleosomes. Perturbation of pairwise interactions demonstrates the function of molecular co-occupancy in binding cooperativity. Our results reveal the interactions regulating {CREs} at molecular resolution and identify {DNA} co-occupancy as a widespread cooperativity mechanism used by transcription factors to remodel chromatin.},
	pages = {255--267.e6},
	number = {2},
	journal = {Molecular Cell},
	shortjournal = {Mol Cell},
	author = {Sönmezer, Can and Kleinendorst, Rozemarijn and Imanci, Dilek and Barzaghi, Guido and Villacorta, Laura and Schübeler, Dirk and Benes, Vladimir and Molina, Nacho and Krebs, Arnaud Regis},
	date = {2021-01-21},
	pmid = {33290745},
}

@article{kleinendorst_genome-wide_2021,
	title = {Genome-wide quantification of transcription factor binding at single-{DNA}-molecule resolution using methyl-transferase footprinting},
	year = {2021},
	issn = {1754-2189, 1750-2799},
	url = {https://www.nature.com/articles/s41596-021-00630-1},
	doi = {10.1038/s41596-021-00630-1},
	journal = {Nature Protocols},
	shortjournal = {Nat Protoc},
	author = {Kleinendorst, Rozemarijn W. D. and Barzaghi, Guido and Smith, Mike L. and Zaugg, Judith B. and Krebs, Arnaud R.},
	urldate = {2021-11-25},
	date = {2021-11-12},
	langid = {english},
}

@article{krebs_genome-wide_2017,
	title = {Genome-wide Single-Molecule Footprinting Reveals High {RNA} Polymerase {II} Turnover at Paused Promoters},
	year = {2017},
	volume = {67},
	issn = {1097-4164},
	doi = {10.1016/j.molcel.2017.06.027},
	abstract = {Transcription initiation entails chromatin opening followed by pre-initiation complex formation and {RNA} polymerase {II} recruitment. Subsequent polymerase elongation requires additional signals, resulting in increased residence time downstream of the start site, a phenomenon referred to as pausing. Here, we harnessed single-molecule footprinting to quantify distinct steps of initiation in vivo throughout the Drosophila genome. This identifies the impact of promoter structure on initiation dynamics in relation to nucleosomal occupancy. Additionally, perturbation of transcriptional initiation reveals an unexpectedly high turnover of polymerases at paused promoters-an observation confirmed at the level of nascent {RNAs}. These observations argue that absence of elongation is largely caused by premature termination rather than by stable polymerase stalling. In support of this non-processive model, we observe that induction of the paused heat shock promoter depends on continuous initiation. Our study provides a framework to quantify protein binding at single-molecule resolution and refines concepts of transcriptional pausing.},
	pages = {411--422.e4},
	number = {3},
	journal = {Molecular Cell},
	shortjournal = {Mol Cell},
	author = {Krebs, Arnaud R. and Imanci, Dilek and Hoerner, Leslie and Gaidatzis, Dimos and Burger, Lukas and Schübeler, Dirk},
	date = {2017-08-03},
	pmid = {28735898},
	pmcid = {PMC5548954},
}

@article{kelly_genome-wide_2012,
	title = {Genome-wide mapping of nucleosome positioning and {DNA} methylation within individual {DNA} molecules.},
	year = {2012},
	volume = {22},
	issn = {1549-5469},
	doi = {10.1101/gr.143008.112},
	abstract = {{DNA} methylation and nucleosome positioning work together to generate chromatin structures that regulate gene expression. Nucleosomes are typically mapped using nuclease digestion requiring significant amounts of material and varying enzyme concentrations. We have developed a method ({NOMe}-seq) that uses a {GpC} methyltransferase (M.{CviPI}) and next generation sequencing to generate a high resolution footprint of nucleosome positioning genome-wide using less than 1 million cells while retaining endogenous {DNA} methylation information from the same {DNA} strand. Using a novel bioinformatics pipeline, we show a striking anti-correlation between nucleosome occupancy and {DNA} methylation at {CTCF} regions that is not present at promoters. We further show that the extent of nucleosome depletion at promoters is directly correlated to expression level and can accommodate multiple nucleosomes and provide genome-wide evidence that expressed non-{CpG} island promoters are nucleosome-depleted. Importantly, {NOMe}-seq obtains {DNA} methylation and nucleosome positioning information from the same {DNA} molecule, giving the first genome-wide {DNA} methylation and nucleosome positioning correlation at the single molecule, and thus, single cell level, that can be used to monitor disease progression and response to therapy.},
	pages = {2497--2506},
	number = {12},
	journal = {Genome research},
	author = {Kelly, Theresa K and Liu, Yaping and Lay, Fides D and Liang, Gangning and Berman, Benjamin P and Jones, Peter A},
	date = {2012},
	pmid = {22960375},
	note = {{ISBN}: 1549-5469 (Electronic)\${\textbackslash}backslash\$n1088-9051 (Linking)},
}

@article{nabilsi_multiplex_2014,
	title = {Multiplex mapping of chromatin accessibility and {DNA} methylation within targeted single molecules identifies epigenetic heterogeneity in neural stem cells and glioblastoma},
	year = {2014},
	volume = {24},
	issn = {10889051},
	doi = {10.1101/gr.161737.113},
	abstract = {Human tumors are comprised of heterogeneous cell populations that display diverse molecular and phenotypic features. To examine the extent to which epigenetic differences contribute to intratumoral cellular heterogeneity, we have developed a high-throughput method, termed {MAPit}-patch. The method uses multiplexed amplification of targeted sequences from submicrogram quantities of genomic {DNA} followed by next generation bisulfite sequencing. This provides highly scalable and simultaneous mapping of chromatin accessibility and {DNA} methylation on single molecules at high resolution. Long sequencing reads from targeted regions maintain the structural integrity of epigenetic information and provide substantial depth of coverage, detecting for the first time minority subpopulations of epigenetic configurations formerly obscured by existing genome-wide and population-ensemble methodologies. Analyzing a cohort of 71 promoters of genes with exons commonly mutated in cancer, {MAPit}-patch uncovered several differentially accessible and methylated promoters that are associated with altered gene expression between neural stem cell ({NSC}) and glioblastoma ({GBM}) cell populations. In addition, considering each promoter individually, substantial epigenetic heterogeneity was observed across the sequenced molecules, indicating the presence of epigenetically distinct cellular subpopulations. At the divergent {MLH}1/{EPM}2AIP1 promoter, a locus with three well-defined, nucleosome-depleted regions ({NDRs}), a fraction of promoter copies with inaccessible chromatin was detected and enriched upon selection of temozolomide-tolerant {GBM} cells. These results illustrate the biological relevance of epigenetically distinct subpopulations that in part underlie the phenotypic heterogeneity of tumor cell populations. Furthermore, these findings show that alterations in chromatin accessibility without accompanying changes in {DNA} methylation may constitute a novel class of epigenetic biomarker.},
	pages = {329--339},
	number = {2},
	journal = {Genome Research},
	author = {Nabilsi, Nancy H. and Deleyrolle, Loic P. and Darst, Russell P. and Riva, Alberto and Reynolds, Brent A. and Kladde, Michael P.},
	date = {2014},
	pmid = {24105770},
	note = {{ISBN}: 1549-5469 (Electronic)\${\textbackslash}backslash\$r1088-9051 (Linking)},
}

@article{ferguson-smith_discovery_2018,
	title = {The discovery and importance of genomic imprinting},
	year = {2018},
	volume = {7},
	issn = {2050-084X},
	doi = {10.7554/eLife.42368},
	abstract = {The discovery of genomic imprinting by Davor Solter, Azim Surani and co-workers in
the mid-1980s has provided a foundation for the study of epigenetic inheritance and
the epigenetic control of gene activity and repression, especially during
development. It also has shed light on a range of diseases, including both rare
genetic disorders and common diseases. This article is being published to celebrate
Solter and Surani receiving a 2018 Canada Gairdner International Award "for the
discovery of mammalian genomic imprinting that causes parent-of-origin specific gene
expression and its consequences for development and disease".},
	pages = {e42368},
	journal = {{eLife}},
	shortjournal = {{eLife}},
	author = {Ferguson-Smith, Anne C and Bourc'his, Deborah},
	pmid = {30343680},
	pmcid = {PMC6197852},
}

@article{feldmann_transcription_2013,
	title = {Transcription Factor Occupancy Can Mediate Active Turnover of {DNA} Methylation at Regulatory Regions},
	year = {2013},
	volume = {9},
	issn = {1553-7404},
	doi = {10.1371/journal.pgen.1003994},
	abstract = {Distal regulatory elements, including enhancers, play a critical role in regulating gene activity. Transcription factor binding to these elements correlates with Low Methylated Regions ({LMRs}) in a process that is poorly understood. Here we ask whether and how actual occupancy of {DNA}-binding factors is linked to {DNA} methylation at the level of individual molecules. Using {CTCF} as an example, we observe that frequency of binding correlates with the likelihood of a demethylated state and sites of low occupancy display heterogeneous {DNA} methylation within the {CTCF} motif. In line with a dynamic model of binding and {DNA} methylation turnover, we find that 5-hydroxymethylcytosine (5hmC), formed as an intermediate state of active demethylation, is enriched at {LMRs} in stem and somatic cells. Moreover, a significant fraction of changes in 5hmC during differentiation occurs at these regions, suggesting that transcription factor activity could be a key driver for active demethylation. Since deletion of {CTCF} is lethal for embryonic stem cells, we used genetic deletion of {REST} as another {DNA}-binding factor implicated in {LMR} formation to test this hypothesis. The absence of {REST} leads to a decrease of hydroxymethylation and a concomitant increase of {DNA} methylation at its binding sites. These data support a model where {DNA}-binding factors can mediate turnover of {DNA} methylation as an integral part of maintenance and reprogramming of regulatory regions.},
	pages = {e1003994},
	number = {12},
	journal = {{PLOS} Genetics},
	shortjournal = {{PLOS} Genetics},
	author = {Feldmann, Angelika and Ivanek, Robert and Murr, Rabih and Gaidatzis, Dimos and Burger, Lukas and Schübeler, Dirk},
	date = {2013-12-19},
	langid = {english},
	note = {publisher: Public Library of Science},
}

@article{wang_widespread_2012,
	title = {Widespread plasticity in {CTCF} occupancy linked to {DNA} methylation},
	year = {2012},
	volume = {22},
	issn = {1088-9051},
	doi = {10.1101/gr.136101.111},
	abstract = {{CTCF} is a ubiquitously expressed regulator of fundamental genomic processes including transcription, intra- and interchromosomal interactions, and chromatin structure. Because of its critical role in genome function, {CTCF} binding patterns have long been assumed to be largely invariant across different cellular environments. Here we analyze genome-wide occupancy patterns of {CTCF} by {ChIP}-seq in 19 diverse human cell types, including normal primary cells and immortal lines. We observed highly reproducible yet surprisingly plastic genomic binding landscapes, indicative of strong cell-selective regulation of {CTCF} occupancy. Comparison with massively parallel bisulfite sequencing data indicates that 41\% of variable {CTCF} binding is linked to differential {DNA} methylation, concentrated at two critical positions within the {CTCF} recognition sequence. Unexpectedly, {CTCF} binding patterns were markedly different in normal versus immortal cells, with the latter showing widespread disruption of {CTCF} binding associated with increased methylation. Strikingly, this disruption is accompanied by up-regulation of {CTCF} expression, with the result that both normal and immortal cells maintain the same average number of {CTCF} occupancy sites genome-wide. These results reveal a tight linkage between {DNA} methylation and the global occupancy patterns of a major sequence-specific regulatory factor.},
	pages = {1680--1688},
	number = {9},
	journal = {Genome Research},
	shortjournal = {Genome Res},
	author = {Wang, Hao and Maurano, Matthew T. and Qu, Hongzhu and Varley, Katherine E. and Gertz, Jason and Pauli, Florencia and Lee, Kristen and Canfield, Theresa and Weaver, Molly and Sandstrom, Richard and Thurman, Robert E. and Kaul, Rajinder and Myers, Richard M. and Stamatoyannopoulos, John A.},
	date = {2012-09},
	pmid = {22955980},
	pmcid = {PMC3431485},
}

@article{hashimoto_structural_2017,
	title = {Structural Basis for the Versatile and Methylation-Dependent Binding of {CTCF} to {DNA}},
	year = {2017},
	volume = {66},
	issn = {1097-2765},
	doi = {10.1016/j.molcel.2017.05.004},
	pages = {711--720.e3},
	number = {5},
	journal = {Molecular Cell},
	shortjournal = {Molecular Cell},
	author = {Hashimoto, Hideharu and Wang, Dongxue and Horton, John R. and Zhang, Xing and Corces, Victor G. and Cheng, Xiaodong},
	date = {2017-06-01},
	pmid = {28529057},
	note = {publisher: Elsevier},
}

@article{donaghey_genetic_2018,
	title = {Genetic determinants and epigenetic effects of pioneer-factor occupancy},
	year = {2018},
	volume = {50},
	issn = {1061-4036, 1546-1718},
	doi = {10.1038/s41588-017-0034-3},
	pages = {250--258},
	number = {2},
	journal = {Nature Genetics},
	shortjournal = {Nat Genet},
	author = {Donaghey, Julie and Thakurela, Sudhir and Charlton, Jocelyn and Chen, Jennifer S. and Smith, Zachary D. and Gu, Hongcang and Pop, Ramona and Clement, Kendell and Stamenova, Elena K. and Karnik, Rahul and Kelley, David R. and Gifford, Casey A. and Cacchiarelli, Davide and Rinn, John L. and Gnirke, Andreas and Ziller, Michael J. and Meissner, Alexander},
	date = {2018-02},
	langid = {english},
}

@article{charlet_bivalent_2016,
	title = {Bivalent Regions of Cytosine Methylation and H3K27 Acetylation Suggest an Active Role for {DNA} Methylation at Enhancers},
	year = {2016},
	volume = {62},
	issn = {10972765},
	doi = {10.1016/j.molcel.2016.03.033},
	abstract = {The role of cytosine methylation in the structure and function of enhancers is not well understood. In this study, we investigate the role of {DNA} methylation at enhancers by comparing the epigenomes of the {HCT}116 cell line and its highly demethylated derivative, {DKO}1. Unlike promoters, a portion of regular and super- or stretch enhancers show active H3K27ac marks co-existing with extensive {DNA} methylation, demonstrating the unexpected presence of bivalent chromatin in both cultured and uncultured cells. Furthermore, our ﬁndings also show that bivalent regions have fewer nucleosomedepleted regions and transcription factor-binding sites than monovalent regions. Reduction of {DNA} methylation genetically or pharmacologically leads to a decrease of the H3K27ac mark. Thus, {DNA} methylation plays an unexpected dual role at enhancer regions, being anti-correlated focally at transcription factor-binding sites but positively correlated globally with the active H3K27ac mark to ensure structural enhancer integrity.},
	pages = {422--431},
	number = {3},
	journal = {Molecular Cell},
	shortjournal = {Molecular Cell},
	author = {Charlet, Jessica and Duymich, Christopher E. and Lay, Fides D. and Mundbjerg, Kamilla and Dalsgaard Sørensen, Karina and Liang, Gangning and Jones, Peter A.},
	date = {2016-05},
	langid = {english},
}

@article{nie_c-myc_2012,
	title = {c-Myc Is a Universal Amplifier of Expressed Genes in Lymphocytes and Embryonic Stem Cells},
	year = {2012},
	volume = {151},
	issn = {0092-8674, 1097-4172},
	doi = {10.1016/j.cell.2012.08.033},
	pages = {68--79},
	number = {1},
	journal = {Cell},
	shortjournal = {Cell},
	author = {Nie, Zuqin and Hu, Gangqing and Wei, Gang and Cui, Kairong and Yamane, Arito and Resch, Wolfgang and Wang, Ruoning and Green, Douglas R. and Tessarollo, Lino and Casellas, Rafael and Zhao, Keji and Levens, David},
	date = {2012-09-28},
	pmid = {23021216},
	note = {publisher: Elsevier},
}

@article{lin_transcriptional_2012,
	title = {Transcriptional Amplification in Tumor Cells with Elevated c-Myc},
	year = {2012},
	volume = {151},
	issn = {0092-8674, 1097-4172},
	doi = {10.1016/j.cell.2012.08.026},
	pages = {56--67},
	number = {1},
	journal = {Cell},
	shortjournal = {Cell},
	author = {Lin, Charles Y. and Lovén, Jakob and Rahl, Peter B. and Paranal, Ronald M. and Burge, Christopher B. and Bradner, James E. and Lee, Tong Ihn and Young, Richard A.},
	date = {2012-09-28},
	pmid = {23021215},
	note = {publisher: Elsevier},
}

@article{kress_myc_2015,
	title = {{MYC}: connecting selective transcriptional control to global {RNA} production},
	year = {2015},
	volume = {15},
	issn = {1474-1768},
	doi = {10.1038/nrc3984},
	shorttitle = {{MYC}},
	abstract = {{MYC} is a key integrator of growth-regulatory and oncogenic signalling pathways.{MYC} binds open chromatin and functions as a transcriptional activator and repressor of selected gene subsets.Detection of {MYC} binding to chromatin should not be systematically equated to a productive transcriptional event.{MYC} regulates genes and pathways that can feed back on global {RNA} production.Various {MYC} target gene products are crucial for tumour maintenance and may be targeted therapeutically: identifying these key effectors is a main priority in the field.},
	pages = {593--607},
	number = {10},
	journal = {Nature Reviews Cancer},
	shortjournal = {Nat Rev Cancer},
	author = {Kress, Theresia R. and Sabò, Arianna and Amati, Bruno},
	date = {2015-10},
	langid = {english},
	note = {number: 10
publisher: Nature Publishing Group},
}

@article{rulands_genome-scale_2018,
	title = {Genome-Scale Oscillations in {DNA} Methylation during Exit from Pluripotency},
	year = {2018},
	volume = {7},
	issn = {2405-4712},
	doi = {10.1016/j.cels.2018.06.012},
	abstract = {Pluripotency is accompanied by the erasure of parental epigenetic memory, with naïve pluripotent cells exhibiting global {DNA} hypomethylation both in vitro and in vivo. Exit from pluripotency and priming for differentiation into somatic lineages is associated with genome-wide de novo {DNA} methylation. We show that during this phase, co-expression of enzymes required for {DNA} methylation turnover, {DNMT}3s and {TETs}, promotes cell-to-cell variability in this epigenetic mark. Using a combination of single-cell sequencing and quantitative biophysical modeling, we show that this variability is associated with coherent, genome-scale oscillations in {DNA} methylation with an amplitude dependent on {CpG} density. Analysis of parallel single-cell transcriptional and epigenetic profiling provides evidence for oscillatory dynamics both in vitro and in vivo. These observations provide insights into the emergence of epigenetic heterogeneity during early embryo development, indicating that dynamic changes in {DNA} methylation might influence early cell fate decisions.},
	pages = {63--76.e12},
	number = {1},
	journal = {Cell Systems},
	shortjournal = {Cell Syst},
	author = {Rulands, Steffen and Lee, Heather J. and Clark, Stephen J. and Angermueller, Christof and Smallwood, Sébastien A. and Krueger, Felix and Mohammed, Hisham and Dean, Wendy and Nichols, Jennifer and Rugg-Gunn, Peter and Kelsey, Gavin and Stegle, Oliver and Simons, Benjamin D. and Reik, Wolf},
	date = {2018-07-25},
	pmid = {30031774},
	pmcid = {PMC6066359},
}

@article{baylin_decade_2011,
	title = {A decade of exploring the cancer epigenome — biological and translational implications},
	year = {2011},
	volume = {11},
	issn = {1474-175X},
	doi = {10.1038/nrc3130},
	abstract = {The past decade has highlighted the central role of epigenetic processes in cancer causation, progression and treatment. Next-generation sequencing is providing a window for visualizing the human epigenome and how it is altered in cancer. This view provides many surprises, including linking epigenetic abnormalities to mutations in genes that control {DNA} methylation, the packaging and the function of {DNA} in chromatin, and metabolism. Epigenetic alterations are leading candidates for the development of specific markers for cancer detection, diagnosis and prognosis. The enzymatic processes that control the epigenome present new opportunities for deriving therapeutic strategies designed to reverse transcriptional abnormalities that are inherent to the cancer epigenome.},
	pages = {726--734},
	number = {10},
	journal = {Nature Reviews. Cancer},
	shortjournal = {Nat Rev Cancer},
	author = {Baylin, Stephen B. and Jones, Peter A.},
	date = {2011-09-23},
	pmid = {21941284},
	pmcid = {PMC3307543},
}

@article{pintacuda_hnrnpk_2017,
	title = {{hnRNPK} Recruits {PCGF}3/5-{PRC}1 to the Xist {RNA} B-Repeat to Establish Polycomb-Mediated Chromosomal Silencing.},
	year = {2017},
	volume = {68},
	rights = {Copyright © 2017 The Authors. Published by Elsevier Inc. All rights reserved.},
	issn = {1097-4164 1097-2765},
	doi = {10.1016/j.molcel.2017.11.013},
	abstract = {The Polycomb-repressive complexes {PRC}1 and {PRC}2 play a key role in chromosome silencing induced by the non-coding {RNA} Xist. Polycomb recruitment is initiated by  the {PCGF}3/5-{PRC}1 complex, which catalyzes chromosome-wide H2A lysine 119  ubiquitylation, signaling recruitment of other {PRC}1 complexes, and {PRC}2. However,  the molecular mechanism for {PCGF}3/5-{PRC}1 recruitment by Xist {RNA} is not understood.  Here we define the Xist {RNA} Polycomb Interaction Domain ({XR}-{PID}), a 600 nt sequence  encompassing the Xist B-repeat element. Deletion of {XR}-{PID} abolishes Xist-dependent  Polycomb recruitment, in turn abrogating Xist-mediated gene silencing and reversing  Xist-induced chromatin inaccessibility. We identify the {RNA}-binding protein {hnRNPK}  as the principal {XR}-{PID} binding factor required to recruit {PCGF}3/5-{PRC}1.  Accordingly, synthetically tethering {hnRNPK} to Xist {RNA} lacking {XR}-{PID} is sufficient  for Xist-dependent Polycomb recruitment. Our findings define a key pathway for  Polycomb recruitment by Xist {RNA}, providing important insights into mechanisms of  chromatin modification by non-coding {RNA}.},
	pages = {955--969.e10},
	number = {5},
	journal = {Molecular cell},
	shortjournal = {Mol Cell},
	author = {Pintacuda, Greta and Wei, Guifeng and Roustan, Chloë and Kirmizitas, Burcu Anil and Solcan, Nicolae and Cerase, Andrea and Castello, Alfredo and Mohammed, Shabaz and Moindrot, Benoît and Nesterova, Tatyana B. and Brockdorff, Neil},
	date = {2017-12-07},
	pmid = {29220657},
	pmcid = {PMC5735038},
	keywords = {*X Chromosome Inactivation, Animals, Binding Sites, Cell Line, Embryonic Stem Cells/*metabolism, Heterogeneous-Nuclear Ribonucleoprotein K, Histones/metabolism, Lysine/metabolism, Mice, Polycomb Repressive Complex 1/genetics/*metabolism, Polycomb-Group Proteins/genetics/*metabolism, Protein Binding, Protein Interaction Domains and Motifs, Ribonucleoproteins/genetics/*metabolism, {RNA} Interference, {RNA}, Long Noncoding/genetics/*metabolism, Transcription, Genetic, Transfection, Ubiquitination, X Chromosome/genetics/*metabolism},
}

@article{gaidatzis_quasr_2015,
	title = {{QuasR}: quantification and annotation of short reads in R},
	year = {2015},
	volume = {31},
	issn = {1367-4803},
	url = {https://doi.org/10.1093/bioinformatics/btu781},
	doi = {10.1093/bioinformatics/btu781},
	abstract = {Summary: {QuasR} is a package for the integrated analysis of high-throughput sequencing data in R, covering all steps from read preprocessing, alignment and quality control to quantification. {QuasR} supports different experiment types (including {RNA}-seq, {ChIP}-seq and Bis-seq) and analysis variants (e.g. paired-end, stranded, spliced and allele-specific), and is integrated in Bioconductor so that its output can be directly processed for statistical analysis and visualization.Availability and implementation: {QuasR} is implemented in R and C/C++. Source code and binaries for major platforms (Linux, {OS} X and {MS} Windows) are available from Bioconductor (www.bioconductor.org/packages/release/bioc/html/{QuasR}.html). The package includes a ‘vignette’ with step-by-step examples for typical work ﬂows.Contact:michael.stadler@fmi.{chSupplementary} information:Supplementary data are available at Bioinformatics online.},
	pages = {1130--1132},
	number = {7},
	journal = {Bioinformatics},
	shortjournal = {Bioinformatics},
	author = {Gaidatzis, Dimos and Lerch, Anita and Hahne, Florian and Stadler, Michael B.},
	urldate = {2022-02-23},
	date = {2015-04-01},
}

@article{khan_jaspar_2018,
	title = {{JASPAR} 2018: update of the open-access database of transcription factor binding profiles and its web framework},
	year = {2018},
	volume = {46},
	issn = {0305-1048},
	url = {https://doi.org/10.1093/nar/gkx1126},
	doi = {10.1093/nar/gkx1126},
	abstract = {{JASPAR} (http://jaspar.genereg.net) is an open-access database of curated, non-redundant transcription factor ({TF})-binding profiles stored as position frequency matrices ({PFMs}) and {TF} flexible models ({TFFMs}) for {TFs} across multiple species in six taxonomic groups. In the 2018 release of {JASPAR}, the {CORE} collection has been expanded with 322 new {PFMs} (60 for vertebrates and 262 for plants) and 33 {PFMs} were updated (24 for vertebrates, 8 for plants and 1 for insects). These new profiles represent a 30\% expansion compared to the 2016 release. In addition, we have introduced 316 {TFFMs} (95 for vertebrates, 218 for plants and 3 for insects). This release incorporates clusters of similar {PFMs} in each taxon and each {TF} class per taxon. The {JASPAR} 2018 {CORE} vertebrate collection of {PFMs} was used to predict {TF}-binding sites in the human genome. The predictions are made available to the scientific community through a {UCSC} Genome Browser track data hub. Finally, this update comes with a new web framework with an interactive and responsive user-interface, along with new features. All the underlying data can be retrieved programmatically using a {RESTful} {API} and through the {JASPAR} 2018 R/Bioconductor package.},
	pages = {D260--D266},
	issue = {D1},
	journal = {Nucleic Acids Research},
	shortjournal = {Nucleic Acids Research},
	author = {Khan, Aziz and Fornes, Oriol and Stigliani, Arnaud and Gheorghe, Marius and Castro-Mondragon, Jaime A and van der Lee, Robin and Bessy, Adrien and Chèneby, Jeanne and Kulkarni, Shubhada R and Tan, Ge and Baranasic, Damir and Arenillas, David J and Sandelin, Albin and Vandepoele, Klaas and Lenhard, Boris and Ballester, Benoît and Wasserman, Wyeth W and Parcy, François and Mathelier, Anthony},
	urldate = {2022-02-23},
	date = {2018-01-04},
}

@article{moore_curated_2020,
	title = {A curated benchmark of enhancer-gene interactions for evaluating enhancer-target gene prediction methods},
	year = {2020},
	volume = {21},
	issn = {1474-760X},
	url = {https://doi.org/10.1186/s13059-019-1924-8},
	doi = {10.1186/s13059-019-1924-8},
	abstract = {Many genome-wide collections of candidate cis-regulatory elements ({cCREs}) have been defined using genomic and epigenomic data, but it remains a major challenge to connect these elements to their target genes.},
	pages = {17},
	number = {1},
	journal = {Genome Biology},
	shortjournal = {Genome Biology},
	author = {Moore, Jill E. and Pratt, Henry E. and Purcaro, Michael J. and Weng, Zhiping},
	date = {2020-01-22},
}

@misc{kuhn_corrr_2022,
	title = {corrr: Correlations in R},
	year = {2022},
	howpublished = {\url{https://github.com/tidymodels/corrr}, \url{https://corrr.tidymodels.org}},
	author = {Kuhn, M and Jackson, S and Cimentada, J},
	date = {2022},
}

@book{wickham_ggplot2_2016,
	title = {ggplot2: Elegant Graphics for Data Analysis},
	year = {2016},
	isbn = {978-3-319-24277-4},
	url = {https://ggplot2.tidyverse.org},
	publisher = {ggplot2: Elegant Graphics for Data Analysis},
	author = {Wickham, Hadley},
	date = {2016},
}

@misc{gu_rgreat_2021,
	title = {{rGREAT}: Client for {GREAT} Analysis},
	year = {2021},
	howpublished = {\url{https://github.com/jokergoo/rGREAT}, \url{http://great.stanford.edu/public/html/}},
	author = {Gu, Zuguang},
	date = {2021},
}


@article{mclean_great_2010,
	title = {{GREAT} improves functional interpretation of cis-regulatory regions.},
	year = {2010},
	volume = {28},
	issn = {1546-1696 1087-0156},
	doi = {10.1038/nbt.1630},
	abstract = {We developed the Genomic Regions Enrichment of Annotations Tool ({GREAT}) to analyze the functional significance of cis-regulatory regions identified by localized  measurements of {DNA} binding events across an entire genome. Whereas previous methods  took into account only binding proximal to genes, {GREAT} is able to properly  incorporate distal binding sites and control for false positives using a binomial  test over the input genomic regions. {GREAT} incorporates annotations from 20  ontologies and is available as a web application. Applying {GREAT} to data sets from  chromatin immunoprecipitation coupled with massively parallel sequencing ({ChIP}-seq)  of multiple transcription-associated factors, including {SRF}, {NRSF}, {GABP}, Stat3 and  p300 in different developmental contexts, we recover many functions of these factors  that are missed by existing gene-based tools, and we generate testable hypotheses.  The utility of {GREAT} is not limited to {ChIP}-seq, as it could also be applied to open  chromatin, localized epigenomic markers and similar functional data sets, as well as  comparative genomics sets.},
	pages = {495--501},
	number = {5},
	journal = {Nature biotechnology},
	shortjournal = {Nat Biotechnol},
	author = {{McLean}, Cory Y. and Bristor, Dave and Hiller, Michael and Clarke, Shoa L. and Schaar, Bruce T. and Lowe, Craig B. and Wenger, Aaron M. and Bejerano, Gill},
	date = {2010-05},
	pmid = {20436461},
	pmcid = {PMC4840234},
	keywords = {Animals, Mice, Protein Binding, Humans, *Genome, *Regulatory Elements, Transcriptional, *Software, Chromatin Immunoprecipitation, Data Mining/*methods, Databases, Genetic, E1A-Associated p300 Protein, Genomics/*methods, Jurkat Cells, Serum Response Factor},
}

@article{xie_base-resolution_2012,
	title = {Base-Resolution Analyses of Sequence and Parent-of-Origin Dependent {DNA} Methylation in the Mouse Genome},
	year = {2012},
	volume = {148},
	issn = {0092-8674},
	url = {https://www.sciencedirect.com/science/article/pii/S0092867412001407},
	doi = {10.1016/j.cell.2011.12.035},
	abstract = {Summary
Differential methylation of the two parental genomes in placental mammals is essential for genomic imprinting and embryogenesis. To systematically study this epigenetic process, we have generated a base-resolution, allele-specific {DNA} methylation ({ASM}) map in the mouse genome. We find parent-of-origin dependent (imprinted) {ASM} at 1,952 {CG} dinucleotides. These imprinted {CGs} form 55 discrete clusters including virtually all known germline differentially methylated regions ({DMRs}) and 23 previously unknown {DMRs}, with some occurring at {microRNA} genes. We also identify sequence-dependent {ASM} at 131,765 {CGs}. Interestingly, methylation at these sites exhibits a strong dependence on the immediate adjacent bases, allowing us to define a conserved sequence preference for the mammalian {DNA} methylation machinery. Finally, we report a surprising presence of non-{CG} methylation in the adult mouse brain, with some showing evidence of imprinting. Our results provide a resource for understanding the mechanisms of imprinting and allele-specific gene expression in mammalian cells.},
	pages = {816--831},
	number = {4},
	journal = {Cell},
	shortjournal = {Cell},
	author = {Xie, Wei and Barr, Cathy L. and Kim, Audrey and Yue, Feng and Lee, Ah Young and Eubanks, James and Dempster, Emma L. and Ren, Bing},
	date = {2012-02-17},
}

@article{bolger_trimmomatic_2014,
	title = {Trimmomatic: a flexible trimmer for Illumina sequence data.},
	year = {2014},
	volume = {30},
	rights = {© The Author 2014. Published by Oxford University Press.},
	issn = {1367-4811 1367-4803},
	doi = {10.1093/bioinformatics/btu170},
	abstract = {{MOTIVATION}: Although many next-generation sequencing ({NGS}) read preprocessing tools already existed, we could not find any tool or combination of tools that met our  requirements in terms of flexibility, correct handling of paired-end data and high  performance. We have developed Trimmomatic as a more flexible and efficient  preprocessing tool, which could correctly handle paired-end data. {RESULTS}: The value  of {NGS} read preprocessing is demonstrated for both reference-based and  reference-free tasks. Trimmomatic is shown to produce output that is at least  competitive with, and in many cases superior to, that produced by other tools, in  all scenarios tested. {AVAILABILITY} {AND} {IMPLEMENTATION}: Trimmomatic is licensed under  {GPL} V3. It is cross-platform (Java 1.5+ required) and available at  http://www.usadellab.org/cms/index.php?page=trimmomatic {CONTACT}:  {SUPPLEMENTARY} {INFORMATION}: Supplementary data are  available at Bioinformatics online.},
	pages = {2114--2120},
	number = {15},
	journal = {Bioinformatics (Oxford, England)},
	shortjournal = {Bioinformatics},
	author = {Bolger, Anthony M. and Lohse, Marc and Usadel, Bjoern},
	date = {2014-08-01},
	pmid = {24695404},
	pmcid = {PMC4103590},
	keywords = {*Software, Databases, Genetic, Computational Biology, High-Throughput Nucleotide Sequencing/*methods},
}

@misc{Picard2019toolkit,
  title = {Picard toolkit},
  year = {2019},
  publisher = {Broad Institute},
  journal = {Broad Institute, GitHub repository},
  howpublished = {\url{https://broadinstitute.github.io/picard/}}
}

@article{liu_catalytic-independent_2017,
	title = {Catalytic-Independent Functions of {PARP}-1 Determine Sox2 Pioneer Activity at Intractable Genomic Loci.},
	year = {2017},
	volume = {65},
	rights = {Copyright © 2017 Elsevier Inc. All rights reserved.},
	issn = {1097-4164 1097-2765},
	doi = {10.1016/j.molcel.2017.01.017},
	abstract = {Pioneer transcription factors ({TFs}) function as genomic first responders, binding to inaccessible regions of chromatin to promote enhancer formation. The mechanism by  which pioneer {TFs} gain access to chromatin remains an important unanswered question.  Here we show that {PARP}-1, a nucleosome-binding protein, cooperates with intrinsic  properties of the pioneer {TF} Sox2 to facilitate its binding to intractable genomic  loci in embryonic stem cells. These actions of {PARP}-1 occur independently of its  poly({ADP}-ribosyl) transferase activity. {PARP}-1-dependent Sox2-binding sites reside  in euchromatic regions of the genome with relatively high nucleosome occupancy and  low co-occupancy by other transcription factors. {PARP}-1 stabilizes Sox2 binding to  nucleosomes at suboptimal sites through cooperative interactions on {DNA}. Our results  define intrinsic and extrinsic features that determine Sox2 pioneer activity. The  conditional pioneer activity observed with Sox2 at a subset of binding sites may be  a key feature of other pioneer {TFs} operating at intractable genomic loci.},
	pages = {589--603.e9},
	number = {4},
	journal = {Molecular cell},
	shortjournal = {Mol Cell},
	author = {Liu, Ziying and Kraus, W. Lee},
	date = {2017-02-16},
	pmid = {28212747},
	pmcid = {PMC5319724},
	keywords = {Animals, Binding Sites, Cell Line, Mice, Protein Binding, Transfection, Humans, *Gene Expression Regulation, Developmental, *Genetic Loci, chromatin, {DNA}/genetics/*metabolism, embryonic stem cells, Embryonic Stem Cells/*enzymology, Euchromatin/*enzymology/genetics, nucleosome, nucleosome rotational positioning, Nucleosomes/*enzymology/genetics, {PARP}-1, pioneer transcription factor, Pluripotent Stem Cells/*enzymology, Poly ({ADP}-Ribose) Polymerase-1/genetics/*metabolism, Signal Transduction, Sox2, {SOXB}1 Transcription Factors/genetics/*metabolism, Time Factors, transcription},
}

@article{shen_map_2012,
	title = {A map of the cis-regulatory sequences in the mouse genome.},
	year = {2012},
	volume = {488},
	issn = {1476-4687 0028-0836},
	doi = {10.1038/nature11243},
	abstract = {The laboratory mouse is the most widely used mammalian model organism in biomedical research. The 2.6 × 10(9) bases of the mouse genome possess a high degree of  conservation with the human genome, so a thorough annotation of the mouse genome  will be of significant value to understanding the function of the human genome. So  far, most of the functional sequences in the mouse genome have yet to be found, and  the cis-regulatory sequences in particular are still poorly annotated. Comparative  genomics has been a powerful tool for the discovery of these sequences, but on its  own it cannot resolve their temporal and spatial functions. Recently, {ChIP}-Seq has  been developed to identify cis-regulatory elements in the genomes of several  organisms including humans, Drosophila melanogaster and Caenorhabditis elegans. Here  we apply the same experimental approach to a diverse set of 19 tissues and cell  types in the mouse to produce a map of nearly 300,000 murine cis-regulatory  sequences. The annotated sequences add up to 11\% of the mouse genome, and include  more than 70\% of conserved non-coding sequences. We define tissue-specific enhancers  and identify potential transcription factors regulating gene expression in each  tissue or cell type. Finally, we show that much of the mouse genome is organized  into domains of coordinately regulated enhancers and promoters. Our results provide  a resource for the annotation of functional elements in the mammalian genome and for  the study of mechanisms regulating tissue-specific gene expression.},
	pages = {116--120},
	number = {7409},
	journal = {Nature},
	shortjournal = {Nature},
	author = {Shen, Yin and Yue, Feng and {McCleary}, David F. and Ye, Zhen and Edsall, Lee and Kuan, Samantha and Wagner, Ulrich and Dixon, Jesse and Lee, Leonard and Lobanenkov, Victor V. and Ren, Bing},
	date = {2012-08-02},
	pmid = {22763441},
	pmcid = {PMC4041622},
	keywords = {Animals, Sequence Analysis, {DNA}, Chromatin Immunoprecipitation, *Physical Chromosome Mapping, Acetylation, Chromatin/metabolism, Conserved Sequence, Enhancer Elements, Genetic/genetics, Evolution, Molecular, Gene Expression Regulation/*genetics, Genome/*genetics, Male, Methylation, Mice, Inbred C57BL, Mice/*genetics, Molecular Sequence Annotation, Nucleotide Motifs, Organ Specificity, Promoter Regions, Genetic/genetics, Regulatory Sequences, Nucleic Acid/*genetics, Transcription Factors/metabolism},
}

@article{ernst_chromhmm_2012,
	title = {{ChromHMM}: automating chromatin-state discovery and characterization},
	year = {2012},
	volume = {9},
	issn = {1548-7105},
	url = {https://doi.org/10.1038/nmeth.1906},
	doi = {10.1038/nmeth.1906},
	pages = {215--216},
	number = {3},
	journal = {Nature Methods},
	shortjournal = {Nature Methods},
	author = {Ernst, Jason and Kellis, Manolis},
	date = {2012-03-01},
}

@article{gu_enrichedheatmap_2018,
	title = {{EnrichedHeatmap}: an R/Bioconductor package for comprehensive visualization of genomic signal associations},
	year = {2018},
	volume = {19},
	issn = {1471-2164},
	url = {https://doi.org/10.1186/s12864-018-4625-x},
	doi = {10.1186/s12864-018-4625-x},
	abstract = {High-throughput sequencing data are dramatically increasing in volume. Thus, there is urgent need for efficient tools to perform fast and integrative analysis of multiple data types. Enriched heatmap is a specific form of heatmap that visualizes how genomic signals are enriched over specific target regions. It is commonly used and efficient at revealing enrichment patterns especially for high dimensional genomic and epigenomic datasets.},
	pages = {234},
	number = {1},
	journal = {{BMC} Genomics},
	shortjournal = {{BMC} Genomics},
	author = {Gu, Zuguang and Eils, Roland and Schlesner, Matthias and Ishaque, Naveed},
	date = {2018-04-04},
}

@article{langmead_ultrafast_2009,
	title = {Ultrafast and memory-efficient alignment of short {DNA} sequences to the human genome.},
	year = {2009},
	volume = {10},
	issn = {1474-760X 1465-6906 1474-7596},
	doi = {10.1186/gb-2009-10-3-r25},
	abstract = {Bowtie is an ultrafast, memory-efficient alignment program for aligning short {DNA} sequence reads to large genomes. For the human genome, Burrows-Wheeler indexing  allows Bowtie to align more than 25 million reads per {CPU} hour with a memory  footprint of approximately 1.3 gigabytes. Bowtie extends previous Burrows-Wheeler  techniques with a novel quality-aware backtracking algorithm that permits  mismatches. Multiple processor cores can be used simultaneously to achieve even  greater alignment speeds. Bowtie is open source (http://bowtie.cbcb.umd.edu).},
	pages = {R25},
	number = {3},
	journal = {Genome biology},
	shortjournal = {Genome Biol},
	author = {Langmead, Ben and Trapnell, Cole and Pop, Mihai and Salzberg, Steven L.},
	date = {2009},
	pmid = {19261174},
	pmcid = {PMC2690996},
	keywords = {Humans, *Base Sequence, Algorithms, Genome, Human/*genetics, Sequence Alignment/*methods},
}

@article{afgan_galaxy_2018,
	title = {The Galaxy platform for accessible, reproducible and collaborative biomedical analyses: 2018 update},
	year = {2018},
	volume = {46},
	issn = {0305-1048},
	url = {https://doi.org/10.1093/nar/gky379},
	doi = {10.1093/nar/gky379},
	abstract = {Galaxy (homepage: https://galaxyproject.org, main public server: https://usegalaxy.org) is a web-based scientific analysis platform used by tens of thousands of scientists across the world to analyze large biomedical datasets such as those found in genomics, proteomics, metabolomics and imaging. Started in 2005, Galaxy continues to focus on three key challenges of data-driven biomedical science: making analyses accessible to all researchers, ensuring analyses are completely reproducible, and making it simple to communicate analyses so that they can be reused and extended. During the last two years, the Galaxy team and the open-source community around Galaxy have made substantial improvements to Galaxy's core framework, user interface, tools, and training materials. Framework and user interface improvements now enable Galaxy to be used for analyzing tens of thousands of datasets, and \&gt;5500 tools are now available from the Galaxy {ToolShed}. The Galaxy community has led an effort to create numerous high-quality tutorials focused on common types of genomic analyses. The Galaxy developer and user communities continue to grow and be integral to Galaxy's development. The number of Galaxy public servers, developers contributing to the Galaxy framework and its tools, and users of the main Galaxy server have all increased substantially.},
	pages = {W537--W544},
	issue = {W1},
	journal = {Nucleic Acids Research},
	shortjournal = {Nucleic Acids Research},
	author = {Afgan, Enis and Baker, Dannon and Batut, Bérénice and van den Beek, Marius and Bouvier, Dave and Čech, Martin and Chilton, John and Clements, Dave and Coraor, Nate and Grüning, Björn A and Guerler, Aysam and Hillman-Jackson, Jennifer and Hiltemann, Saskia and Jalili, Vahid and Rasche, Helena and Soranzo, Nicola and Goecks, Jeremy and Taylor, James and Nekrutenko, Anton and Blankenberg, Daniel},
	urldate = {2022-03-05},
	date = {2018-07-02},
}

@misc{krueger_felixkruegertrimgalore_2021,
	title = {{FelixKrueger}/{TrimGalore}: v0.6.7},
	year = {2021},
	author = {Krueger, Felix and James, Frankie and Ewels, Phil and Afyounian, Ebrahim and Schuster-Boeckler, Benjamin},
	howpublished = {\url{https://doi.org/10.5281/zenodo.5127899#.YiNnZnY9FYc.mendeley}},
	doi = {10.5281/ZENODO.5127899},
	date = {2021-07},
}

@article{ramirez_deeptools2_2016,
	title = {{deepTools}2: a next generation web server for deep-sequencing data analysis},
	year = {2016},
	volume = {44},
	issn = {0305-1048},
	url = {https://doi.org/10.1093/nar/gkw257},
	doi = {10.1093/nar/gkw257},
	abstract = {We present an update to our Galaxy-based web server for processing and visualizing deeply sequenced data. Its core tool set, {deepTools}, allows users to perform complete bioinformatic workflows ranging from quality controls and normalizations of aligned reads to integrative analyses, including clustering and visualization approaches. Since we first described our {deepTools} Galaxy server in 2014, we have implemented new solutions for many requests from the community and our users. Here, we introduce significant enhancements and new tools to further improve data visualization and interpretation. {deepTools} continue to be open to all users and freely available as a web service at deeptools.ie-freiburg.mpg.de. The new {deepTools}2 suite can be easily deployed within any Galaxy framework via the toolshed repository, and we also provide source code for command line usage under Linux and Mac {OS} X. A public and documented {API} for access to {deepTools} functionality is also available.},
	pages = {W160--W165},
	issue = {W1},
	journal = {Nucleic Acids Research},
	shortjournal = {Nucleic Acids Research},
	author = {Ramírez, Fidel and Ryan, Devon P and Grüning, Björn and Bhardwaj, Vivek and Kilpert, Fabian and Richter, Andreas S and Heyne, Steffen and Dündar, Friederike and Manke, Thomas},
	urldate = {2022-03-05},
	date = {2016-07-08},
}

@article{glaser_assessing_2021,
	title = {Assessing genome-wide dynamic changes in enhancer activity during early {mESC} differentiation by {FAIRE}-{STARR}-seq},
	year = {2021},
	volume = {49},
	issn = {0305-1048},
	url = {https://doi.org/10.1093/nar/gkab1100},
	doi = {10.1093/nar/gkab1100},
	abstract = {Embryonic stem cells ({ESCs}) can differentiate into any given cell type and therefore represent a versatile model to study the link between gene regulation and differentiation. To quantitatively assess the dynamics of enhancer activity during the early stages of murine {ESC} differentiation, we analyzed accessible genomic regions using {STARR}-seq, a massively parallel reporter assay. This resulted in a genome-wide quantitative map of active {mESC} enhancers, in pluripotency and during the early stages of differentiation. We find that only a minority of accessible regions is active and that such regions are enriched near promoters, characterized by specific chromatin marks, enriched for distinct sequence motifs, and modeling shows that active regions can be predicted from sequence alone. Regions that change their activity upon retinoic acid-induced differentiation are more prevalent at distal intergenic regions when compared to constitutively active enhancers. Further, analysis of differentially active enhancers verified the contribution of individual {TF} motifs toward activity and inducibility as well as their role in regulating endogenous genes. Notably, the activity of retinoic acid receptor alpha ({RARα}) occupied regions can either increase or decrease upon the addition of its ligand, retinoic acid, with the direction of the change correlating with spacing and orientation of the {RARα} consensus motif and the co-occurrence of additional sequence motifs. Together, our genome-wide enhancer activity map elucidates features associated with enhancer activity levels, identifies regulatory regions disregarded by computational prediction tools, and provides a resource for future studies into regulatory elements in {mESCs}.},
	pages = {12178--12195},
	number = {21},
	journal = {Nucleic Acids Research},
	shortjournal = {Nucleic Acids Research},
	author = {Glaser, Laura V and Steiger, Mara and Fuchs, Alisa and van Bömmel, Alena and Einfeldt, Edda and Chung, Ho-Ryun and Vingron, Martin and Meijsing, Sebastiaan H},
	urldate = {2022-03-05},
	date = {2021-12-02},
}

@article{gu_complex_2016,
	title = {Complex heatmaps reveal patterns and correlations in multidimensional genomic data.},
	year = {2016},
	volume = {32},
	rights = {© The Author 2016. Published by Oxford University Press. All rights reserved. For Permissions, please.},
	issn = {1367-4811 1367-4803},
	doi = {10.1093/bioinformatics/btw313},
	abstract = {Parallel heatmaps with carefully designed annotation graphics are powerful for efficient visualization of patterns and relationships among high dimensional genomic  data. Here we present the {ComplexHeatmap} package that provides rich functionalities  for customizing heatmaps, arranging multiple parallel heatmaps and including  user-defined annotation graphics. We demonstrate the power of {ComplexHeatmap} to  easily reveal patterns and correlations among multiple sources of information with  four real-world datasets. {AVAILABILITY} {AND} {IMPLEMENTATION}: The {ComplexHeatmap}  package and documentation are freely available from the Bioconductor project:  http://www.bioconductor.org/packages/devel/bioc/html/{ComplexHeatmap}.html {CONTACT}:  {SUPPLEMENTARY} {INFORMATION}: Supplementary data are available at  Bioinformatics online.},
	pages = {2847--2849},
	number = {18},
	journal = {Bioinformatics (Oxford, England)},
	shortjournal = {Bioinformatics},
	author = {Gu, Zuguang and Eils, Roland and Schlesner, Matthias},
	date = {2016-09-15},
	pmid = {27207943},
	note = {Place: England},
	keywords = {Humans, *Software, *Genomics, Computer Graphics, Gene Expression, Metabolic Networks and Pathways},
}

@article{lawrence_software_2013,
	title = {Software for Computing and Annotating Genomic Ranges},
	year = {2013},
	volume = {9},
	url = {https://doi.org/10.1371/journal.pcbi.1003118},
	doi = {10.1371/journal.pcbi.1003118},
	abstract = {We describe Bioconductor infrastructure for representing and computing on annotated genomic ranges and integrating genomic data with the statistical computing features of R and its extensions. At the core of the infrastructure are three packages: {IRanges}, {GenomicRanges}, and {GenomicFeatures}. These packages provide scalable data structures for representing annotated ranges on the genome, with special support for transcript structures, read alignments and coverage vectors. Computational facilities include efficient algorithms for overlap and nearest neighbor detection, coverage calculation and other range operations. This infrastructure directly supports more than 80 other Bioconductor packages, including those for sequence analysis, differential expression analysis and visualization.},
	pages = {e1003118},
	number = {8},
	journal = {{PLOS} Computational Biology},
	shortjournal = {{PLOS} Computational Biology},
	author = {Lawrence, Michael and Huber, Wolfgang and Pagès, Hervé and Aboyoun, Patrick and Carlson, Marc and Gentleman, Robert and Morgan, Martin T. and Carey, Vincent J.},
	date = {2013-08-08},
	note = {Publisher: Public Library of Science},
}

@article{huang_chromosomal_2021,
	title = {The chromosomal protein {SMCHD}1 regulates {DNA} methylation and the 2c-like state of embryonic stem cells by antagonizing {TET} proteins.},
	year = {2021},
	volume = {7},
	rights = {Copyright © 2021 The Authors, some rights reserved; exclusive licensee American Association for the Advancement of Science. No claim to original U.S. Government  Works. Distributed under a Creative Commons Attribution {NonCommercial} License 4.0  ({CC} {BY}-{NC}).},
	issn = {2375-2548},
	doi = {10.1126/sciadv.abb9149},
	abstract = {5-Methylcytosine (5mC) oxidases, the ten-eleven translocation ({TET}) proteins, initiate {DNA} demethylation, but it is unclear how 5mC oxidation is regulated. We  show that the protein {SMCHD}1 (structural maintenance of chromosomes flexible hinge  domain containing 1) is found in complexes with {TET} proteins and negatively  regulates {TET} activities. Removal of {SMCHD}1 from mouse embryonic stem ({ES}) cells  induces {DNA} hypomethylation, preferentially at {SMCHD}1 target sites and accumulation  of 5-hydroxymethylcytosine (5hmC), along with promoter demethylation and activation  of the Dux double-homeobox gene. In the absence of {SMCHD}1, {ES} cells acquire a  two-cell (2c) embryo-like state characterized by activation of an early embryonic  transcriptome that is substantially imposed by Dux Using Smchd1/Tet1/Tet2/Tet3  quadruple-knockout cells, we show that {DNA} demethylation, activation of Dux, and  other genes upon {SMCHD}1 loss depend on {TET} proteins. These data identify {SMCHD}1 as  an antagonist of the 2c-like state of {ES} cells and of {TET}-mediated {DNA}  demethylation.},
	number = {4},
	journal = {Science advances},
	shortjournal = {Sci Adv},
	author = {Huang, Zhijun and Yu, Jiyoung and Cui, Wei and Johnson, Benjamin K. and Kim, Kyunggon and Pfeifer, Gerd P.},
	date = {2021-01},
	pmid = {33523915},
	pmcid = {PMC7817097},
}

@article{grand_banp_2021,
	title = {{BANP} opens chromatin and activates {CpG}-island-regulated genes},
	year = {2021},
	volume = {596},
	issn = {1476-4687},
	url = {https://doi.org/10.1038/s41586-021-03689-8},
	doi = {10.1038/s41586-021-03689-8},
	abstract = {The majority of gene transcripts generated by {RNA} polymerase {II} in mammalian genomes initiate at {CpG} island ({CGI}) promoters1,2, yet our understanding of their regulation remains limited. This is in part due to the incomplete information that we have on transcription factors, their {DNA}-binding motifs and which genomic binding sites are functional in any given cell type3–5. In addition, there are orphan motifs without known binders, such as the {CGCG} element, which is associated with highly expressed genes across human tissues and enriched near the transcription start site of a subset of {CGI} promoters6–8. Here we combine single-molecule footprinting with interaction proteomics to identify {BTG}3-associated nuclear protein ({BANP}) as the transcription factor that binds this element in the mouse and human genome. We show that {BANP} is a strong {CGI} activator that controls essential metabolic genes in pluripotent stem and terminally differentiated neuronal cells. {BANP} binding is repelled by {DNA} methylation of its motif in vitro and in vivo, which epigenetically restricts most binding to {CGIs} and accounts for differential binding at aberrantly methylated {CGI} promoters in cancer cells. Upon binding to an unmethylated motif, {BANP} opens chromatin and phases nucleosomes. These findings establish {BANP} as a critical activator of a set of essential genes and suggest a model in which the activity of {CGI} promoters relies on methylation-sensitive transcription factors that are capable of chromatin opening.},
	pages = {133--137},
	number = {7870},
	journal = {Nature},
	shortjournal = {Nature},
	author = {Grand, Ralph S. and Burger, Lukas and Gräwe, Cathrin and Michael, Alicia K. and Isbel, Luke and Hess, Daniel and Hoerner, Leslie and Iesmantavicius, Vytautas and Durdu, Sevi and Pregnolato, Marco and Krebs, Arnaud R. and Smallwood, Sébastien A. and Thomä, Nicolas and Vermeulen, Michiel and Schübeler, Dirk},
	date = {2021-08-01},
}

@article{love_moderated_2014,
	title = {Moderated estimation of fold change and dispersion for {RNA}-seq data with {DESeq}2},
	year = {2014},
	volume = {15},
	issn = {1474-760X},
	url = {https://doi.org/10.1186/s13059-014-0550-8},
	doi = {10.1186/s13059-014-0550-8},
	abstract = {In comparative high-throughput sequencing assays, a fundamental task is the analysis of count data, such as read counts per gene in {RNA}-seq, for evidence of systematic changes across experimental conditions. Small replicate numbers, discreteness, large dynamic range and the presence of outliers require a suitable statistical approach. We present {DESeq}2, a method for differential analysis of count data, using shrinkage estimation for dispersions and fold changes to improve stability and interpretability of estimates. This enables a more quantitative analysis focused on the strength rather than the mere presence of differential expression. The {DESeq}2 package is available at http://www.bioconductor.org/packages/release/bioc/html/{DESeq}2.html.},
	pages = {550},
	number = {12},
	journal = {Genome Biology},
	shortjournal = {Genome Biology},
	author = {Love, Michael I. and Huber, Wolfgang and Anders, Simon},
	date = {2014-12-05},
}

@article{bibel_generation_2007,
	title = {Generation of a defined and uniform population of {CNS} progenitors and neurons from mouse embryonic stem cells.},
	year = {2007},
	volume = {2},
	issn = {1750-2799},
	doi = {10.1038/nprot.2007.147},
	abstract = {A detailed protocol is described allowing the generation of essentially pure populations of glutamatergic neurons from mouse embryonic stem ({ES}) cells. It is  based on the culture of {ES} cells that are kept undifferentiated by repeated  splitting and subsequently amplified as non-adherent cell aggregates. Treatment with  retinoic acid causes these {ES} cells to essentially become neural progenitors with  the characteristics of Pax6-positive radial glial cells. As they do in vivo, these  progenitors differentiate in glutamatergic pyramidal neurons that form functional  synaptic contacts and can be kept in culture for long periods of time. This protocol  does not require the use of {ES} lines expressing resistance or fluorescent markers  and can thus be applied in principle to any wild-type or mutant {ES} line of interest.  At least 2 weeks are required from starting {ES} cell culture until plating  progenitors and differentiating neurons establish synaptic transmission within about  10 days.},
	pages = {1034--1043},
	number = {5},
	journal = {Nature protocols},
	shortjournal = {Nat Protoc},
	author = {Bibel, Miriam and Richter, Jens and Lacroix, Emmanuel and Barde, Yves-Alain},
	date = {2007},
	pmid = {17546008},
	note = {Place: England},
	keywords = {Animals, Mice, Cell Culture Techniques/*methods, Cell Differentiation/*drug effects, Central Nervous System/*cytology, Embryonic Stem Cells/*cytology, Neurons/*cytology, Tretinoin/pharmacology},
}

@article{keane_mouse_2011,
	title = {Mouse genomic variation and its effect on phenotypes and gene regulation},
	year = {2011},
	volume = {477},
	issn = {1476-4687},
	url = {https://doi.org/10.1038/nature10413},
	doi = {10.1038/nature10413},
	abstract = {We report genome sequences of 17 inbred strains of laboratory mice and identify almost ten times more variants than previously known. We use these genomes to explore the phylogenetic history of the laboratory mouse and to examine the functional consequences of allele-specific variation on transcript abundance, revealing that at least 12\% of transcripts show a significant tissue-specific expression bias. By identifying candidate functional variants at 718 quantitative trait loci we show that the molecular nature of functional variants and their position relative to genes vary according to the effect size of the locus. These sequences provide a starting point for a new era in the functional analysis of a key model organism.},
	pages = {289--294},
	number = {7364},
	journal = {Nature},
	shortjournal = {Nature},
	author = {Keane, Thomas M. and Goodstadt, Leo and Danecek, Petr and White, Michael A. and Wong, Kim and Yalcin, Binnaz and Heger, Andreas and Agam, Avigail and Slater, Guy and Goodson, Martin and Furlotte, Nicholas A. and Eskin, Eleazar and Nellåker, Christoffer and Whitley, Helen and Cleak, James and Janowitz, Deborah and Hernandez-Pliego, Polinka and Edwards, Andrew and Belgard, T. Grant and Oliver, Peter L. and {McIntyre}, Rebecca E. and Bhomra, Amarjit and Nicod, Jérôme and Gan, Xiangchao and Yuan, Wei and van der Weyden, Louise and Steward, Charles A. and Bala, Sendu and Stalker, Jim and Mott, Richard and Durbin, Richard and Jackson, Ian J. and Czechanski, Anne and Guerra-Assunção, José Afonso and Donahue, Leah Rae and Reinholdt, Laura G. and Payseur, Bret A. and Ponting, Chris P. and Birney, Ewan and Flint, Jonathan and Adams, David J.},
	date = {2011-09-01},
}

@misc{patel_nf-corernaseq_2022,
	title = {nf-core/rnaseq: nf-core/rnaseq v3.6 - Platinum Platypus},
	year = {2022},
	howpublished = {\url{https://doi.org/10.5281/zenodo.6327553#.YifOASRAq5E.mendeley}},
	doi = {10.5281/ZENODO.6327553},
	author = {Patel, Harshil and Ewels, Phil and Peltzer, Alexander and Hammarén, Rickard and Botvinnik, Olga and Sturm, Gregor and Moreno, Denis and Vemuri, Pranathi and {silviamorins} and Pantano, Lorena and Binzer-Panchal, Mahesh and Kelly, Gavin and {FriederikeHanssen} and Garcia, Maxime U. and bot, nf-core and Cheshire, Chris and {rfenouil} and Espinosa-Carrasco, Jose and {marchoeppner} and Zhou, Peng and Gabernet, Gisela and Mertes, Christian and Straub, Daniel and Hörtenhuber, Matthias and Tommaso, Paolo Di and F, Sven and Hall, George and Panneerselvam, Senthilkumar and OMeally, Denis and {jun-wan}},
	date = {2022-03},
}

@article{ewels_nf-core_2020,
	title = {The nf-core framework for community-curated bioinformatics pipelines},
	year = {2020},
	volume = {38},
	issn = {1546-1696},
	url = {https://doi.org/10.1038/s41587-020-0439-x},
	doi = {10.1038/s41587-020-0439-x},
	pages = {276--278},
	number = {3},
	journal = {Nature Biotechnology},
	shortjournal = {Nature Biotechnology},
	author = {Ewels, Philip A. and Peltzer, Alexander and Fillinger, Sven and Patel, Harshil and Alneberg, Johannes and Wilm, Andreas and Garcia, Maxime Ulysse and Di Tommaso, Paolo and Nahnsen, Sven},
	date = {2020-03-01},
}

@article{heinz_simple_2010,
	title = {Simple combinations of lineage-determining transcription factors prime cis-regulatory elements required for macrophage and B cell identities.},
	year = {2010},
	volume = {38},
	rights = {Copyright 2010 Elsevier Inc. All rights reserved.},
	issn = {1097-4164 1097-2765},
	doi = {10.1016/j.molcel.2010.05.004},
	abstract = {Genome-scale studies have revealed extensive, cell type-specific colocalization of transcription factors, but the mechanisms underlying this phenomenon remain poorly  understood. Here, we demonstrate in macrophages and B cells that collaborative  interactions of the common factor {PU}.1 with small sets of macrophage- or B cell  lineage-determining transcription factors establish cell-specific binding sites that  are associated with the majority of promoter-distal H3K4me1-marked genomic regions.  {PU}.1 binding initiates nucleosome remodeling, followed by H3K4 monomethylation at  large numbers of genomic regions associated with both broadly and specifically  expressed genes. These locations serve as beacons for additional factors,  exemplified by liver X receptors, which drive both cell-specific gene expression and  signal-dependent responses. Together with analyses of transcription factor binding  and H3K4me1 patterns in other cell types, these studies suggest that simple  combinations of lineage-determining transcription factors can specify the genomic  sites ultimately responsible for both cell identity and cell type-specific responses  to diverse signaling inputs.},
	pages = {576--589},
	number = {4},
	journal = {Molecular cell},
	shortjournal = {Mol Cell},
	author = {Heinz, Sven and Benner, Christopher and Spann, Nathanael and Bertolino, Eric and Lin, Yin C. and Laslo, Peter and Cheng, Jason X. and Murre, Cornelis and Singh, Harinder and Glass, Christopher K.},
	date = {2010-05-28},
	pmid = {20513432},
	pmcid = {PMC2898526},
	keywords = {Animals, Binding Sites, Mice, Male, Mice, Inbred C57BL, *Cell Lineage/genetics, B-Lymphocytes/cytology/*metabolism, Macrophages/cytology/*metabolism, Oligonucleotide Array Sequence Analysis, Proto-Oncogene Proteins/genetics/*metabolism, Regulatory Elements, Transcriptional/*genetics, Trans-Activators/genetics/*metabolism, Transcription Factors/genetics/*metabolism},
}

@article{das_distinct_2014,
	title = {Distinct and combinatorial functions of Jmjd2b/Kdm4b and Jmjd2c/Kdm4c in mouse embryonic stem cell identity.},
	year = {2014},
	volume = {53},
	rights = {Copyright © 2014 Elsevier Inc. All rights reserved.},
	issn = {1097-4164 1097-2765},
	doi = {10.1016/j.molcel.2013.11.011},
	abstract = {Self-renewal and pluripotency of embryonic stem cells ({ESCs}) are established by multiple regulatory pathways operating at several levels. The roles of histone  demethylases ({HDMs}) in these programs are incompletely defined. We conducted a  functional {RNAi} screen for {HDMs} and identified five potential {HDMs} essential for  mouse {ESC} identity. In-depth analyses demonstrate that the closely related {HDMs}  Jmjd2b and Jmjd2c are necessary for self-renewal of {ESCs} and induced pluripotent  stem cell generation. Genome-wide occupancy studies reveal that Jmjd2b unique,  Jmjd2c unique, and Jmjd2b-Jmjd2c common target sites belong to functionally  separable Core, Polycomb repressive complex ({PRC}), and Myc regulatory modules,  respectively. Jmjd2b and Nanog act through an interconnected regulatory loop,  whereas Jmjd2c assists {PRC}2 in transcriptional repression. Thus, two {HDMs} of the  same subclass exhibit distinct and combinatorial functions in control of the {ESC}  state. Such complexity of {HDM} function reveals an aspect of multilayered  transcriptional control.},
	pages = {32--48},
	number = {1},
	journal = {Molecular cell},
	shortjournal = {Mol Cell},
	author = {Das, Partha Pratim and Shao, Zhen and Beyaz, Semir and Apostolou, Eftychia and Pinello, Luca and De Los Angeles, Alejandro and O'Brien, Kassandra and Atsma, Jennifer Marino and Fujiwara, Yuko and Nguyen, Minh and Ljuboja, Damir and Guo, Guoji and Woo, Andrew and Yuan, Guo-Cheng and Onder, Tamer and Daley, George and Hochedlinger, Konrad and Kim, Jonghwan and Orkin, Stuart H.},
	date = {2014-01-09},
	pmid = {24361252},
	pmcid = {PMC3919500},
	keywords = {Animals, Cell Line, Mice, Embryonic Stem Cells/cytology/*enzymology, Genome-Wide Association Study, Homeodomain Proteins/genetics/metabolism, Jumonji Domain-Containing Histone Demethylases/genetics/*metabolism, Nanog Homeobox Protein, Pluripotent Stem Cells/cytology/*enzymology, Polycomb Repressive Complex 2/genetics/metabolism, Proto-Oncogene Proteins c-myc/genetics/metabolism, Transcription, Genetic/*physiology},
}

@article{chronis_cooperative_2017,
	title = {Cooperative Binding of Transcription Factors Orchestrates Reprogramming.},
	year = {2017},
	volume = {168},
	rights = {Copyright © 2017 Elsevier Inc. All rights reserved.},
	issn = {1097-4172 0092-8674},
	doi = {10.1016/j.cell.2016.12.016},
	abstract = {Oct4, Sox2, Klf4, and {cMyc} ({OSKM}) reprogram somatic cells to pluripotency. To gain a mechanistic understanding of their function, we mapped {OSKM}-binding, stage-specific  transcription factors ({TFs}), and chromatin states in discrete reprogramming stages  and performed loss- and gain-of-function experiments. We found that {OSK}  predominantly bind active somatic enhancers early in reprogramming and immediately  initiate their inactivation genome-wide by inducing the redistribution of somatic  {TFs} away from somatic enhancers to sites elsewhere engaged by {OSK}, recruiting Hdac1,  and repressing the somatic {TF} Fra1. Pluripotency enhancer selection is a stepwise  process that also begins early in reprogramming through collaborative binding of {OSK}  at sites with high {OSK}-motif density. Most pluripotency enhancers are selected later  in the process and require {OS} and other pluripotency {TFs}. Somatic and pluripotency  {TFs} modulate reprogramming efficiency when overexpressed by altering {OSK} targeting,  somatic-enhancer inactivation, and pluripotency enhancer selection. Together, our  data indicate that collaborative interactions among {OSK} and with stage-specific {TFs}  direct both somatic-enhancer inactivation and pluripotency-enhancer selection to  drive reprogramming.},
	pages = {442--459.e20},
	number = {3},
	journal = {Cell},
	shortjournal = {Cell},
	author = {Chronis, Constantinos and Fiziev, Petko and Papp, Bernadett and Butz, Stefan and Bonora, Giancarlo and Sabri, Shan and Ernst, Jason and Plath, Kathrin},
	date = {2017-01-26},
	pmid = {28111071},
	pmcid = {PMC5302508},
	keywords = {Animals, Mice, Chromatin/metabolism, *Cellular Reprogramming, *collaborative binding, *enhancers, *induced pluripotent stem cells, *Klf4, *Oct4, *pluripotency, *reprogramming, *Sox2, *transcription factors, Fibroblasts/metabolism, Histone Code, Kruppel-Like Factor 4, Kruppel-Like Transcription Factors/metabolism, Octamer Transcription Factor-3/metabolism, Proto-Oncogene Proteins c-fos/metabolism, Proto-Oncogene Proteins c-myc/metabolism, Regulatory Elements, Transcriptional, Silencer Elements, Transcriptional, {SOXB}1 Transcription Factors/metabolism, Transcription Factors/*metabolism},
}

@article{mor_neutralizing_2018,
	title = {Neutralizing Gatad2a-Chd4-Mbd3/{NuRD} Complex Facilitates Deterministic Induction of Naive Pluripotency.},
	year = {2018},
	volume = {23},
	rights = {Copyright © 2018 Elsevier Inc. All rights reserved.},
	issn = {1875-9777 1934-5909},
	doi = {10.1016/j.stem.2018.07.004},
	abstract = {Mbd3, a member of nucleosome remodeling and deacetylase ({NuRD}) co-repressor complex, was previously identified as an inhibitor for deterministic induced pluripotent stem  cell ({iPSC}) reprogramming, where up to 100\% of donor cells successfully complete the  process. {NuRD} can assume multiple mutually exclusive conformations, and it remains  unclear whether this deterministic phenotype can be attributed to a specific  Mbd3/{NuRD} subcomplex. Moreover, since complete ablation of Mbd3 blocks somatic cell  proliferation, we aimed to explore functionally relevant alternative ways to  neutralize Mbd3-dependent {NuRD} activity. We identify Gatad2a, a {NuRD}-specific  subunit, whose complete deletion specifically disrupts Mbd3/{NuRD} repressive activity  on the pluripotency circuitry during {iPSC} differentiation and reprogramming without  ablating somatic cell proliferation. Inhibition of Gatad2a facilitates deterministic  murine {iPSC} reprogramming within 8 days. We validate a distinct molecular axis,  Gatad2a-Chd4-Mbd3, within Mbd3/{NuRD} as being critical for blocking reestablishment  of naive pluripotency and further highlight signaling-dependent and  post-translational modifications of Mbd3/{NuRD} that influence its interactions and  assembly.},
	pages = {412--425.e10},
	number = {3},
	journal = {Cell stem cell},
	shortjournal = {Cell Stem Cell},
	author = {Mor, Nofar and Rais, Yoach and Sheban, Daoud and Peles, Shani and Aguilera-Castrejon, Alejandro and Zviran, Asaf and Elinger, Dalia and Viukov, Sergey and Geula, Shay and Krupalnik, Vladislav and Zerbib, Mirie and Chomsky, Elad and Lasman, Lior and Shani, Tom and Bayerl, Jonathan and Gafni, Ohad and Hanna, Suhair and Buenrostro, Jason D. and Hagai, Tzachi and Masika, Hagit and Vainorius, Gintautas and Bergman, Yehudit and Greenleaf, William J. and Esteban, Miguel A. and Elling, Ulrich and Levin, Yishai and Massarwa, Rada and Merbl, Yifat and Novershtern, Noa and Hanna, Jacob H.},
	date = {2018-09-06},
	pmid = {30122475},
	pmcid = {PMC7116536},
	keywords = {Animals, Mice, Male, Mice, Inbred C57BL, *pluripotency, *reprogramming, Transcription Factors/*metabolism, *{CHD}4, *epigenetics, *Gatad2a, *{iPSCs}, *Mbd3, *{NuRD}, *P66α, Cells, Cultured, {DNA} Helicases/*metabolism, {DNA}-Binding Proteins/*metabolism, Female, {GATA} Transcription Factors/*metabolism, Induced Pluripotent Stem Cells/cytology/*metabolism, Mi-2 Nucleosome Remodeling and Deacetylase Complex/*metabolism, Mice, Inbred {CBA}, Mice, Knockout, Mice, Transgenic},
}

@article{xu_genome-wide_2011,
	title = {Genome-wide regulation of 5hmC, 5mC, and gene expression by Tet1 hydroxylase in mouse embryonic stem cells.},
	year = {2011},
	volume = {42},
	rights = {Copyright © 2011 Elsevier Inc. All rights reserved.},
	issn = {1097-4164 1097-2765},
	doi = {10.1016/j.molcel.2011.04.005},
	abstract = {{DNA} methylation at the 5 position of cytosine (5mC) in the mammalian genome is a key epigenetic event critical for various cellular processes. The ten-eleven  translocation (Tet) family of 5mC-hydroxylases, which convert 5mC to  5-hydroxymethylcytosine (5hmC), offers a way for dynamic regulation of {DNA}  methylation. Here we report that Tet1 binds to unmodified C or 5mC- or 5hmC-modified  {CpG}-rich {DNA} through its {CXXC} domain. Genome-wide mapping of Tet1 and 5hmC reveals  mechanisms by which Tet1 controls 5hmC and 5mC levels in mouse embryonic stem cells  ({mESCs}). We also uncover a comprehensive gene network influenced by Tet1.  Collectively, our data suggest that Tet1 controls {DNA} methylation both by binding to  {CpG}-rich regions to prevent unwanted {DNA} methyltransferase activity, and by  converting 5mC to 5hmC through hydroxylase activity. This Tet1-mediated antagonism  of {CpG} methylation imparts differential maintenance of {DNA} methylation status at  Tet1 targets, ultimately contributing to {mESC} differentiation and the onset of  embryonic development.},
	pages = {451--464},
	number = {4},
	journal = {Molecular cell},
	shortjournal = {Mol Cell},
	author = {Xu, Yufei and Wu, Feizhen and Tan, Li and Kong, Lingchun and Xiong, Lijun and Deng, Jie and Barbera, Andrew J. and Zheng, Lijuan and Zhang, Haikuo and Huang, Stephen and Min, Jinrong and Nicholson, Thomas and Chen, Taiping and Xu, Guoliang and Shi, Yang and Zhang, Kun and Shi, Yujiang Geno},
	date = {2011-05-20},
	pmid = {21514197},
	pmcid = {PMC3099128},
	keywords = {Animals, Mice, Protein Binding, *Gene Expression Regulation, Developmental, Embryonic Stem Cells/*enzymology, Genome-Wide Association Study, {DNA}-Binding Proteins/*metabolism, 5-Methylcytosine/*metabolism, Cytosine/*analogs \& derivatives/metabolism, {DNA} (Cytosine-5-)-Methyltransferase 1, {DNA} (Cytosine-5-)-Methyltransferases/antagonists \& inhibitors, Mixed Function Oxygenases/*metabolism, Protein Structure, Tertiary, Proto-Oncogene Proteins/*metabolism},
}

@misc{team_bc_txdbmmusculusucscmm10knowngene_2019,
	title = {{TxDb}.Mmusculus.{UCSC}.mm10.{knownGene}: Annotation package for {TxDb} object(s)},
	year = {2019},
	howpublished = {\url{http://https/packages/TxDb.Mmusculus.UCSC.mm10.knownGene/}},
	doi = {10.18129/B9.bioc.TxDb.Mmusculus.UCSC.mm10.knownGene},
	author = {Team {BC} and Maintainer {BP}},
	date = {2019},
}

@article{giorgetti_structural_2016,
	title = {Structural organization of the inactive X chromosome in the mouse.},
	year = {2016},
	volume = {535},
	issn = {1476-4687 0028-0836},
	doi = {10.1038/nature18589},
	abstract = {X-chromosome inactivation ({XCI}) involves major reorganization of the X chromosome as it becomes silent and heterochromatic. During female mammalian development, {XCI} is  triggered by upregulation of the non-coding Xist {RNA} from one of the two X  chromosomes. Xist coats the chromosome in cis and induces silencing of almost all  genes via its A-repeat region, although some genes (constitutive escapees) avoid  silencing in most cell types, and others (facultative escapees) escape {XCI} only in  specific contexts. A role for Xist in organizing the inactive X (Xi) chromosome has  been proposed. Recent chromosome conformation capture approaches have revealed  global loss of local structure on the Xi chromosome and formation of large  mega-domains, separated by a region containing the {DXZ}4 macrosatellite. However, the  molecular architecture of the Xi chromosome, in both the silent and expressed  regions,remains unclear. Here we investigate the structure, chromatin accessibility  and expression status of the mouse Xi chromosome in highly polymorphic clonal neural  progenitors ({NPCs}) and embryonic stem cells. We demonstrate a crucial role for Xist  and the {DXZ}4-containing boundary in shaping Xi chromosome structure using  allele-specific genome-wide chromosome conformation capture (Hi-C) analysis, an  assay for transposase-accessible chromatin with high throughput sequencing  ({ATAC}-seq) and {RNA} sequencing. Deletion of the boundary disrupts mega-domain  formation, and induction of Xist {RNA} initiates formation of the boundary and the  loss of {DNA} accessibility. We also show that in {NPCs}, the Xi chromosome lacks  active/inactive compartments and topologically associating domains ({TADs}), except  around genes that escape {XCI}. Escapee gene clusters display {TAD}-like structures and  retain {DNA} accessibility at promoter-proximal and {CTCF}-binding sites. Furthermore,  altered patterns of facultative escape genes indifferent neural progenitor clones  are associated with the presence of different {TAD}-like structures after {XCI}. These  findings suggest a key role for transcription and {CTCF} in the formation of {TADs} in  the context of the Xi chromosome in neural progenitors.},
	pages = {575--579},
	number = {7613},
	journal = {Nature},
	shortjournal = {Nature},
	author = {Giorgetti, Luca and Lajoie, Bryan R. and Carter, Ava C. and Attia, Mikael and Zhan, Ye and Xu, Jin and Chen, Chong Jian and Kaplan, Noam and Chang, Howard Y. and Heard, Edith and Dekker, Job},
	date = {2016-07-28},
	pmid = {27437574},
	pmcid = {PMC5443622},
	keywords = {Animals, Binding Sites, Mice, Transcription, Genetic, Alleles, Male, Promoter Regions, Genetic/genetics, Female, *X Chromosome Inactivation/genetics, {CCCTC}-Binding Factor, Chromatin/chemistry/genetics/metabolism, Chromosomes, Mammalian/chemistry/genetics/*metabolism, Embryonic Stem Cells/metabolism, Gene Silencing, Neural Stem Cells/metabolism, Repressor Proteins/metabolism, {RNA}, Long Noncoding/genetics/metabolism, Sequence Analysis, X Chromosome/chemistry/genetics/*metabolism},
}

@article{habibi_whole-genome_2013,
	title = {Whole-genome bisulfite sequencing of two distinct interconvertible {DNA} methylomes of mouse embryonic stem cells.},
	year = {2013},
	volume = {13},
	rights = {Copyright © 2013 Elsevier Inc. All rights reserved.},
	issn = {1875-9777},
	doi = {10.1016/j.stem.2013.06.002},
	abstract = {The use of two kinase inhibitors (2i) enables derivation of mouse embryonic stem cells ({ESCs}) in the pluripotent ground state. Using whole-genome bisulfite  sequencing ({WGBS}), we show that male 2i {ESCs} are globally hypomethylated compared to  conventional {ESCs} maintained in serum. In serum, female {ESCs} are hypomethyated  similarly to male {ESCs} in 2i, and {DNA} methylation is further reduced in 2i. Regions  with elevated {DNA} methylation in 2i strongly correlate with the presence of H3K9me3  on endogenous retroviruses ({ERVs}) and imprinted loci. The methylome of male {ESCs} in  serum parallels postimplantation blastocyst cells, while 2i stalls {ESCs} in a  hypomethylated, {ICM}-like state. {WGBS} analysis during adaptation of 2i {ESCs} to serum  suggests that deposition of {DNA} methylation is largely random, while loss of {DNA}  methylation during reversion to 2i occurs passively, initiating at {TET}1 binding  sites. Together, our analysis provides insight into {DNA} methylation dynamics in  cultured {ESCs} paralleling early developmental processes.},
	pages = {360--369},
	number = {3},
	journal = {Cell stem cell},
	shortjournal = {Cell Stem Cell},
	author = {Habibi, Ehsan and Brinkman, Arie B. and Arand, Julia and Kroeze, Leonie I. and Kerstens, Hindrik H. D. and Matarese, Filomena and Lepikhov, Konstantin and Gut, Marta and Brun-Heath, Isabelle and Hubner, Nina C. and Benedetti, Rosaria and Altucci, Lucia and Jansen, Joop H. and Walter, Jörn and Gut, Ivo G. and Marks, Hendrik and Stunnenberg, Hendrik G.},
	date = {2013-09-05},
	pmid = {23850244},
	note = {Place: United States},
	keywords = {Animals, Histones/metabolism, Mice, Sequence Analysis, {DNA}, Male, Methylation, Proto-Oncogene Proteins/genetics/*metabolism, Cells, Cultured, Female, Blastocyst/*physiology, {DNA} Methylation/drug effects, {DNA}-Binding Proteins/genetics/*metabolism, Embryonic Stem Cells/drug effects/*physiology, Fetal Development, Genome/genetics, Histone Demethylases/*metabolism, Leukemia Inhibitory Factor/metabolism, Pluripotent Stem Cells/*physiology, Protein Kinase Inhibitors/pharmacology, Sulfites/chemistry},
}

@article{mattei_dna_2022,
	title = {{DNA} methylation: a historical perspective},
	year = {2022},
	issn = {0168-9525},
	url = {https://www.sciencedirect.com/science/article/pii/S0168952522000713},
	doi = {10.1016/j.tig.2022.03.010},
	abstract = {In 1925, 5-methylcytosine was first reported in bacteria. However, its biological importance was not intuitive for several decades. After this initial lag, the ubiquitous presence of this methylated base emerged across all domains of life and revealed a range of essential biological functions. Today, we are armed with the knowledge of the key factors that establish, maintain, and remove {DNA} methylation and have access to a staggering and rapidly growing number of base-resolution methylation maps. Despite this, several fundamental details about the precise role and interpretation of {DNA} methylation patterns remain under investigation. Here, we review the field of {DNA} methylation from its beginning to present day, with an emphasis on findings in mammalian systems, and point the reader to select experiments that form the foundation of this field.},
	journal = {Trends in Genetics},
	shortjournal = {Trends in Genetics},
	author = {Mattei, Alexandra L. and Bailly, Nina and Meissner, Alexander},
	date = {2022-04-30},
	keywords = {5-methylcytosine, development, {DNA} methylation, epigenetics, gene regulation},
}

@article{walsh_transcription_1998,
	title = {Transcription of {IAP} endogenous retroviruses is constrained by cytosine methylation},
	year = {1998},
	volume = {20},
	issn = {1546-1718},
	url = {https://doi.org/10.1038/2413},
	doi = {10.1038/2413},
	pages = {116--117},
	number = {2},
	journal = {Nature Genetics},
	shortjournal = {Nature Genetics},
	author = {Walsh, Colum P. and Chaillet, J. Richard and Bestor, Timothy H.},
	date = {1998-10-01},
}

@article{ren_detecting_2018,
	title = {Detecting and interpreting {DNA} methylation marks},
	year = {2018},
	volume = {53},
	issn = {0959-440X},
	url = {https://www.sciencedirect.com/science/article/pii/S0959440X17302026},
	doi = {10.1016/j.sbi.2018.06.004},
	abstract = {The generation, alteration, recognition, and erasure of epigenetic modifications of {DNA} are fundamental to controlling gene expression in mammals. These covalent {DNA} modifications include cytosine methylation by {AdoMet}-dependent methyltransferases and 5-methylcytosine oxidation by Fe({II})-dependent and α-ketoglutarate-dependent dioxygenases. Sequence-specific transcription factors are responsible for interpreting the modification status of specific regions of chromatin. This review focuses on recent developments in characterizing the functional and structural links between the modification status of two {DNA} bases: 5-methylcytosine and 5-methyluracil (thymine).},
	pages = {88--99},
	journal = {Protein–nucleic acid interactions • Catalysis and regulation},
	shortjournal = {Current Opinion in Structural Biology},
	author = {Ren, Ren and Horton, John R and Zhang, Xing and Blumenthal, Robert M and Cheng, Xiaodong},
	date = {2018-12-01},
}

@article{cusack_distinct_2020,
	title = {Distinct contributions of {DNA} methylation and histone acetylation to the genomic occupancy of transcription factors.},
	year = {2020},
	volume = {30},
	rights = {© 2020 Cusack et al.; Published by Cold Spring Harbor Laboratory Press.},
	issn = {1549-5469 1088-9051},
	doi = {10.1101/gr.257576.119},
	abstract = {Epigenetic modifications on chromatin play important roles in regulating gene expression. Although chromatin states are often governed by multilayered  structure, how individual pathways contribute to gene expression remains poorly  understood. For example, {DNA} methylation is known to regulate transcription  factor binding but also to recruit methyl-{CpG} binding proteins that affect  chromatin structure through the activity of histone deacetylase complexes  ({HDACs}). Both of these mechanisms can potentially affect gene expression, but the  importance of each, and whether these activities are integrated to achieve  appropriate gene regulation, remains largely unknown. To address this important  question, we measured gene expression, chromatin accessibility, and transcription  factor occupancy in wild-type or {DNA} methylation-deficient mouse embryonic stem  cells following {HDAC} inhibition. We observe widespread increases in chromatin  accessibility at retrotransposons when {HDACs} are inhibited, and this is magnified  when cells also lack {DNA} methylation. A subset of these elements has elevated  binding of the {YY}1 and {GABPA} transcription factors and increased expression. The  pronounced additive effect of {HDAC} inhibition in {DNA} methylation-deficient cells  demonstrates that {DNA} methylation and histone deacetylation act largely  independently to suppress transcription factor binding and gene expression.},
	pages = {1393--1406},
	number = {10},
	journal = {Genome research},
	shortjournal = {Genome Res},
	author = {Cusack, Martin and King, Hamish W. and Spingardi, Paolo and Kessler, Benedikt M. and Klose, Robert J. and Kriaucionis, Skirmantas},
	date = {2020-10},
	pmid = {32963030},
	pmcid = {PMC7605266},
	keywords = {Acetylation, Chromatin/metabolism, Transcription Factors/*metabolism, *{DNA} Methylation, *Epigenesis, Genetic, Embryonic Stem Cells/drug effects/enzymology/metabolism, Genome, Histone Deacetylase Inhibitors, Histone Deacetylases/*metabolism/pharmacology, Histones/*metabolism, Retroelements},
}

@article{long_ever-changing_2016,
	title = {Ever-Changing Landscapes: Transcriptional Enhancers in Development and Evolution},
	year = {2016},
	volume = {167},
	issn = {0092-8674},
	url = {https://www.sciencedirect.com/science/article/pii/S009286741631251X},
	doi = {10.1016/j.cell.2016.09.018},
	abstract = {A class of cis-regulatory elements, called enhancers, play a central role in orchestrating spatiotemporally precise gene-expression programs during development. Consequently, divergence in enhancer sequence and activity is thought to be an important mediator of inter- and intra-species phenotypic variation. Here, we give an overview of emerging principles of enhancer function, current models of enhancer architecture, genomic substrates from which enhancers emerge during evolution, and the influence of three-dimensional genome organization on long-range gene regulation. We discuss intricate relationships between distinct elements within complex regulatory landscapes and consider their potential impact on specificity and robustness of transcriptional regulation.},
	pages = {1170--1187},
	number = {5},
	journal = {Cell},
	shortjournal = {Cell},
	author = {Long, Hannah K. and Prescott, Sara L. and Wysocka, Joanna},
	date = {2016-11-17},
}
