## Supplementary material for "Single molecule multi-omics reveals context-dependent regulation of enhancers by DNA methylation": LaTeX files: Article.tex

\documentclass[times, twoside, watermark]{zHenriquesLab-StyleBioRxiv}
\usepackage{blindtext}
\usepackage{indentfirst}

% Please give the surname of the lead author for the running footer
\leadauthor{Kreibich} 
\setlength{\parindent}{2em}
\DeclareCaptionLabelFormat{adja-page}{\hrulefill\\#1 #2 \emph{(previous page)}}


\begin{document}

\title{Single molecule multi-omics reveals context-dependent regulation of enhancers by DNA methylation}
\shorttitle{Context-dependent regulation of enhancers by DNA methylation}

% Use letters for affiliations, numbers to show equal authorship (if applicable) and to indicate the corresponding author
\author[1,2]{Elisa Kreibich}\author[1]{Rozemarijn Kleinendorst}\author[1,2]{Guido Barzaghi}\author[1]{Sarah Kaspar}\author[1,\Letter]{Arnaud R Krebs}


\affil[1]{European Molecular Biology Laboratory (EMBL), Genome Biology unit, Meyerhofstraße 1, 69117 Heidelberg.Faculty of Biosciences}\affil[2]{Collaboration for Joint PhD Degree between EMBL and Heidelberg University, Heidelberg, Germany.}

\maketitle

%TC:break Abstract
%the command above serves to have a word count for the abstract
\begin{abstract}
Enhancers are \emph{cis}-regulatory elements that control the establishment of cell identities during development. In mammals, enhancer activation is tightly coupled with local DNA demethylation. Yet, whether this epigenetic remodelling is necessary for enhancer activation is unknown. Here, we developed a single molecule multi-omics approach to measure chromatin accessibility and transcription factor binding as a function of the presence of methylation on the same DNA molecules. We leveraged natural epigenetic heterogeneity at active enhancers to test the impact of DNA methylation on their activity in multiple cell lineages. While reduction of DNA methylation appears dispensable for the activity of most enhancers, we identify a class of cell-type-specific enhancers where presence of DNA methylation antagonises the binding of transcription factors. Genetic perturbations reveal that chromatin accessibility and transcription factor binding are a direct function of active demethylation at these loci. Thus, in addition to safeguarding the genome from spurious activation, DNA methylation directly controls transcription factor occupancy at cell-type-specific active enhancers.
\end {abstract}
%TC:break main
%the command above serves to have a word count for the abstract

\noindent\begin{keywords}
DNA methylation | epigenetics | enhancers | transcription factors
\end{keywords}

\begin{corrauthor}
%\texttt{r.henriques{@}ucl.ac.uk}
arnaud.krebs\at embl.de
\end{corrauthor}

\section*{Introduction}
In mammals, DNA methylation (5mC) is a repressive epigenetic mark that occurs on the 5th carbon of cytosines in the CpG dinucleotide context. Shortly after implantation, 70-80\% of CpGs in the genome become methylated \cite{iurlaro_dna_2017, luo_dynamic_2018}. Acquisition of 5mC is critical for the transcriptional silencing of repetitive elements \cite{greenberg_diverse_2019,iurlaro_dna_2017,walsh_transcription_1998}, the maintenance of the mono-allelic expression of genes within Imprinting Control Regions (ICRs) \cite{ferguson-smith_discovery_2018, greenberg_diverse_2019} and the repression of germline gene expression programs \cite{dahlet_genome-wide_2020}. Pervasive methylation is a consistent feature shared between most embryonic and somatic cell types \cite{ehrlich_amount_1982, lister_human_2009, meissner_genome-scale_2008, stadler_dna-binding_2011, ziller_charting_2013} and genetic deletion of DNA methyltransferases (DNMTs) is lethal at early stages of embryogenesis \cite{li_targeted_1992, sakaue_dna_2010}. Transcriptional repression by 5mC has been shown to occur indirectly through the recruitment of repressive complexes that compact chromatin \cite{baubec_genomic_2014}, and also by direct inhibition of transcription factor (TF) binding \cite{kribelbauer_toward_2020, ren_detecting_2018}. Large-scale \emph{in vitro} binding studies have shown that for many TFs the affinity for their recognition motif is altered by the presence of 5mC, and that these effects vary depending on the sequence context where methylation occurs \cite{kribelbauer_quantitative_2017,yin_impact_2017}. Moreover, \emph{in vivo} perturbation of methylation levels has shown that 5mC modulates the binding of NRF1 \cite{domcke_competition_2015}, CTCF \cite{maurano_role_2015}, Max-Myc \cite{cusack_distinct_2020, domcke_competition_2015}, Oct4 \cite{yin_impact_2017} and BANP \cite{grand_banp_2021}, preventing their binding outside of \emph{cis}-regulatory elements (CREs) and controlling the formation of cryptic transcription sites. 

Enhancers are critical for the precise control of genes involved in the acquisition and maintenance of cellular identity \cite{long_ever-changing_2016}. Upon activation, they show a characteristic reduction in the level of 5mC, which has been observed across a large collection of tissues and cell types \cite{hartl_cis-regulatory_2017,hodges_directional_2011,stadler_dna-binding_2011,ziller_charting_2013}. This makes enhancers the most variable class of differentially methylated elements \cite{luo_dynamic_2018}, and suggesting that removal of methylation could be an important step in their activation \cite{luo_dynamic_2018,parry_active_2021}. Indeed, binding of TFs at methylated regions has been shown to cause a local reduction of 5mC \cite{stadler_dna-binding_2011}. Moreover, active enhancers experience high 5mC turnover \cite{ginno_genome-scale_2020}, and have elevated activity of the ten-eleven translocation (TET) enzymes that initiate active DNA demethylation \cite{lu_role_2014}. Together, this provides a mechanistic model for the dynamic methylation reduction observed at enhancers.

The average methylation at active enhancers typically ranges from 10-60\% \cite{hodges_directional_2011,stadler_dna-binding_2011}. This incomplete demethylation indicates a cell-to-cell variability in their methylation level within phenotypically homogeneous cell populations. This epigenetic heterogeneity has been directly quantified by single-molecule analysis of whole genome bisulfite sequencing (WGBS) \cite{landan_epigenetic_2012,shipony_dynamic_2014} and single-cell methylation profiling studies \cite{farlik_single-cell_2015,smallwood_single-cell_2014}. Using a single-cell fluorescent reporter system, it was recently shown that 5mC heterogeneity modulates the activity of two >10kb-large enhancer clusters (super-enhancers) that are involved in controlling the expression of essential genes in mouse Embryonic Stem Cells (mESCs) \cite{song_dynamic_2019}.

The bidirectional regulation between TF binding and 5mC makes it challenging to determine if TF occupancy at enhancers is a cause or a consequence of 5mC removal. A prerequisite to study this complex interplay is to identify enhancers that are regulated by 5mC. Heterogeneity of DNA methylation patterns indicates that within a homogeneous cellular population the DNA sequence of most enhancers exists at variable proportions of unmethylated or methylated states. Here, we exploit this naturally occurring differential methylation at CREs to ask if the DNA molecules marked by 5mC at enhancers show reduced activity of TFs. We used single molecule footprinting (SMF) to measure the co-occurrence of 5mC with chromatin accessibility (CA), nucleosome occupancy and TF binding on individual DNA molecules of mouse CREs. We found that for the vast majority of active enhancers tested, presence of 5mC on DNA does not antagonise the chromatin opening nor the binding of TFs on the same molecules. Yet, we identify a subset of methylation-sensitive CREs in mESCs as well as three somatic lineages, where TF binding preferentially occurs on the fraction of enhancer molecules that are devoid of 5mC \emph{in vivo}. Using genetic perturbation, we confirm that this molecular antagonism is the result of direct repression of TF binding by 5mC. Additionally, we show that the occupancy levels of TFs at these active enhancers is a direct function of 5mC removal by active demethylation. 


\begin{figure*}%[tbhp]
\centering

 \makebox[\textwidth]{\includegraphics[width=.8\paperwidth]{Figures/Figure_1}}
%\includegraphics[width=.8\linewidth]{Figures/Figure_1.png}
\caption{\textbf{DNA methylation is neutral to chromatin accessibility at the majority of \emph{cis}-regulatory elements.} \textbf{(A)} Single locus example of the heterogeneity in DNA methylation (5mC) levels at enhancers and promoters when resolved on individual molecules. Upper panel: Genome browser track displaying the chromatin marks and the average methylation as measured by Whole Genome Bisulfite Sequencing (WGBS) around the \textit{Macf1} promoter and its intragenic enhancer. Lower panel: At the molecular level, 5mC at enhancers is heterogeneous. 5mC of individual CpGs (red) is shown at molecular level (blue - methylated, white - unmethylated). \textbf{(B)} Schematic representation of the experimental strategy used to identify molecular antagonisms between 5mC and chromatin accessibility (CA). Single molecule footprinting (SMF) is performed using the methyltransferase M.CviPI to measure CA in the GpC dinucleotide context, that is distinct from the endogenous 5mC in CpG context. Bisulfite sequencing of the DNA provides continuous information on CA and 5mC over 300 bp long DNA molecules. The accessibility of each molecule is calculated using the methylation of GpCs within the 100 bp surrounding a CpG. The average accessibility is used to classify DNA molecules into accessible and inaccessible fractions. In both fractions, the average 5mC levels are calculated. White/blue lollipops represent the endogenous un-/methylated state, white/black lollipops represent the CA as measured by SMF. \textbf{(C)} Most intermediately methylated CpGs show no difference in 5mC between the two separated CA fractions. Violin bar plots showing the CA (top) and 5mC (bottom) at CpGs in bulk and in the two separated fractions (n = 97,808). Boxplots show median (black middle line), 25th and 75th percentiles (black boundaries). \textbf{(D)} Single locus example of the 5mC and CA of individual molecules at a neutral CpG. Upper panel: Genome browser track displaying the chromatin marks and the average methylation as measured by WGBS around an intergenic enhancer. Middle panel: Depicted is the average SMF signal (1 - methylation \%) of individual GpCs (black dots) and endogenous 5mC of individual CpGs (blue diamonds). Lower panels: Depicted are the single DNA molecules mapped to this genomic locus sorted by CA into an accessible (grey) and an inaccessible (black) fraction, and the 5mC of the molecules in both fractions. In the lower left panel, every column is a single GpC dinucleotide depicting its accessibility status (grey: accessible, black: inaccessible). In the lower right panel, every column is a single CpG dinucleotide depicting its methylation status (light blue: unmethylated, dark blue: methylated). Percentages represent average 5mC of the CpG of interest at the center in each fraction. The number of single DNA molecules used for the analysis is indicated.}
\label{fig:1}
\end{figure*}


\section*{Results}

\subsection*{Simultaneous quantification of 5mC and CA on individual DNA molecules}

Although 5mC can reduce the affinity of many TFs \emph{in vitro}, its functional impact on enhancer activity \emph{in vivo} remains unclear. Addressing this question using bulk assays is challenging, as 5mC is highly heterogeneous at enhancers in homogeneous cell populations \cite{hartl_cis-regulatory_2017, hodges_directional_2011, stadler_dna-binding_2011, ziller_charting_2013}. This epigenetic heterogeneity means that at steady state, the binding motifs of TFs contained within active enhancers are methylated in a fraction of the cells, while the same genetic sequence is found unmethylated in the rest of the population (exemplified in Figure \ref{fig:1}A). We postulate that if 5mC interferes with TF binding at enhancers, TFs should occupy more frequently the fraction of DNA molecules that are unmethylated. Thus, differential analysis of the TF binding activity as a function of the presence of 5mC could reveal if 5mC influences enhancer activity \emph{in vivo}, without requiring experimental perturbations. 
To address this question, we advanced Single Molecule Footprinting (SMF) \cite{kleinendorst_genome-wide_2021,krebs_genome-wide_2017,sonmezer_molecular_2021} to simultaneously quantify 5mC, CA and TF binding on individual DNA molecules genome-wide. We footprinted the genome of mESCs using M.CviPI, a methyltransferase that targets cytosines in the GpC dinucleotide context, which is distinct from the endogenously methylated CpG context (Figure \ref{fig:1}B). We then profiled methylation levels of all cytosines using bisulfite sequencing and analysed the CpG and GpC methylation patterns separately on individual DNA molecules (Figure \ref{fig:1}B). Since M.CviPI is only able to methylate accessible GpCs, TFs and nucleosomes leave characteristic footprints in the GpC methylation signal \cite{kelly_genome-wide_2012,nabilsi_multiplex_2014,sonmezer_molecular_2021}. To ensure that the M.CviPI treatment does not affect CpG methylation levels, we quantified its activity on naked DNA (Figure \ref{fig:S1}A). We observed that M.CviPI does not methylate CpGs with the exception of the previously reported CmCG context (Figure \ref{fig:S1}A) \cite{kelly_genome-wide_2012}. Thus, we excluded the CCG context from downstream analyses and confirmed that footprinting did not affect endogenous 5mC levels in CpG context (Figure \ref{fig:S1}B). To reach a coverage compatible with analysis at single molecule resolution (median coverage of ~100 molecules - Figure \ref{fig:S1}C), we used bait-capture against regulatory regions that cover a large fraction of mESC regulatory elements (59.7\%) \cite{sonmezer_molecular_2021}, leading to a highly reproducible measure of CA and accurate CpG methylation levels (Figure \ref{fig:S1}D-E). By measuring continuous accessibility levels over 100 bp, we were able to distinguish nucleosome-bound DNA molecules (inaccessible), from DNA molecules where chromatin is open (accessible), which is a proxy for TF binding (Figure \ref{fig:1}B). Comparison of the methylation status of CpGs on each group of molecules allowed us to ask if 5mC is associated with reduced regulatory activity of CREs (Figure \ref{fig:1}B). 

\subsection*{DNA methylation is not reduced on chromatin accessible molecules of most enhancers}

We postulated that if 5mC represses the activity of enhancers, its presence should be reduced at DNA molecules that are in an active state, where chromatin is accessible. To test this hypothesis, we focused on the 97,808 CpGs that are highly covered in our assay and that show heterogeneous methylation levels across the cell population (10-60\% average 5mC, >30x coverage, Figure \ref{fig:S1}F). Most of these intermediately methylated CpGs are intergenic and are enriched at active CREs such as enhancers (36\%), insulators (11\%) and promoters (8\%) (Figure \ref{fig:S1}G). For each locus, we compared the average methylation across the whole population with that of chromatin accessible or inaccessible DNA molecules (Figure \ref{fig:1}C). We observed that 5mC is not reduced from accessible molecules for most CpGs, showing comparable intermediate methylation level as the total population or the inaccessible DNA molecules. We observed a similar distribution when focusing only on CpGs within active enhancers (Figure \ref{fig:S1}H) suggesting that presence of 5mC is generally neutral to the CA status of enhancers. A single locus example of such neutral enhancers shows that 5mC is present in similar proportions on accessible molecules (28\%) and inaccessible molecules (27\%) (Figure \ref{fig:1}D). Together, this suggests that 5mC does not globally impact the CA of most enhancers. 

\subsection*{Genome-wide identification of molecular antagonism at enhancers}

While we did not observe a global effect of the presence of 5mC on CA at enhancers, we next asked if this effect could be restricted to a subset of enhancers. We tested if there are CpGs for which the proportion of molecules with 5mC is significantly lower when they are accessible. We could indeed identify 3,218 CpGs where 5mC was reproducibly lower on accessible molecules, resulting in a significant methylation difference between the two molecular fractions (Figure \ref{fig:2}A). This molecular antagonism between 5mC and CA was only observed for 3.3\% of all tested CpGs, suggesting that a defined subset of enhancers is potentially regulated by 5mC. An example antagonist CpG contained within an active enhancer illustrates how 5mC is reduced at accessible molecules (31\%), while inaccessible molecules have significantly higher methylation levels (67\%, Figure \ref{fig:2}B). 
We envision two possible models to explain this molecular antagonism. It could either result from a cell-to-cell variability in enhancer usage across the cell population, or from a differential usage of the two alleles within each cell. The latter is expected to be observed at Imprinting Control Regions (ICRs), where 5mC confers monoallelic gene expression at the loci \cite{ferguson-smith_discovery_2018}. At these regions, 5mC and CA are expected to occur on separate alleles and thus to be antagonist at the molecular level. Indeed, 2.6\% of the identified antagonist sites are located within previously annotated ICRs (Figure \ref{fig:S2}A), confirming the sensitivity of our approach. Thus, we wondered whether differential usage of alleles can further explain the heterogeneity observed at antagonist sites located outside of annotated ICRs. To test this, we generated SMF data from F1 hybrid mESCs (129/CAST), enabling us to discriminate the parental origin of homologous alleles using Single Nucleotide Polymorphisms (SNPs) to resolve 5mC and CA on individual alleles (see Methods). As expected, the paternal and maternal alleles are differentially methylated at ICRs (Figure \ref{fig:2}C and Figure \ref{fig:S2}B), and CA occurs primarily on the unmethylated allele (Figure \ref{fig:S2}B). In contrast, non-ICR antagonist sites have comparable intermediate 5mC levels and CA on both alleles (Figure \ref{fig:2}C) and the antagonism can be observed on both alleles (Figure \ref{fig:S2}C). This suggests that the molecular heterogeneity observed at antagonist sites reflect the heterogeneity of CRE usage within the cellular population, rather than a differential allelic usage in individual cells.

Analysis of the genomic distribution of the non-ICR antagonist sites revealed that a majority of them are located away from gene promoters, and are annotated as enhancers based on chromatin modifications (Figure \ref{fig:2}D, Figure \ref{fig:S2}D). Additionally, a fraction of these CpGs is found at insulator regions bound by CTCF (Figure \ref{fig:2}D and Figure \ref{fig:S2}D). This distribution is similar to that of neutral sites (Figure \ref{fig:2}D). Antagonist and neutral loci have comparably low CpG density (Figure \ref{fig:S2}E); are located within open chromatin regions in mESC (Figure \ref{fig:S2}F); and harbour comparably high hydroxy-methylation levels characteristic of active CREs (Figure \ref{fig:S2}G). One distinctive feature of CpGs with antagonist properties is that they tend to show higher levels of CA on average (Figure \ref{fig:S2}F) and are located close to DNase-seq peaks, where the highest level of TF binding is presumed to occur within a CRE (Figure \ref{fig:2}E). In summary, we identify a class of enhancers where presence of 5mC and CA are antagonist at the molecular level, suggesting a possible functional role for 5mC at these sites.

\begin{figure*}%[tbhp]
\centering
 \makebox[\textwidth]{\includegraphics[width=.7\paperwidth]{Figures/Figure_2}}
\caption{\textbf{Molecular antagonism between DNA methylation and CA at a subset of active enhancers.} \textbf{(A)} Molecular antagonism between DNA methylation (5mC) and chromatin accessibility (CA) occurs at a small subset of CpGs genome-wide. Volcano plot depicting the common odds ratio and the p-value of a Cochran-Mantel-Haenszel test that probes the methylation difference between the chromatin accessible and inaccessible fraction of molecules covering 97,808 individual intermediately methylated CpGs. For most CpGs, 5mC occurs at equivalent levels between chromatin accessible and inaccessible molecules (black dots), yet 3,218 CpGs show antagonist behavior (pink dots). The pie chart represents the relative proportion of neutral (black) and antagonist CpGs (pink) identified within the tested CpGs. The histogram on top shows the distribution of the common odds ratio. \textbf{(B)} Single locus example of the 5mC and CA of individual molecules at an antagonist CpG. Upper panel: Genome browser track displaying the chromatin marks and the average methylation as measured by WGBS around an intragenic antagonist \textit{Mcu} enhancer. Middle panel: Depicted is the average SMF signal (1 - methylation \%) of individual GpCs (black dots) and endogenous 5mC of individual CpGs (blue diamonds). Lower panels: Depicted are the single DNA molecules sorted by CA into an accessible (grey) and an inaccessible (black) fraction, and the 5mC of the molecules in both fractions. In the lower left panel, every column is a single GpC dinucleotide depicting its accessibility status (grey: accessible, black: inaccessible). In the lower right panel, every column is a single CpG dinucleotide depicting its methylation status (light blue: unmethylated, dark blue: methylated). Percentages represent average 5mC of the CpG of interest at the center in each fraction. The number of single DNA molecules used for the analysis is indicated. \textbf{(C)} Antagonist CpGs are not differentially methylated between alleles. Violin box plots showing the average 5mC distribution of antagonist sites that do not overlap with annotated Imprinting Control Regions (ICRs) (left), maternal ICRs (middle) and paternal ICRs (right). Average 5mC levels of individual CpGs were calculated by analysing both alleles together (both) or each allele of a F1 hybrid (129/CAST) mESC line individually. Boxplots show median (black middle line), 25th and 75th percentiles (black boundaries). Number of stars illustrates the significance of a Wilcoxon rank test when comparing the two alleles (ns: p > 0.01; *: p < 0.01; **: p < 0.001 ***: p < 0.0001 ****: p < 0.00001). \textbf{(D)} The majority of the antagonist CpGs lie within annotated enhancer sites. Stacked barplot showing the genomic context annotation for neutral and antagonist sites using publicly available chromHMM annotations clustered into 5 groups. ChromHMM data from Pintacuda \textit{et al}., 2017 \cite{pintacuda_hnrnpk_2017}. \textbf{(E)} Antagonist CpGs are frequently located at the center of active \emph{cis}-regulatory elements. Lower panel: Relative position of neutral (black) and antagonist CpGs (pink) to the highest accessibility nucleotides of the regulatory element (defined by the peak summit of DNase Hyper Sensitivity (DHS)). Upper panel: The average DHS signal for the same regions shown as reference. Data from Domcke \textit{et al}., 2015 \cite{domcke_competition_2015}.}
\label{fig:2}
\end{figure*}


\subsection*{5mC controls the regulatory activity of antagonist enhancers}

Establishing the regulatory function of 5mC at enhancers requires to move beyond the correlative evidence of their antagonism at the molecular level, for example by determining the consequence of perturbing 5mC levels on their regulatory activity. To test this, we used two complementary genetic perturbations that either remove or increase 5mC levels globally and compared the changes in CA at antagonist and neutral CREs. We generated SMF data in an isogenic cell line where 5mC was entirely removed by the deletion of all three DNMTs (DNMT TKO) \cite{domcke_competition_2015}. Here, we focused our analysis on active enhancer sites, ignoring for instance insulators (see Methods for details). Despite having comparable levels of starting 5mC in wild-type (WT) cells (Figure \ref{fig:S3}A), neutral and antagonist enhancers respond differently to 5mC depletion. While the fraction of accessible DNA molecules remained constant at neutral enhancers, antagonist enhancers showed a significant increase in their CA (Figure \ref{fig:3}A-C). For instance, the neutral \textit{Mpp7} enhancer that is methylated on 27\% of its molecules in WT cells has almost identical accessibility in the absence of 5mC (Figure \ref{fig:3}B). In contrast, the antagonist \textit{Mcu} enhancer that is methylated on 50\% of its DNA molecules in WT cells, gains CA in 29\% of the cells upon depletion of 5mC (Figure \ref{fig:3}C). Complementarily, we performed SMF in cells depleted for the three 5mC turnover enzymes (TETs, TET TKO)\cite{ginno_genome-scale_2020}, where 5mC is increased at many CREs (Figure \ref{fig:S3}B). We compared the CA changes at neutral and antagonist sites that gain 5mC upon TET depletion and observed a significant decrease in the proportion of accessible molecules at antagonist enhancers, while CA at neutral enhancers remained constant (Figure \ref{fig:3}D-F). In this case, the antagonist \textit{Mcu} enhancer, which gained methylation on 36\% of its molecules upon TET depletion, concomitantly lost CA in 19\% of the molecules (Figure \ref{fig:3}F). Interestingly, the loss of accessibility is proportional to the gain in 5mC in TET TKOs at antagonist sites (R = -0.369, Figure \ref{fig:S3}D), suggesting that CA may be directly controlled by the 5mC levels at these sites. We observed a similar correlation for the gain of CA in DNMT TKOs, albeit with lower amplitude of changes (R = 0.227, Figure \ref{fig:S3}C). Together these data show that 5mC controls the levels of CA at antagonist enhancers.

Establishing the direct impact of changes in enhancer activity on gene expression is limited by the fact that most genes are regulated by multiple enhancers, and that enhancer-gene mapping methods are prone to high error rates \cite{moore_curated_2020}. Yet, we could associate 104 antagonist enhancers with genes that are gaining expression in DNMT TKO, and 74 that were linked to genes that lose activity upon an increase of 5mC levels in TET TKO cells. We identified instances of dysregulated genes that were associated with multiple antagonist sites, suggesting a functional link between changes in methyl-dependant enhancer activity and gene expression (Figure \ref{fig:S3}E). We also observed examples of antagonist sites that associate with multiple genes that are dysregulated upon changes in 5mC (Figure \ref{fig:S3}F). For instance, \textit{Gli2} (GLI Family Zinc Finger 2) and \textit{Inhbb} (Inhibin Subunit Beta B) that are predicted to be regulated by an antagonist enhancer located at 114 kb and 255 kb from their promoters, respectively, both lose expression upon TET TKO (Figure \ref{fig:3}G). Additionally, we identified antagonist CpGs within the enhancer cluster regulating the core pluripotency factor \textit{Sox2} (two out of 20 analysed CpGs, Figure \ref{fig:S3}G). This is in agreement with previous observations that the activity of the enhancer cluster regulating \textit{Sox2} depends on the presence of 5mC at these loci \cite{song_dynamic_2019}. In summary, molecular antagonism between 5mC and CA identifies enhancers whose regulatory activity is controlled by presence of 5mC.

\begin{figure*}%[tbhp]
\centering
 \makebox[\textwidth]{\includegraphics[width=.7\paperwidth]{Figures/Figure_3}}
\caption{\textbf{DNA methylation represses the regulatory activity of antagonistic enhancers.} \textbf{(A)} Chromatin accessibility (CA) is increased upon depletion of DNA methylation (5mC) at antagonist enhancers. Violin box plots representing the changes in CA as measured by SMF upon removal of 5mC in mESC knocked out for the three endogenous DNMTs (DNMT TKO). Accessibility is significantly increased at antagonist sites (pink, n = 900) and ICRs (rose, n= 70) but not at neutral sites (black, stringent neutral sites, n = 7,286). Depicted is the distribution of the differences in average accessibility (GpC) at a 101 bp window of individual loci. Loci where starting CpG methylation is lower than 30\% in WT cells were excluded from the analysis, as well as sites annotated as insulator or inactive by chromHMM. Red line depicts the median of neutral sites. The number of stars illustrate the significance of a Wilcoxon rank test when compared to the neutral sites (ns: p > 0.05 *: p < 0.05; **: p < 0.01 ***: p < 0.001 ****: p < 0.0001). \textbf{(B-C)} Single locus example of the differences between WT and DNMT TKOs at (B) a neutral or (C) an antagonist site. Top panels display the location and methylation status of the CpGs in the two cell lines. Bottom panels depict the average SMF signal (1 - methylation \%) of individual GpCs in WT (black dots) and DNMT TKO mESCs (red dots). \textbf{(D)} CA is reduced upon increase of 5mC at antagonist enhancers. Violin box plots representing the changes in CA as measured by SMF upon increase of 5mC in mESC knocked out for the three TET enzymes (TET TKO) involved in 5mC turnover. Accessibility is significantly reduced at antagonist sites (pink, n = 516) and ICRs (rose, n= 22) but not at neutral sites (black, n = 2,455). Loci where CpG methylation increase is less than 30\% in TET TKO cells were excluded from the analysis, as well as sites annotated as insulator or inactive by chromHMM. Yellow line depicts the median of neutral sites. Same representation as Figure \ref{fig:3}A. \textbf{(E-F)} Single locus example of the differences in CA between WT and TET TKOs at (E) a neutral or (F) an antagonist site. Top panels display the location and methylation status of the CpGs in the two cell lines. Bottom panels depict the average SMF signal (1 - methylation \%) of individual GpCs in WT (black dots) and TET TKO mESCs (yellow dots). \textbf{(G)} Example of a locus where loss of CA at the \textit{Gli2/ Inhbb} antagonist enhancer is correlated with a decrease in gene expression at the connected genes upon 5mC increase. Top panel: Genome browser track displaying the chromatin marks and the average methylation around an intergenic antagonist enhancer (same representation as Figure \ref{fig:1}A). The enhancer was linked to the \textit{Gli2} and \textit{Inhbb} genes by GREAT (pink dotted lines). The fold change and adjusted p-value of the expression changes as measure by RNA-seq are displayed. Bottom panels: Accessibility changes at the \textit{Gli2/ Inhbb} antagonist enhancer in WT versus TET TKO. Same representation as Figure \ref{fig:3}F. RNA-seq data from Huang \textit{et al}., 2021 \cite{huang_chromosomal_2021}.}
\label{fig:3}
\end{figure*}

\subsection*{Cell-type specific activity of methyl-sensitive enhancers}

In mESCs, antagonist enhancers are found in a dual state, where a significant fraction of the DNA molecules are epigenetically repressed and transcriptionally inactive. Thus, these enhancers could be poised by 5mC in pluripotent cells, for rapid activation at later developmental stages. To test this, we probed the fate of antagonist enhancers identified in mESCs upon \emph{in vitro} differentiation into neural progenitor cells (NPs). Upon differentiation, we observed a strong increase in 5mC at antagonist sites (Figure \ref{fig:4}A), that is concomitant with a loss of CA (Figure \ref{fig:4}B). For example, at an antagonist enhancer that is methylated in 16\% of the mESCs with a CA of 59\%, methylation increases to 73\% in NPs and CA decreases to 30\% (Figure \ref{fig:4}C). In line with these observations, gene ontology (GO) terms analysis of the genes associated with antagonist enhancers showed an enrichment for ES specific processes (Figure \ref{fig:4}D). Together, this shows that most antagonist sites are bona fide mESC enhancers that get inactivated upon differentiation.

During early development, 5mC is dynamic with timed waves of global de- and re-methylation. When expanded \emph{in vitro} in serum/LIF conditions, the epigenetic state of pluripotent mESCs is representative of early embryos just after re-acquisition of 5mC \cite{habibi_whole-genome_2013}. We thus wondered if enhancer regulation by 5mC is specific to the pluripotent state. To explore this question, we generated additional SMF datasets in somatic cells representing the erythroid (murine erythroleukemia cells, MEL) and muscle (myoblast cells, C2C12) lineages. In each cell lineage, we could identify a small proportion of antagonist enhancers (1.9-3.9\%, Figure \ref{fig:4}E). The majority of the enhancers were only active in one of them, with only a subset shared between cell types (Figure \ref{fig:4}F). This is in agreement with the observation that the genes associated with those antagonist enhancers are enriched for GOs associated with the respective cellular identity (Figure \ref{fig:4}D, Figure \ref{fig:S4}A). Motif enrichment analysis revealed that cell-type specific antagonist loci are highly enriched for various E-box motifs, suggesting that the sensitivity to 5mC at these sites could be explained by the binding of basic helix-loop-helix (bHLH) TFs such as Max-Myc in mESCs or USF1 in erythrocytes (Figure \ref{fig:S4}B). Additionally, we also found that antagonist enhancers are enriched for motifs of TFs involved in the control of the cell identity in each cell type. These include for instance KLF5 in ESCs, Sox10 in NPs, GATA3/6 in MELs, and HIF2a and MyoD in C2C12s. We conclude that enhancer regulation by 5mC is not a specific feature of pluripotent cells, and that 5mC contributes to the regulation of specific sets of cell-type specific enhancers in multiple somatic lineages.

\begin{figure*}%[tbhp]
\centering
 \makebox[\textwidth]{\includegraphics[width=.7\paperwidth]{Figures/Figure_4}}
\caption{\textbf{Cell-type specific activity of antagonist enhancers.} \textbf{(A-B)} Antagonist sites identified in embryonic stem cells (ES) show increase in DNA methylation (5mC) and loss of chromatin accessibility (CA) upon differentiation into neural progenitors (NP). Violin box plots representing the average (A) 5mC and (B) CA measured by SMF at antagonist sites identified in mESC. The number of stars illustrate the significance of a Wilcoxon rank test (ns: p > 0.05 *: p <= 0.05; **: p <= 0.01 ***: p <= 0.001 ****: p <= 0.0001). \textbf{(C)} Single locus example of the fate of an antagonist enhancer identified in mESC during neural differentiation. The top panel shows the methylation of the CpGs in the two cell lines. Bottom panel depicts the average SMF signal (1 – methylation \%) of individual GpCs in ES (black dots) and NP (grey dots). \textbf{(D)} ES antagonist enhancers are connected to ES specific genes. Dot plots showing the adjusted p-value of the 7 top GO term processes (fold change > 2) identified by GREAT analysis of ES specific antagonist sites. Black triangles indicate ES specific processes. \textbf{(E)} Similar proportions of antagonist CpGs are identified in stem cells and somatic cell lines. Pie charts represent the relative proportion of neutral (black) and antagonist CpGs (pink) identified in embryonic stem cells (ES), erythrocytes (murine erythroleukemia cells, MEL), neural progenitors (NP) and myoblasts (C2C12). \textbf{(F)} Antagonism is cell-type specific at most CpGs. Heatmap showing the -log of the methylation difference between the accessible and inaccessible molecules (measured as common odds ratio by the Cochran-Mantel-Haenszel test) for antagonist CpGs covered in all four cell lines (n = 1,671). Antagonist sites were clustered by k-means (k = 5). }
\label{fig:4}
\end{figure*}

\subsection*{Active 5mC turnover is required for TF binding at antagonist loci}

We next thought to investigate the molecular mechanism behind the observed methyl-sensitivity of antagonist enhancers. Based on the knowledge that some TFs can be sensitive to the presence of 5mC, we hypothesized that antagonist sites are regulated by methyl-sensitive TFs that preferentially bind to their recognition motif in absence of 5mC. To test this hypothesis, we leveraged the ability of SMF to resolve TF occupancy at molecular resolution \cite{kleinendorst_genome-wide_2021, sonmezer_molecular_2021} and quantified 5mC simultaneously on the same DNA molecules (see Methods). We scanned intermediately methylated CREs across the genome for TF footprints (see Methods), and compared the 5mC levels of TF bound molecules with those occupied by nucleosomes at the same locus. We postulated that 5mC levels should be reduced from the fraction of DNA molecules bound by those TFs that are methyl-sensitive (Figure \ref{fig:5}A). From this analysis, we identified 156 high confidence antagonist TF binding sites (TFBS) for which TF bound molecules had significantly reduced 5mC levels, representing 37\% of all analysed TFBS (Figure \ref{fig:5}B). For instance, we could identify a CTCF binding site where 54\% of the nucleosome-occupied molecules are methylated, while only 7\% of those bearing a TF footprint were methylated at the same locus (Figure \ref{fig:5}C). This is in contrast to the TF bound molecules at neutral CTCF sites, that show comparable 5mC levels on the TF bound and the nucleosome occupied molecules (Figure \ref{fig:S5}A). This enrichment of CTCF binding over unmethylated DNA molecules is supported by the reanalysis of data obtained by CTCF ChIP coupled with bisulfite sequencing (ChIP-bis) \cite{feldmann_transcription_2013} (Figure \ref{fig:S5}B), confirming the accuracy of the identification of methyl-sensitive TFBS by SMF \emph{in vivo}. 


\begin{figure*}%[tbhp]
  \captionsetup{labelformat=empty}
  \centering
  \makebox[\textwidth]{\includegraphics[width=.8\paperwidth]{Figures/Figure_5}}
  \caption{}
\end{figure*}
\begin{figure*}
  \captionsetup{labelformat=adja-page}
  \ContinuedFloat
  \caption{\textbf{Direct control of TF binding occupancy by DNA methylation at enhancers.} \textbf{(A)} Schematic representation of the experimental strategy used to identify antagonist TFBS. Short footprints created by TFs were distinguished from large nucleosome footprints by measuring the accessibility of a bin at the TFBS and two bins surrounding it. DNA molecules were classified into TF bound (purple bar), accessible (green bar) and nucleosome bound (black bar) fractions. The CpG methylation (5mC) of the TF bound and nucleosome occupied fractions are then compared to test for differential methylation. White/blue lollipops represent the endogenous un-/methylated state, white/black lollipops represent the chromatin accessibility as measured by SMF. \textbf{(B)} Molecular antagonism between 5mC and TF binding occurs at a subset of TFBS genome-wide. Volcano plot depicting the common odds ratio and p-value of a Cochran-Mantel-Haenszel test that probes the methylation difference between the TF bound and nucleosome bound fraction. For most TFBS, 5mC occurs at equivalent levels between TF bound and nucleosome bound molecules (black dots), yet 156 TFBS show antagonist behavior (pink dots). The representative motif names are shown for the top antagonist sites. The pie chart represents the relative proportion of neutral (black) and antagonist (pink) TFBS. Only TFBS with >5\% TF binding frequency were considered. \textbf{(C)} Single locus example of the 5mC difference between TF bound and nucleosome bound molecules at an antagonist CTCF site. Top panel depicts average SMF signal (1 - methylation \%) of individual GpCs (black dots) and average 5mC of individual CpGs (blue diamonds) at a CTCF binding site (grey box). Lower panels show single DNA molecules stack sorted into a TF bound (purple), accessible (green) and nucleosome bound (black) fraction. In the lower left panel, every column is a single GpC dinucleotide depicting its accessibility status (grey: accessible, black: inaccessible). In the lower right panel, every column is a single CpG dinucleotide depicting its methylation status (light blue: unmethylated, dark blue: methylated). Percentages represent average 5mC of the CpG in each fraction. The number of single DNA molecules used for the analysis is indicated. \textbf{(D)} Bar chart representing the percentage of TFBS classified as antagonist for TF motifs where >10 binding sites are covered by SMF. Depicted is the percentage of binding sites lying within unmethylated regions (<10\% 5mC, grey), neutral* (black) and antagonist* (pink). \textbf{(E)} Antagonist TFs lose binding upon increase in 5mC in TET TKOs. Violin box plots representing the changes in states frequency upon increase of 5mC in TET TKO. The TF bound fraction is significantly reduced at antagonist sites (n = 29) but not at neutral sites (n = 17), while the nucleosome bound fraction is increased upon KO. Loci where 5mC increase in TET TKO cells is less than 30\% were excluded from the analysis. The number of stars illustrate the significance of a Wilcoxon rank test when compared to the neutral sites (ns: p > 0.05 *: p <= 0.05; **: p <= 0.01 ***: p <= 0.001 ****: p <= 0.0001). \textbf{(F)} TF occupancy is a function of 5mC levels at antagonist sites. Scatterplot comparing the changes in TF binding and 5mC in TET TKO. The decrease in TF occupancy is anti-correlated with the increase in 5mC at the locus (n = 173). The regression line and Pearson correlation coefficients are displayed. A large fraction of Max-Myc binding sites is directly controlled by 5mC (pink dots, n = 65). \textbf{(G)} Single locus example illustrating the loss of TF binding upon 5mC increase at an antagonist Max-Myc binding site. Upper panel: Genome browser track displaying the chromatin marks and the average methylation (WGBS) around an intragenic antagonist \textit{Tfdp1} enhancer with a Max-Myc binding site. Middle panel: On top, the average 5mC of the CpGs in the two cell lines is shown. Below, the average SMF signal (1 - methylation \%) of individual GpCs in WT (black dots) and TET TKO (yellow dots) at a single locus containing a Max-Myc binding site (grey box) is displayed. Lower panels: Single molecules stacks for WT (lower left panels) and TET TKO (lower right panels) as described in Figure \ref{fig:5}C. \textbf{(H)} Mechanistic model describing the effects of 5mC on enhancers activity. 5mC does not affect chromatin accessibility nor TF binding at a majority of active enhancers (left), while it controls the activity of a subset of cell-type specific enhancers (right). Some antagonist enhancers regulate critical cell identity genes and active 5mC turnover by TET enzymes is required for TF binding at these sites. The balance between demethylation by TETs and \emph{de novo} methylation by DNMTs controls the activity of these enhancers. Neutral* and antagonist* sites in this figure are defined with soft criteria using only the common odds ratio of the Cochran-Mantel-Haenszel test.}
\label{fig:5}
\end{figure*}

Most of the identified antagonist binding sites are bound by known methyl-sensitive TFs such as CTCF, NRF1 or Max-Myc. Yet, we noted that the distribution of average 5mC at the regions bound by these TFs, as well as the proportion of antagonist binding sites within their binding events, varies considerably between TFs (Figure \ref{fig:5}D). For instance, NRF1 is almost exclusively bound at unmethylated loci, while CTCF and Max-Myc frequently occupy regions with intermediate methylation (Figure \ref{fig:5}D, Figure \ref{fig:S5}C). At those methylated sites, we detect a strong antagonism for most of the Max-Myc binding events (63\%, Figure \ref{fig:5}D, Figure \ref{fig:S5}D). In contrast, depending on the locus considered, 5mC is either neutral or antagonist to CTCF binding (Figure \ref{fig:5}D, Figure \ref{fig:S5}D). This differential behaviour is unlikely to be explained by the binding of different TFs at the motifs as we detected comparably high ChIP-seq enrichments at neutral and antagonist binding sites for CTCF or Myc (Figure \ref{fig:S5}E-F). In fact, for both TFs, we observed an anti-correlation between TF occupancy and the 5mC level at their binding sites at antagonist, but not neutral sites (Figure \ref{fig:S5}G-H). This suggests that TF occupancy is directly regulated by DNA methylation at antagonist binding sites.

It has been previously observed that the methyl-sensitivity of CTCF depends on the specific position of CpGs in its recognition motif \emph{in vivo} \cite{hashimoto_structural_2017,maurano_role_2015,wang_widespread_2012}. We thus compared the binding motifs for CTCF between antagonist and neutral sites, and observed that antagonism is associated with a frequent occurrence of CpGs at positions \#7 of the motif (Figure \ref{fig:S5}I). This suggests that modulation of CTCF binding levels by 5mC depends in part on the sequence of its binding motif. Max-Myc recognizes a short and rather constrained E-box motif that contains a CpG at its central position. In contrast to CTCF, we did not detect specific sequence features distinguishing antagonist from neutral binding sites (Figure \ref{fig:S5}J), suggesting that other mechanisms are required to explain its binding at methylated loci. 

We next explored the hypothesis that the binding of methyl-sensitive TFs at those sites could be mediated by TET-induced demethylation. To test the dependency of antagonist binding events on active demethylation, we measured the changes in TF binding frequency in WT versus TET TKO cells. We observed a global loss of TF binding at antagonist sites in TET TKO cells (Figure \ref{fig:5}E, purple), often accompanied by a significant increase in nucleosome occupancy at the region (Figure \ref{fig:5}E, black). The changes in TF binding frequencies at antagonist sites are a direct function of the methylation gain (Figure \ref{fig:5}F, Figure \ref{fig:S5}K-L). In turn, 5mC explains 28\% of the variance in TF binding at these sites, suggesting a direct dependency of TF binding on 5mC at these regions. This effect is pervasive at Max-Myc bound regions, where occupancy levels appear to be largely controlled by 5mC, and its active removal by TET enzymes (Figure \ref{fig:5}F). This is for example evident at an example Max-Myc bound locus where in WT mESCs all TF bound molecules are unmethylated (0\%), despite an average methylation of 37\% at the locus (Figure \ref{fig:5}G, left panel). This selective binding of a small proportion of unmethylated molecules suggests that Max-Myc is only capable of binding this target motif when unmethylated. Upon TET TKO, 5mC is drastically increased to 85\% at the central CpG, leading to a complete loss in TF binding (Figure \ref{fig:5}G, right panel), revealing the dependency of Max-Myc on active DNA demethylation for its binding at this locus. Together, this suggests that 5mC controls the occupancy levels of Max-Myc at enhancers. Moreover, it suggests a model in which Max-Myc occupancy depends on the equilibrium between \emph{de novo} methylation by DNMTs and active DNA demethylation by TET enzymes (Figure \ref{fig:5}H).


\section*{Discussion}

A topical question in the epigenetics field is to establish the functional impact of chromatin and DNA modifications on the activity of CREs and transcription activation. Enhancer activation is tightly coupled with a reduction of their 5mC, making them the most dynamic differentially methylated regions across tissues and cell types, and suggesting that epigenome remodelling may be required for their activation. However, so far it could not be determined whether 5mC generally contributes to the regulation of enhancer activity \cite{luo_dynamic_2018,mattei_dna_2022}. By quantifying the molecular co-occurrence of 5mC with CA and TF occupancy across the genome using a single molecule multi-omics approach, we were able to systematically determine the sensitivity of enhancers to 5mC and estimate the proportion of methyl-sensitive enhancers genome-wide.

Our data argue that 5mC is instructive for TF binding and gene regulation at a limited subset of enhancers, while for most of them, changes in methylation levels are a consequence of TF binding activity and chromatin opening of the locus. This is an important consideration when interpreting changes in 5mC during development \cite{iurlaro_dna_2017} or cell-to-cell epigenetic variation data \cite{farlik_single-cell_2015,smallwood_single-cell_2014}, as most variations in 5mC are unlikely to drive changes in gene regulation. In general, it appears that most TFs can bind in presence of 5mC on the DNA \emph{in vivo}. This extends previous observations that 5mC can remain at enhancers after their chromatin has been opened \cite{barnett_atac-me_2020}, bound by TFs \cite{donaghey_genetic_2018, feldmann_transcription_2013} and marked by active transcription marks such as H3K27ac \cite{charlet_bivalent_2016}.

Most of the existing knowledge on the effects of 5mC on TF occupancy arose from experiments that identified newly gained TF binding events upon global perturbation of 5mC levels \cite{cusack_distinct_2020,domcke_competition_2015,maurano_role_2015,yin_impact_2017}. Depletion of 5mC is lethal in almost all somatic cell lines and tissues, preventing the study of the majority of enhancers that are not active in ESCs. SMF overcomes these limitations and identifies methyl-sensitive regions in absence of experimental perturbations making it applicable to any cell-type or tissue. Moreover, the methyl-sensitive binding sites previously identified upon depletion of 5mC are often occurring outside of active regulatory elements illustrating how 5mC represses cryptic binding sites that are normally not occupied by TFs \cite{cusack_distinct_2020,domcke_competition_2015,maurano_role_2015}. Here, the single molecule resolution of our analysis unravels how the presence of 5mC affects the binding of TFs at active regulatory elements such as enhancers. The ability of SMF to detect methyl-sensitive enhancers in multiple cell lineages revealed that 5mC and 5mC turnover may play a role in the establishment of cell-type specific gene expression profiles.

Analysis of TF binding in relation to 5mC at molecular level shows that the occupancy levels of TFs such as the general transcription activator Max-Myc are regulated by 5mC at many enhancers. Thus, in addition to repressing cryptic binding sites, we show that 5mC plays a role in controlling the binding intensity of TFs at enhancers. Max-Myc was proposed to act as a transcriptional amplifier that broadly enhances the transcription levels of active genes \cite{lin_transcriptional_2012,nie_c-myc_2012}, promoting cell proliferation in various cellular contexts \cite{kress_myc_2015}. Our finding suggests that 5mC could participate in this global control of transcription levels through the tuning of Max-Myc occupancy levels. It will be interesting to evaluate this functional link between 5mC levels and driven proliferation in the biological contexts where global reduction of 5mC has been described. This encompasses the timed acquisition of 5mC during development \cite{iurlaro_dna_2017}, the oscillations of 5mC levels in stem cell populations \cite{rulands_genome-scale_2018}, or the global hypomethylation observed in many cancers \cite{baylin_decade_2011}.


\section*{Methods}

\subsection*{Experimental model and subject details}
Mouse ES cells (129 WT \cite{stadler_dna-binding_2011}, DNMT TKO \cite{domcke_competition_2015}, TET TKO \cite{ginno_genome-scale_2020} and F1 hybrid cells (129/CAST) \cite{giorgetti_structural_2016}) were cultured on 0.2\% gelatin-coated plates in ES medium (Dulbecco’s Modified Eagle Medium (DMEM), supplemented with 15\% Fetal Bovine Serum (FBS), Leukemia Inhibitory Factor (LIF), 2-Mercaptoethanol, 2 mM L-Glutamine and 1x non-essential amino acids) at 37°C and 5\% CO2. Medium was changed daily and cells were split every second day. The myoblast (C2C12) cells were grown on collagen coated plates in low glucose DMEM medium/F10 nutrient mix supplemented with 20\% FBS and 2 mM L-glutamine. The murine erythroleukemia (MEL) cells, obtained from DSMZ (ACC 501), were grown in DMEM without Sodium Pyruvate media supplemented with 20\% FBS and 2 mM L-glutamine. ES to NP differentiation was performed as previously described \cite{bibel_generation_2007}. 

\subsection*{Single Molecule Footprinting}
Single Molecule Footprinting with targeted enrichment (SMF) was performed as previously described \cite{kleinendorst_genome-wide_2021,sonmezer_molecular_2021}. In short, cultured cells were harvested using trypsin and washed twice with 1x PBS. Cells were counted and 250,000 cells were used per reaction. Cell pellets were resuspended in ice-cold lysis buffer, incubated on ice for 10 min and spun at 1,000x g at 4°C for 5 min. Nuclei were resuspended in 1x M.GpC buffer (NEB, \#M0227L). For the GpC methyltransferase treatment, freshly made GpC methyltransferase mix (1x M.GpC buffer, 300 mM sucrose, 64 μM SAM (NEB, \#B9003S)) and 200 U M.CviPI (NEB, \#M0227L) were added and incubated at 37°C for 7.5 min. For a second incubation round at 37°C for 7.5 min the reaction was replenished with 100U of M.CviPI and 128 pmol of SAM. Prewarmed stop solution and proteinase K were added and incubated overnight at 55°C. The next day, DNA was extracted using phenol-chloroform and treated with RNase A at 37°C for 30 min. For making the Whole Genome Bisulfite Sequencing (WGBS) library with targeted enrichment, DNA was fragmented into 300 bp fragments via sonication using Covaris model S2. For library preparation and targeted enrichment of \emph{cis}-regulatory elements, the SureSelect XT Mouse Methyl-Seq Kit Enrichment System for Illumina Multiplexed Sequencing Library protocol (Agilent Technologies, version E0, April 2018) was used as described before \cite{kleinendorst_genome-wide_2021}. Bisulfite conversion was performed using the ZYMO EZ DNA Methylation-Gold Kit (Zymo, \#D5005) according to the manufacturer’s protocol. The bisulfite-converted library was PCR amplified and indexed using the SureSelect XT Mouse Methyl-Seq Kit (Agilent, \#G9651A). Prepared libraries were run on an Illumina sequencing platform using a NextSeq High 150 bp paired-end mode. 

\subsection*{Computational procedures}
\subsubsection*{Sequencing data pre-processing}
SMF data was processed as previously described \cite{kleinendorst_genome-wide_2021,sonmezer_molecular_2021}. In short, Illumina adaptors and low quality bases were trimmed and low quality reads were removed using Trimmomatic (0.36) \cite{bolger_trimmomatic_2014}. Pre-processed reads were mapped to the bisulfite indexed \textit{Mus musculus} reference genome (BSgenome.Mmusculus.UCSC.mm10) using the R package QuasR \cite{gaidatzis_quasr_2015}, which uses Bowtie \cite{langmead_ultrafast_2009} as an aligner, with specific alignment parameters (alignmentParameter = -e 70 -X 1000 -k 2 --best –strata) and keeping only uniquely aligned reads. Duplicated reads were removed using the tool MarkDuplicates from Picard (2.15.0) \cite{Picard2019toolkit}.

SMF F1 sequencing reads were trimmed for Illumina adaptors using Trim Galore! (v0.6.3) \cite{krueger_felixkruegertrimgalore_2021} and competitively aligned to two versions of the \textit{Mus musculus} reference genome (BSgenome.Mmusculus.UCSC.mm10) injected with the SNPs \cite{keane_mouse_2011} for the species of interest using QuasR \cite{gaidatzis_quasr_2015}. Finally, duplicated reads were removed using the tool MarkDuplicates from Picard (2.15.0) \cite{Picard2019toolkit}.

WGBS and CTCF ChIP-bis datasets were pre-processed and aligned as SMF data. 

DNase-seq data was also pre-processed with Trimmomatic \cite{bolger_trimmomatic_2014} (0.36) and then aligned to the \textit{Mus musculus} genome (BSgenome.Mmusculus.UCSC.mm10) using QuasR \cite{gaidatzis_quasr_2015}, keeping only uniquely aligned reads. Duplicated reads were removed using the tool MarkDuplicates from Picard (2.15.0) \cite{Picard2019toolkit}.

ChIP-seq data was pre-processed using the Galaxy platform \cite{afgan_galaxy_2018} using Trim Galore! \cite{krueger_felixkruegertrimgalore_2021} (Galaxy Tool version 0.4.3.1) to trim adaptor sequences and low quality bases, and aligned with Bowtie \cite{langmead_ultrafast_2009} (Galaxy Tool version 1.2.0; default parameters but for paired-end; for single-end parameters --best --strata --maxbts 800 -m 1 were used) to the \textit{Mus musculus} reference genome (USCS mm10), keeping only uniquely aligned reads (Galaxy Tool "Filter SAM or BAM, output SAM or BAM" 1.1.2 with filters on MAPQ>10 and bitwise filters to remove unmapped, not primary and supplementary alignments). Duplicated reads were removed using Picard \cite{Picard2019toolkit} MarkDuplicates (Galaxy Tool version Galaxy Version 2.7.1.1). Finally, normalized (reads per genome coverage) signal coverage bigwig files were obtained with deepTools \cite{ramirez_deeptools2_2016} bamCoverage (Galaxy tool version 3.0.2.0) and a bin size of 50 bases.

RNA sequencing data was processed using the nf-core/rnaseq pipeline (v3.4) \cite{patel_nf-corernaseq_2022} distributed through nf-core \cite{ewels_nf-core_2020}. The pipeline was launched with default parameters and each sample was annotated for strandedness.

\subsubsection*{SMF – Single molecule DNA methylation call}
Single molecule methylation state calling for all cytosines with at least 10-fold coverage was performed using QuasR \cite{gaidatzis_quasr_2015} as previously described \cite{kleinendorst_genome-wide_2021, sonmezer_molecular_2021}. Only CpGs in NWCGW (N = any base, W = no C or G) and GpCs in DGCHN (D = no C, H = no G, N = any base) context were considered to avoid interferences between CpG and GpC methylation calling (Fig. S1A). 

\subsubsection*{SMF – Quantification of chromatin accessibility and 5mC at individual DNA molecules}
The analysis was focused on CpGs which have heterogeneous methylation across cells by selecting those with intermediate methylation (10-60\%). The chromatin accessibility of each molecule covering those CpGs was calculated using the mean GpC methylation within a 101 bp bin surrounding it. This mean accessibility value was then used to classify each molecule into an accessible (>=50\% methylated) or inaccessible (<50\% methylated) fraction. 
To test for the significance of the difference in DNA methylation between the two fractions of molecules, a Cochran-Mantel-Haenszel test was performed including a pseudo-count of 1 and applying a continuity correction. We only considered CpGs covered in at least 2 out of 2 or 3 biological replicates and had a 30-fold coverage. The resulting common odds ratio and p-value were used to discriminate antagonist and neutral sites. Antagonist sites were defined with a common odds ratio of <= 0.5 and a p-value of < 0.05. All other sites were defined as neutral. Consequently, neutral sites included sites with high methylation difference but no significant p-value due to variability between the replicates or low coverage. Therefore, when comparing antagonist (high methylation difference) and neutral (no significant methylation difference) sites in subsequent analyses (Figs. 2D, E, 3A, B, D, E, Figs. S2E, F, G, 3A), higher stringency neutral sites (neutral+) were defined by further selecting for a common odds ratio between 0.9 and 1.1. 

\subsubsection*{SMF – Chromatin accessibility, DNA methylation, fraction size and fraction methylation}
For the mean chromatin accessibility (GpC methylation), first the mean methylation of all GpCs within the 101 bp bin surrounding a CpG was calculated for each biological replicate and then of this the weighted mean of all replicates. For the mean DNA methylation (CpG methylation) the weighted mean of all replicates was calculated. Coverage was used as weighing value. A coverage cutoff of 10 reads per bin was applied to ensure reliable methylation calling.

\subsubsection*{SMF – Single locus plots}
For plotting SMF data for single loci, functions of the R package SingleMoleculeFootprinting \cite{kleinendorst_genome-wide_2021} were adapted to plot CpGs and GpCs individually and to include the chromatin accessibility state sorting.

\subsubsection*{Defining chromatin states with chromHMM}
To annotate the chromatin states of antagonist and neutral sites, previously defined ChromHMM states for mouse embryonic stem cells for USCS mm10 (\url{https://github.com/guifengwei/ChromHMM_mESC_mm10}) \cite{pintacuda_hnrnpk_2017} were used that were in addition manually grouped into five ChromHMM clusters: C1 – CTCF (1\_Insulator), C2 – Inactive (2\_Intergenic, 3\_Heterochromatin, 5\_RepressedChromatin, 6\_BivalentChromatin, 12\_LowSignal/RepetitiveElements), C3 – Transcription (9\_TranscriptionTransition, 10\_TranscriptionElongation), C4 – CREs promoter (7\_ActivePromoter), C5 – CREs enhancer (4\_Enhancer, 8\_StrongEnhancer, 11\_WeakEnhancer). 

\subsubsection*{Genome browser tracks}
Bigwig files of chromatin modifications (H3K27me3, H3K27ac, H3K4me1, H3K4me3, DNase-seq, WGBS) and RNA-seq data of publicly available datasets were loaded and the signal scores at the given genomic locus window plotted as area plots or point plots (for WGBS and SMF) using the R package ggplot2 \cite{wickham_ggplot2_2016} using different degrees of smoothing depending on the window size. Gene promoter and exons for the USCS mm10 reference genome were identified using the R package TxDb.Mmusculus.UCSC.mm10.knownGene \cite{team_bc_txdbmmusculusucscmm10knowngene_2019}.

\subsubsection*{Allelic differences between antagonist and ICR sites}
Imprinting Control Regions (ICRs) were defined by manually curating previous annotations \cite{xie_base-resolution_2012}. Average DNA methylation of all antagonist sites was calculated for both alleles together or separately. In addition, antagonist sites were checked for C to T SNPs to avoid wrong methylation measurements. Average DNA methylation of all ES non-ICR antagonist sites, and all maternal and paternal ICR-annotated antagonist sites was plotted as violin box plots using ggplot2 \cite{wickham_ggplot2_2016} and an unpaired Wilcoxon rank test between the two separated alleles was applied using the stats\_compare\_means function of the ggpubr package with the maternal allele as reference group (Fig. 2C).

\subsubsection*{DHS peak analysis}
Published DHS peaks were used to calculate the distance of CpGs classified as neutral+ or antagonist to the summit of the closest DHS peak and plotted as a density plot using ggplot2 \cite{wickham_ggplot2_2016}. Only those CpGs were considered that are annotated as enhancers by chromHMM (C5 chromHMM cluster) and that have a DHS peak within 500 bp (n antagonist = 1,336; n neutral+ = 6,735). Furthermore, the read density of the DHS data was calculated at a 1 kb window around the DHS peak summits with the function qProfile from the QuasR \cite{gaidatzis_quasr_2015} package (Fig. 2E). 

\subsubsection*{Chromatin modifications}
Bigwig files of chromatin modifications (H3K27me3, H3K27ac, H3K4me1, H3K4me3, CTCF, DNase-seq) of publicly available datasets were loaded and the signal scores at 4 kb surrounding antagonist CpGs plotted using the R package EnrichedHeatmap \cite{gu_enrichedheatmap_2018} using 50 bp windows and smoothing (Fig. S2D).

\subsubsection*{CpG density}
CpGs were counted in a 400 bp window around all CpGs. The number of CpG per 100 bp was plotted as CpG density for all ES antagonist, neutral+ and ICR sites as violin box plots using ggplot2 \cite{wickham_ggplot2_2016} (Fig. S2E). 

\subsubsection*{hMeDIP-seq analysis – 5-hydroxymethylation (5hmC)}
Reads of hMeDIP-seq data (data from \cite{xu_genome-wide_2011} at a 500 bp window surrounding all NWCGWs were counted using the qCount function from the R package QuasR \cite{gaidatzis_quasr_2015}. Read counts were normalized by the total amounts of reads and a pseudo-count of 5 was added. The log2 of this counts per million (cpm+pc) was plotted for all CpGs and only antagonist or neutral+ sites as violin box plots using ggplot2 \cite{wickham_ggplot2_2016} (Fig. S2G).

\subsubsection*{Change in chromatin accessibility in perturbation assays}
Delta chromatin accessibility (CA) between wild type (WT) and triple-knockout (TKO) ES cell lines was calculated by subtraction of the CA in TKO from the CA in WT for CpGs identified as neutral+, antagonist and ICR in WT ES cells. To focus on \emph{cis}-regulatory elements, sites annotated as insulator (chromHMM cluster C1) or inactive (chromHMM cluster C2) were removed. To focus on sites where a potential change in CA could be correlated with changes in DNA methylation, CpGs were filtered for their DNA methylation change. For DNMT TKOs, only those CpGs were considered that had a starting DNA methylation of >30\% in WT cells, resulting in a loss of DNA methylation of >30\% in DNMT TKOs. For TET TKOs, only those CpGs were considered that showed an increase in DNA methylation of >30\% from WT to TET TKOs. Delta CA was plotted as violin box plots using ggplot2 \cite{wickham_ggplot2_2016} and an unpaired Wilcoxon rank test was applied using the stats\_compare\_means function of the ggpubr package with neutral+ sites as reference group (Fig. 3A, D). Note that the changes in chromatin accessibility of all tested sites as a function of DNA methylation changes is displayed in Figs. S3C-D.

\subsubsection*{Enhancer-Gene associations and GO term analysis}
Genomic Regions Enrichment of Annotations Tool (GREAT, 4.0.4) \cite{mclean_great_2010} was used using the R package rGREAT \cite{gu_rgreat_2021} to perform enhancer-gene associations and GO term analysis. For this, antagonist sites of the cell lines ES, NP, MEL and C2C12 were used as test regions, whole genome as background region and USCS mm10 as reference genome. The model “Basal plus extension” was applied with 5 kb upstream and 1 kb downstream for proximal, and 1000 kb for distal. Curated regulatory domains were included. The GO Biological Process enrichment tables were extracted and filtered for a fold enrichment of the binominal test (Binom\_Fold\_Enrichment) of >= 2. The top 8 processes were selected for plotting, based on their Bonferroni corrected p-value of the binominal test (Binom\_Adjp\_BH). To focus on cell-type specific antagonist sites, only those sites where considered that were not annotated as antagonist in another cell line (ES cells were used as comparison for all somatic cell lines; NPs were used as comparison for ES cells). The enhancer-gene association from the GREAT gene association files were used for RNA-seq analysis of ES antagonist site associated genes.

\subsubsection*{RNA-seq analysis (DNMT TKO and TET TKO)}
Differential gene expression analysis was performed using DESeq2 (v1.30.1) \cite{love_moderated_2014} after filtering out genes with less than 10 counts across all samples. Genes associated with a log2 fold change greater, or smaller, than 2, or -2, and an adjusted p-value lower than 0.05 were considered differentially expressed for DNMT TKOs and TET TKOs, respectively. Enhancer-gene associations from the GREAT analysis were used to identify genes that are potentially regulated by antagonist sites. For this only those antagonist sites where considered that showed DNA methylation and chromatin accessibility changes in the TKO cell lines of at least 10\% and 5\%, respectively (Fig 3G, Figs. S3E-F).

\subsubsection*{Defining cell-type specificity of antagonist sites}
To define whether antagonist sites are cell-type specific, a k-means clustering of the -log common odds ratio from the Cochran-Mantel-Haenszel test of the antagonist sites of all four cell lines (ES, NP, MEL, C2C12) was applied with k = 5. Since the antagonist sites of one cell line show increased DNA methylation levels in the other cell lines (Fig. 4A), we used an average DNA methylation cutoff of 5-85\% for the Cochran-Mantel-Haenszel test. A heatmap of the k-means clustered common odds ratios was plotted using the function Heatmap of the R package ComplexHeatmap \cite{gu_complex_2016} (Fig. 4F). 

\subsubsection*{Motif enrichment analysis}
Motif enrichment analysis of cell-type specific antagonist enhancers was performed using Hypergeometric Optimization of Motif EnRichment (HOMER, v4.11) \cite{heinz_simple_2010} with a window size of 101 bp around the antagonist CpGs, USCS mm10 as reference genome, and using the repeat-masked sequence. Z-score of the -log p-value of known motifs was calculated for each group and top 20 motifs of each group were selected for plotting. Motifs were hierarchical clustered and the z-score of the -log p-value and the enrichment (“\% of Target Sequences with Motif”) were plotted (Fig. S4B). The same definition of cell-type specific antagonist sites was used as for the GREAT analysis.

\subsubsection*{SMF – Quantification of TF binding and 5mC on individual DNA molecules}
To identify transcription factor (TF) binding states, a 30 bp bin was created centered on a curated list of transcription factor binding sites (TFBS) from the JASPAR database (2018) \cite{khan_jaspar_2018} for \textit{Mus musculus} USCS mm10 reference genome, and two 10 bp bins 10 bp up- and downstream of the center bin. To ensure to have only single instances of each genomic location of a TFBS, the TFBS were annotated with a published list of TFBS motif clusters \cite{glaser_assessing_2021,khan_jaspar_2018} and redundant TFBS were removed. 
We only analysed TFBS that had a least one CpG within the 30 bp bin centered around the motif. The mean GpC methylation of each molecule was calculated in each of the three bins for every considered TFBS, which was then used to classify each molecule into a TF bound, nucleosome bound or accessible fraction and to define the TF binding frequency as previously described \cite{kleinendorst_genome-wide_2021, sonmezer_molecular_2021}. 
The analysis was focused on sites that are bound by the TF (>5\% of the molecules) and with average CpG methylation levels of 10-80\%. To test for a significant difference in the DNA methylation between the TF bound and nucleosome bound fraction, a Cochran-Mantel-Haenszel test was performed including a pseudo-count of 1 and applying a continuity correction. We only considered TFBS that were covered in at least 2 out of 3 biological replicates and had a 30-fold coverage. The common odds ratio and p-value of the Cochran-Mantel-Haenszel test were used to discriminate antagonist and neutral sites. Antagonist sites were defined with a common odds ratio of <= 0.5 and a p-value of < 0.05. All other sites were defined as neutral. For subsequent analyses (Fig. 5D-F and Fig. S5D-L), only the common odds ratio was used to define antagonist (antagonist*, common odds ratio < 0.5) and neutral sites (neutral*, common odds ratio > 0.5). 

\subsubsection*{SMF – state frequencies and fraction methylation, TFBS methylation}
Binding state frequencies were calculated as means of all replicates. The CpG methylation of each TFBS and of the binding state fractions of each TFBS were calculated as weighted means of all replicates using the coverage as a weighing value. 

\subsubsection*{Comparison CTCF ChIP-bis vs SMF}
For all CTCF sites covered by SMF, we separated the nucleosome bound fraction of the reads and the TF bound fraction of the reads. We collected the average DNA methylation in each of the fractions and calculated the difference between the fractions. For the same binding sites, we collected the average DNA methylation in the input fraction (WGBS \cite{stadler_dna-binding_2011}) and in the CTCF bound fraction (ChIP-bis data \cite{feldmann_transcription_2013}) and calculated the methylation difference between the fractions. We applied a minimal coverage of >9 reads to calculate the average methylation. The delta-delta plot shows the comparison of the methylation difference between the TF bound/nucleosome bound fractions as measured by SMF or ChIP-bis (Fig. S5B).

\subsubsection*{ChIP-seq analysis – CTCF and Max-Myc}
Myc ChIP-seq datasets \cite{das_distinct_2014, chronis_cooperative_2017, mor_neutralizing_2018} were merged. Reads of ChIP-seq data (CTCF ChIP data \cite{stadler_dna-binding_2011} and merged Myc ChIP-seq data) at a 500 bp window surrounding unique CTCF or Max-Myc binding sites were counted using the qCount function from the R package QuasR \cite{gaidatzis_quasr_2015}. Read counts were normalized by the total amounts of reads and the log2 of this counts per million (cpm) was plotted for all binding sites and only antagonist* or neutral* sites as violin box plots using ggplot2 \cite{wickham_ggplot2_2016} (Figs. S5G-H).

\subsubsection*{Motif analysis CTCF and Max-Myc}
For each unique TFBS covered by SMF and annotated as antagonist* or neutral*, we extracted the DNA sequences of the motifs. We then calculated the CpG frequency at each motif position. We compared the CpG frequency for all motifs for which there are at least 10 instance in both the antagonist and neutral category. To detect enrichment of CpGs at specific motif positions, we plotted the CpG frequency at each position for the antagonist* and the neutral* category (Figs. S5I-J).

\subsubsection*{TF – Change in fraction size at TFBS in perturbation assay}
The difference in binding state fraction size (log2 fold change) between wild type (WT) ES and TET triple-knockout (TET TKO) ES cell lines was calculated (using a pseudo-count of 0.001), for all TFBS identified as neutral* and antagonist* in WT ES cells. To focus on sites where a potential change in TF binding could be correlated with a change in DNA methylation, only those TFBS were considered that showed an increase in DNA methylation of >30\% from WT to TET TKOs. In Fig. 4E, the log2 fold change of all binding state fractions was plotted as violin box plots using ggplot2 \cite{wickham_ggplot2_2016} and an unpaired Wilcoxon rank test was applied using the stats\_compare\_means function of the ggpubr package with neutral* sites as reference group. In Fig. 4F, the log2 fold change of the TF bound fraction of the antagonist* sites is plotted as a function of the change in DNA methylation of the TFBS from WT to TET TKOs using ggplot2 \cite{wickham_ggplot2_2016}. Note that the changes in TF occupancy of all tested TFBS as a function of DNA methylation changes is displayed in Figs. 5E-F and Figs. S5K-L.

\subsubsection*{Statistical analysis and reproducibility}
For SMF, at least two (three for WT ES and DNMT TKO ES) replicates were performed for each cell line.
All statistical analyses were performed using R software and are described in the figure legends and methods section. 
Violin box plots (Figs. 1C, 2C, 3A, D, 4A-B, 5E, Figs. S1H, 2E-G, 3A, 5G-H) were plotted by the R package ggplot2 \cite{wickham_ggplot2_2016} with arguments outlier.alpha = 0 and coef = 0. The upper and lower boundaries of the box plot represent the 25th and the 75th percentiles. The central line represents the median. P-values for Figs. 2C, 3A, D, 4A-B, 5E were calculated by unpaired Wilcoxon rank test using the R package ggpubr.
Pearson correlation efficient in Figs. 5F and Figs. S3C-D, 5E-F, K-L was calculated using the R package corrr \cite{kuhn_corrr_2022} with the option pairwise.complete.obs, and a linear regression line was plotted with the R package ggplot2 \cite{wickham_ggplot2_2016} using method = “lm”.
The Cochran-Mantel-Haenszel test (Figs. 2A, 5B) was performed under the criteria that the region of interest was covered by at least two replicates with a mean coverage of 30 reads. A pseudo-count of 1 was added to all possible states and the test was performed with a continuity correction.
SMF single locus plots (Figs. 1D, 2B, 3B-C, E-F, H, 4C, 5C, G, Figs. S2B-C, 3G, 5A) were plotted with the SingleMoleculeFootprinting \cite{kleinendorst_genome-wide_2021} R package functions as basis adapted to the needs of this publication.

\vspace{\baselineskip}

\noindent\begin{contributions}
A.R.K designed the study. E.K. and A.R.K wrote the manuscript. E.K performed the experiments with support of R.K. E.K. analysed the data with support from G.B. and S.K for statistical analysis. A.R.K supervised conduction of the experiments and the data analysis. All authors discussed the results and commented on the manuscript.
\end{contributions}

\noindent\begin{acknowledgements}
The authors are grateful to the members of the Krebs laboratory and Jamie Hackett for helpful discussions; Tuncay Baubec, Eileen Furlong, Mathieu Boulard, Ralph Grand, Thomas Dahlet and Sam Corless for comments on the manuscript; Alexander Stark for feedback; Ângela Goncalves for suggestions on allele-specific analyses. The authors are grateful to GBCS and GeneCore at EMBL for support in sequencing data generation and management; Luca Giorgetti, Dirk Schübeler and Moritz Mall for sharing cell lines. The salary of G.B. is supported by the Deutsche Forschungsgemeinschaft (KR 5247/1-1). Research in the laboratory of A.R.K is supported by core funding of the European Molecular Biology Laboratory, Deutsche Forschungsgemeinschaft (KR 5247/1-1, KR 5247/1-2). 
\end{acknowledgements}


\noindent\begin{interests}
The authors declare no competing interests.
\end{interests}


\section*{References}
\bibliography{5mC_citation_library}


\onecolumn
\newpage

%%%%%%%%%%%%%%%%%%%%%%%%%%%%%
% Supplementary Information %
%%%%%%%%%%%%%%%%%%%%%%%%%%%%%
\captionsetup*{format=largeformat}
\section*{Supplement Figures}
\let\originalthefigure\thefigure

\makeatletter 
\renewcommand{\thefigure}{S\@arabic\c@figure}
\makeatother
\setcounter{figure}{0}

\begin{figure}[b]
\centering
 \makebox[\textwidth]{\includegraphics[width=.7\paperwidth]{Figures/Figure_S1}}
\caption{\textbf{(A)} Nucleotide contexts methylated by M.CviPI \emph{in vitro}. Barplot representing the average methylation levels of lambda DNA in various trinucleotide contexts upon \emph{in vitro} methylation with M.CviPI in saturating conditions. M.CviPI fully methylates GpCs regardless of the nucleotide context (green bars). It methylates CpGs in GCmG contexts and displays a low level of non-specific activity in CCmG context (<10\%) (red bars). Both of these contexts were excluded from our analysis, that only considered NWCGW and DGCHN to avoid technical interferences between the quantification of the DNA footprinting signal and the measure of endogenous DNA methylation (5mC) levels. The CpGs in other contexts (blue bars) show levels comparable to those observed at non targeted cytosines representing technical noise of our assay (grey bar). Data from Kleinendorst \textit{et al}., 2021 \cite{kleinendorst_genome-wide_2021}. \textbf{(B)} Treatment by M.CviPI does not interfere with endogenous methylation levels in NWCGW contexts. Smoothed scatter plot comparing the average CpG methylation as measured by whole genome bisulfite sequencing in untreated mESC (data from Stadler \textit{et al}., 2011 \cite{stadler_dna-binding_2011}), with the methylation upon treatment by M.CviPI in SMF data. The Pearson correlation coefficient is indicated. \textbf{(C)} Distribution of read coverage over CpGs and GpCs. Histograms showing the distribution of read counts over CpGs (NWCGW context, top) and GpCs (DGCHN context) in 101 bp bins surrounding CpGs (bottom) considered for analysis. Red lines indicate the median coverage. \textbf{(D)} Measure of DNA footprinting in SMF data is highly reproducible across replicates. Smoothed scatter plot comparing the average methylation levels in DGCHN measured by bisulfite sequencing in two independent biological replicates. The Pearson correlation coefficient is indicated. \textbf{(E)} Measure of endogenous DNA methylation in SMF data is highly reproducible across replicates. Smoothed scatter plot comparing the average methylation levels in CpGs (NWCGW context) measured by bisulfite sequencing in SMF in two independent biological replicates. The Pearson correlation coefficient is indicated. \textbf{(F)} Comparison of CpGs distributions at different methylation states. Bar plots showing the distribution of CpGs (NWCGW context) at binned methylation states in SMF data in mESC. Colored in black are those methylation states used for subsequent analyses (10-60\%). \textbf{(G)} Intermediately methylated CpGs are enriched for \emph{cis}-regulatory elements as enhancers. Stacked barplot showing the genomic context annotation for all CpGs (NWCGW context) and intermediately methylated CpGs used for subsequent analyses using publicly available chromHMM annotations clustered into 5 groups (chromHMM data from Pintacuda \textit{et al}., 2017 \cite{pintacuda_hnrnpk_2017}). \textbf{(H)} Most active enhancers show no difference in 5mC between the two separated chromatin accessibility fractions. Violin bar plots showing the CA (top) and 5mC (bottom) at CpGs (NWCGW context) in bulk and in the accessible and inaccessible fractions of molecules. Boxplots show median (black middle line), 25th and 75th percentiles (black boundaries).}
\label{fig:S1}
\end{figure}

\begin{figure}%[tbhp]
  \captionsetup{labelformat=empty}
  \centering
  \makebox[\textwidth]{\includegraphics[width=.8\paperwidth]{Figures/Figure_S2}}
  \caption{}
\end{figure}
\begin{figure}[t]
  \captionsetup{labelformat=adja-page}
  \ContinuedFloat
  \caption{\textbf{(A)} Most CpGs within annotated Imprinting Control Regions (ICRs) show strong methylation differences and are detected as antagonists. Volcano plots depicting the common odds ratio and the p-value of a Cochran-Mantel-Haenszel test as shown in Figure \ref{fig:2}A split by categories: neutral, antagonist and annotated ICR sites. A fraction of CpGs in ICRs show high methylation differences between accessible and inaccessible molecules and, thus, depict a subclass of antagonist sites (n = 82). Those are excluded from the antagonist sites, leaving 3,065 non-ICR antagonist CpGs for downstream analyses. \textbf{(B-C)} Single loci exemplifying the difference between an annotated ICR at the \textit{Dlk1-Gtl2} imprinted locus (B) and a newly identified antagonist locus (C) at allelic resolution. At the antagonist locus, both alleles show methylation heterogeneity and antagonism with chromatin accessibility. At the annotated ICR locus, the two alleles are differentially methylated and differentially accessible. SMF was performed in F1 hybrid (129/CAST) mESCs and the chromatin accessibility and DNA methylation were analysed for both alleles separately. Top panels depict average SMF signals (1 - methylation \%) of individual GpCs in the maternal (black dots) or paternal (grey dots) allele. On top, location and methylation status of the CpGs in the two alleles are shown. Lower panels show single DNA molecules for the two alleles separately (top: maternal, bottom: paternal) sorted by chromatin accessibility into an accessible (grey) and inaccessible (black) fraction. In the left panel, every column is a single GpC dinucleotide depicting its accessibility status. In the right panel, every column is a single CpG dinucleotide depicting its methylation status (light blue: unmethylated, dark blue: methylated). The number of DNA molecules analysed for each allele at both loci is indicated. \textbf{(D)} Antagonist sites harbor characteristic chromatin marks of enhancers. Heatmap displaying the ChIP-seq counts for H3K27ac, H3K27me3, H3K4me1, H3K4me3, CTCF as well as chromatin accessibility as measured by DNase hypersensitive sites (DHS) from publicly available data in a 4 kb window centered on 3065 antagonist CpGs. \textbf{(E)} Neutral and antagonist sites have low CpG densities. Violin box plots showing the CpG density distribution for neutral (n = 23,656), antagonist (n = 3,065) and ICR sites (n = 153) calculated by the number of CpGs per 100 bp in a 400 bp window surrounding the CpGs in each category. Shown neutral sites are high confidence neutrals with minor methylation difference between fractions (common odds ratio between 0.9 and 1.1). Boxplots show median (black middle line), 25th and 75th percentiles (black boundaries). \textbf{(F)} Neutral and antagonist sites show intermediate chromatin accessibility. Violin box plots showing the distribution of average accessibility in a 100 bp window surrounding CpGs in the neutral (n = 23,656), antagonist (n = 3,065) and ICR sites (n = 153) category measured by SMF. Shown neutral sites are high confidence neutrals with minor methylation difference between fractions (common odds ratio between 0.9 and 1.1). Boxplots show median (black middle line), 25th and 75th percentiles (black boundaries). \textbf{(G)} Neutral and antagonist sites show similar levels of 5-hydroxymethylation (5hmC). Violin box plots showing the distribution of 5hmC signal measured by hMeDIP-seq in a 500 bp window surrounding all CpGs or CpGs in the neutral (n = 23,656) or antagonist (n = 3,065) category. Shown neutral sites are high confidence neutrals with minor methylation difference between fractions (common odds ratio between 0.9 and 1.1). Plotted is the log2 of hMeDIP signal as counts per millions with added pseudo-count (cpm+pc). Boxplots show median (black middle line), 25th and 75th percentiles (black boundaries). hMeDIP-seq data from Xu \textit{et al}., 2011 \cite{xu_genome-wide_2011}.
}
\label{fig:S2}
\end{figure}
\clearpage

\begin{figure*}[t]
\centering
 \makebox[\textwidth]{\includegraphics[width=.8\paperwidth]{Figures/Figure_S3}}
\caption{\textbf{(A)} Neutral and antagonist sites show comparable intermediate DNA methylation levels in mESCs measured by SMF. Violin box plots showing the distribution of average CpG methylation of the neutral (n = 23,656) or antagonist (n = 3,065) category. Shown neutral sites are high confidence neutrals with minor methylation difference between fractions (common odds ratio between 0.9 and 1.1). Boxplots show median (black middle line), 25th and 75th percentiles (black boundaries). \textbf{(B)} Global gain of DNA methylation in TET TKO for CpGs with intermediate DNA methylation. Smoothed scatter plot comparing the average methylation levels in NWCGW measured by bisulfite sequencing in SMF in wild type (WT) mESC and an isogenic line where the TET enzymes have been knocked out (TET TKO). The Pearson correlation coefficient is indicated. \textbf{(C-D)} Chromatin accessibility is a function of DNA methylation levels at antagonist sites. Scatterplot comparing the changes in chromatin accessibility and DNA methylation in (C) DNMT TKO (n = 1,964) and (D) TET TKO (n = 1,275) for antagonist sites. The decrease in accessibility is anti-correlated with the increase in DNA methylation at the locus. The regression line and Pearson correlation coefficients are indicated. \textbf{(E)} Some dysregulated genes are regulated by multiple antagonist enhancers. Barplot showing the number of dysregulated genes in RNA-seq (upregulated in DNMT TKO, downregulated in TET TKO) that have one or multiple annotated antagonist sites. \textbf{(F)} Some antagonist enhancers are annotated to multiple dysregulated genes. Barplot showing the number of antagonist sites that are annotated to multiple dysregulated genes (upregulated in DNMT TKO, downregulated in TET TKO). \textbf{(G)} Identification of two antagonist CpGs at the \textit{Sox2} enhancer cluster. Upper panel: Genome browser track displaying the chromatin marks and the average CpG methylation as measured by WGBS and average methylation of intermediately methylated NWCGWs as measured by SMF around an intergenic antagonist enhancer at the \textit{Sox2} enhancer cluster locus. Two CpGs at the locus were identified as antagonist (pink rectangles on top). Middle and lower panels: Single locus example of the DNA methylation and accessibility of individual molecules at one of the antagonist CpGs contained within the \textit{Sox2} enhancer cluster. Same representation as in Figure \ref{fig:2}B. RNA-seq data in DNMT TKO from Domcke \textit{et al}., 2015 \cite{domcke_competition_2015} and in TET TKO from Huang \textit{et al}., 2021 \cite{huang_chromosomal_2021}.}
\label{fig:S3}
\end{figure*}
\clearpage
\newpage

\begin{figure*}[t]
\centering
 \makebox[\textwidth]{\includegraphics[width=.8\paperwidth]{Figures/Figure_S4}}
\caption{\textbf{(A)} GO term analysis of cell-type specific antagonist enhancers identifies involvement in cell-type specific processes. Dot plots showing the adjusted p-value of top GO term processes (fold change > 2) identified by GREAT analysis of cell-type specific antagonist enhancers in neural progenitors (NP), myoblasts (C2C12) and erythrocytes (MEL). Triangles indicate cell-type specific processes for each cell line (black: highly relevant, grey: relevant). \textbf{(B)} Cell-type specific antagonist sites are enriched for E-box motifs as well as for motifs of cell-type specific transcription factors. Dot plot showing the z-score of the -log of the adjusted p-value and the percentage of enrichment of motifs identified with HOMER motif enrichment analysis of cell-type specific antagonist enhancers in embryonic stem cells (ES), neural progenitors (NP), myoblasts (C2C12) and erythrocytes (MEL). Black triangles indicate cell-type specific motifs.
}
\label{fig:S4}
\end{figure*}
\clearpage
\newpage

\begin{figure*}%[tbhp]
  \captionsetup{labelformat=empty}
  \centering
  \makebox[\textwidth]{\includegraphics[width=.7\paperwidth]{Figures/Figure_S5}}
  \caption{}
\end{figure*}
\begin{figure*}[th]
  \captionsetup{labelformat=adja-page}
  \ContinuedFloat
  \caption{\textbf{(A)} Single locus example of the methylation difference between transcription factor (TF) bound and nucleosome bound molecules at a neutral CTCF site. The TF bound fraction is not depleted from methylated molecules. Representation as in Figure \ref{fig:5}C. \textbf{(B)} Validation of the antagonism observed at CTCF binding sites. Scatterplot comparing the difference in DNA methylation (5mC) between the TF bound and the nucleosome occupied fraction, with the 5mC enrichment upon CTCF ChIP-bis. Each dot is a CTCF binding sites covered in both experiments. The enrichment of CTCF for unmethylated molecules at antagonist binding sites (red dots) is observed by both methods. ChIP-bis data from Feldmann \textit{et al}., 2013 \cite{feldmann_transcription_2013}. \textbf{(C)} Distribution of 5mC at the binding sites of various TFs. Histograms describing the average methylation at the binding sites of TFs. The binding of all TFs is enriched for unmethylated regions (grey), yet CTCF, Max-Myc, NFYA and REST are also binding at regions with intermediate methylation (black). Conservative TF motif names have been used. \textbf{(D)} Differential methylation between TF bound molecules and nucleosome occupied molecules at a subset of TF binding sites. Scatterplot showing the average 5mC for the fraction of molecules showing a TF footprint against those showing a nucleosome footprint at the same locus. The analysis is shown for each TF separately. The color code indicates the category of the binding site (pink – antagonist*, black – neutral*, grey – unmethylated). Conservative TF motif names have been used. \textbf{(E-F)} TF binding at antagonist and neutral sites is confirmed by ChIP-seq data. Violin box plots showing the ChIP-seq signal (log2 of the read counts per million (cpm)) at all binding sites and antagonist* and neutral* binding sites used in the analyses for (E) CTCF and (F) Myc. Boxplots show median (black middle line), 25th and 75th percentiles (black boundaries). The vertical line depicts the boundary of the 50th percentile. CTCF ChIP-seq data from Stadler \textit{et al}., 2011 \cite{stadler_dna-binding_2011}. Myc ChIP-seq data from Chronis \textit{et al}., 2017 \cite{chronis_cooperative_2017}, Das \textit{et al}., 2014 \cite{das_distinct_2014} and Mor \textit{et al}., 2018 \cite{mor_neutralizing_2018}). \textbf{(G-H)} TF occupancy is a function of 5mC levels at antagonist sites. Scatterplots comparing the changes in TF occupancy and 5mC in WT mESC at neutral* (top) and antagonist* (bottom) sites for (G) CTCF and (H) Max-Myc binding sites. The decrease in TF binding is anti-correlated with the increase in 5mC at the locus at antagonist sites. The regression line and the Pearson correlation coefficients are displayed. \textbf{(I)} CpG occurrence at specific positions of the intermediately methylated CTCF motifs. Barplot displaying the frequency of CpGs in antagonist* (pink) and neutral* (grey) sites. Position \#7 frequently has a CpG in antagonist binding sites, but not in neutral* binding sites. This partially overlaps with the motifs positions that contain a CpG in the binding site that are reactivated upon deletion of 5mC (position 5, 7, 15, indicated by a star) \cite{maurano_role_2015}. \textbf{(J)} CpG occurrence at specific positions of the intermediately methylated Max-Myc motifs. Barplot displaying the frequency of CpGs in antagonist* (pink) and neutral* (grey) sites. CpGs mainly occur at position \#6, with no significant difference between neutral and antagonist binding sites. \textbf{(K-L)} TF occupancy is a function of 5mC levels at antagonist sites. Scatterplots comparing the change in TF occupancy and the change in 5mC from WT to TET TKO at (K) neutral* and (L) antagonist* sites. The decrease in occupancy is anti-correlated with the increase in 5mC at the locus. The regression line, Pearson correlation coefficients and coefficient of determination are displayed. Neutral* and antagonist* sites in this plot are defined with soft criteria using only the common odds ratio of the Cochran-Mantel-Haenszel test as indicated.}
\label{fig:S5}
\end{figure*}
\clearpage

\end{document}
