## Supplementary figures and images for "Single molecule multi-omics reveals context-dependent regulation of enhancers by DNA methylation"

### Figure_1.png

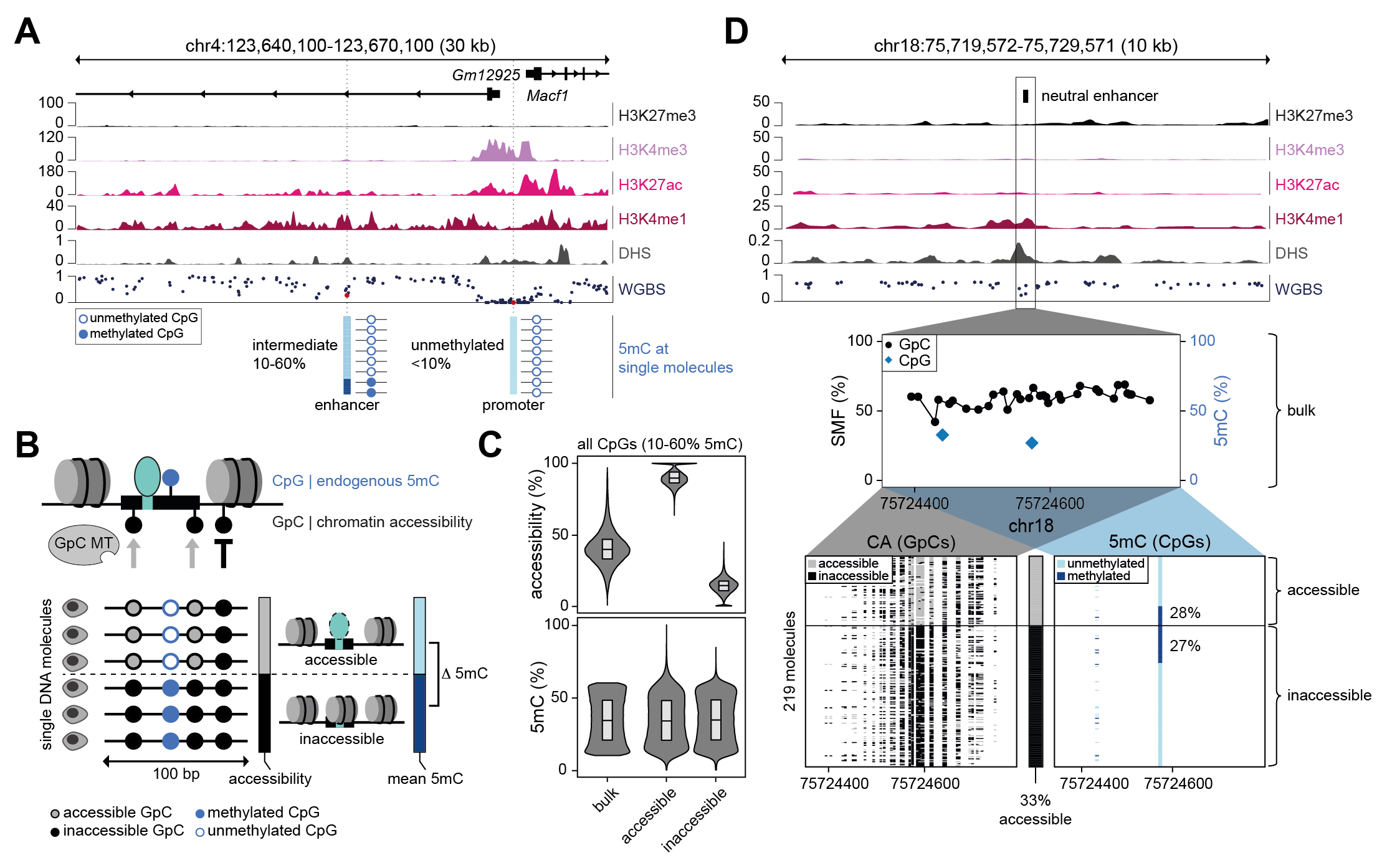

### Figure_2.png

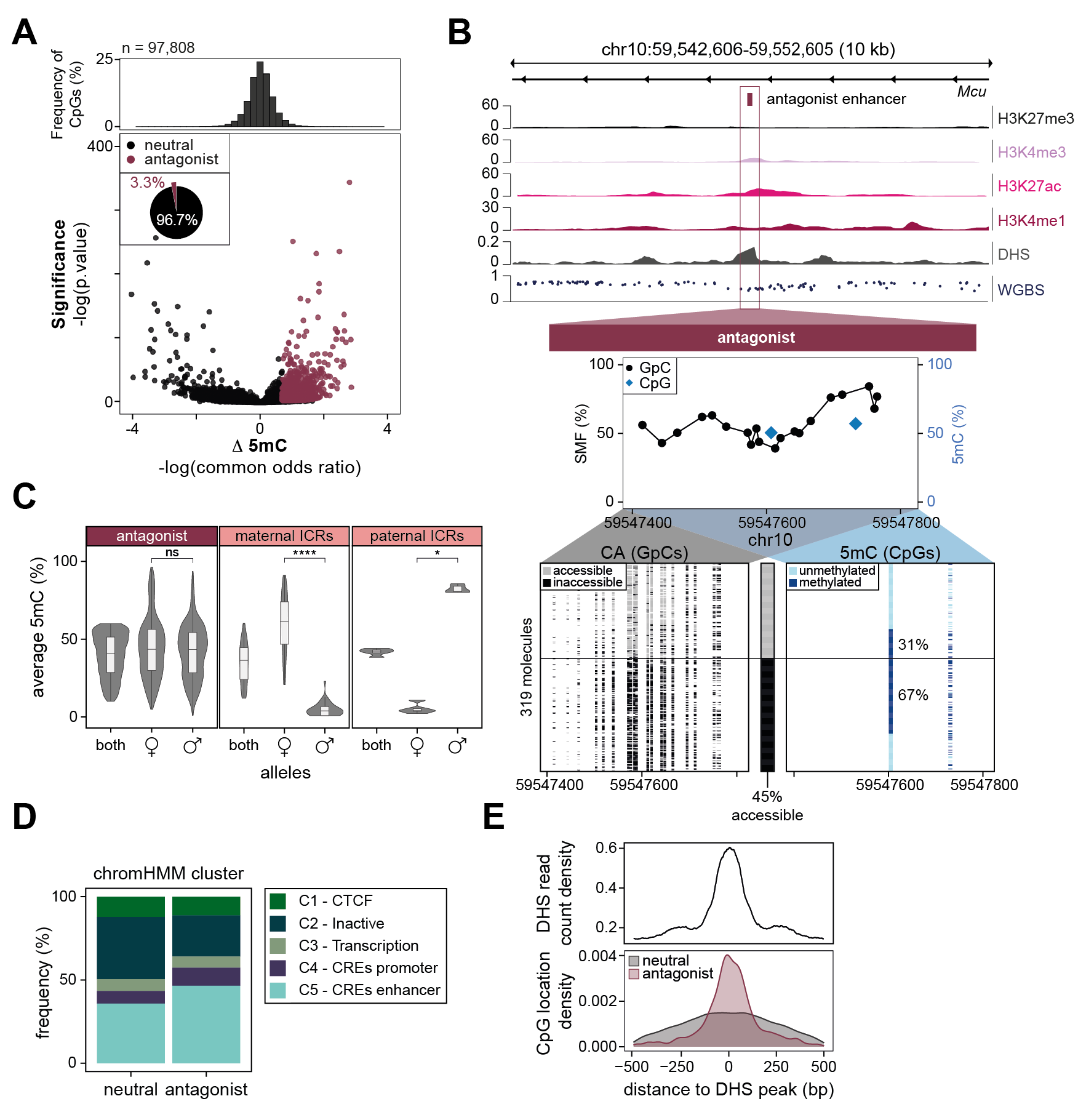

### Figure_3.png

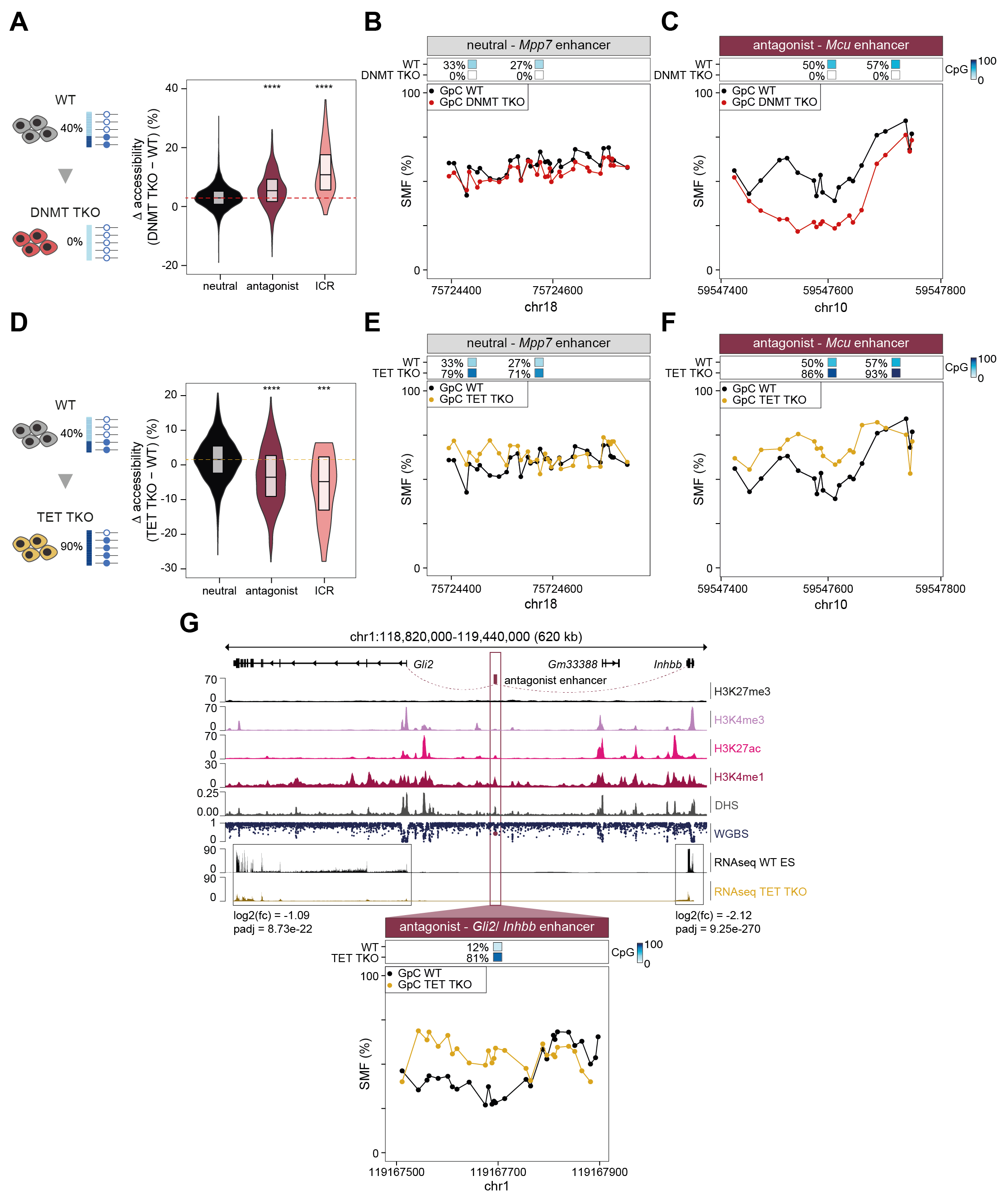

### Figure_4.png

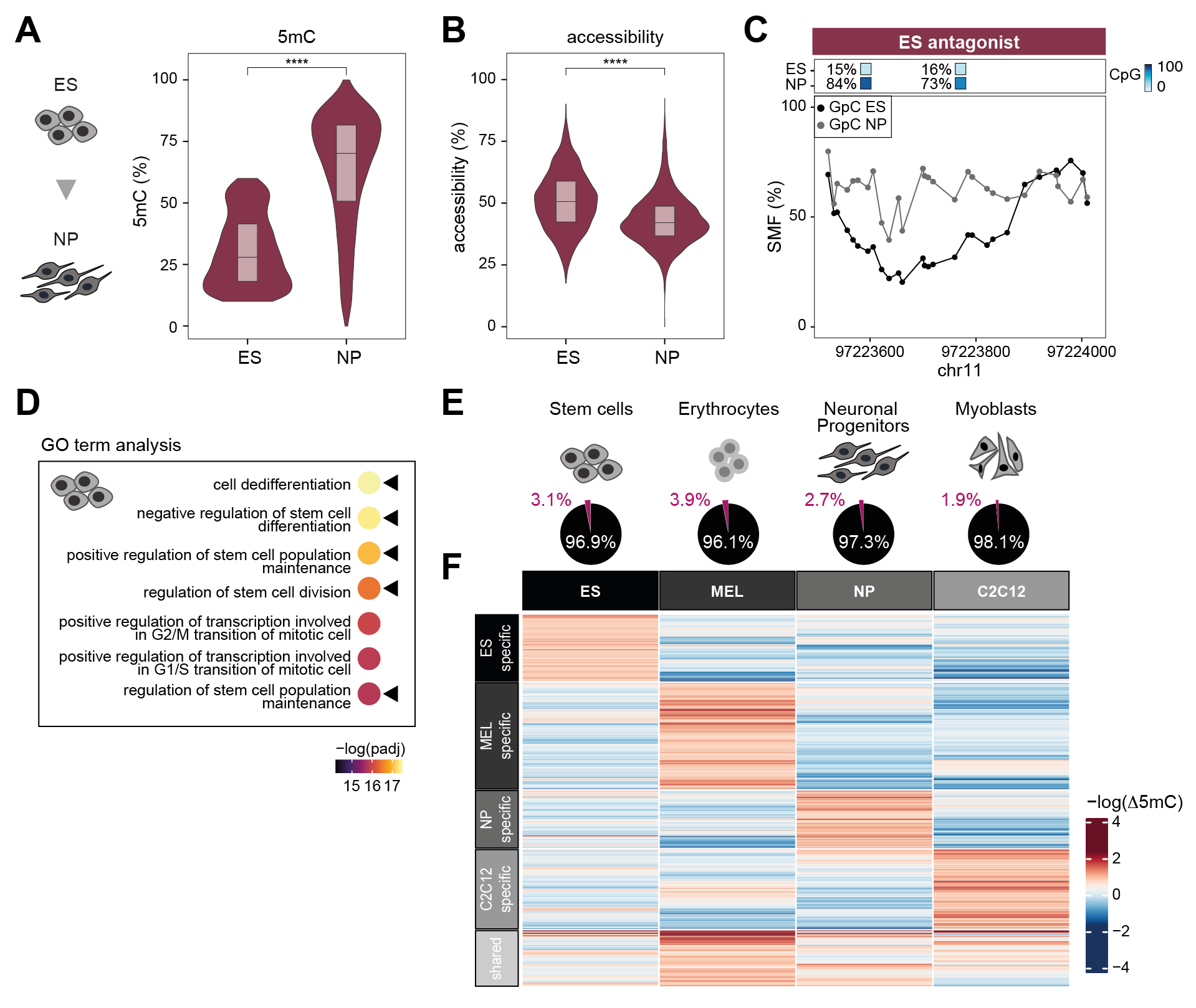

### Figure_5.png

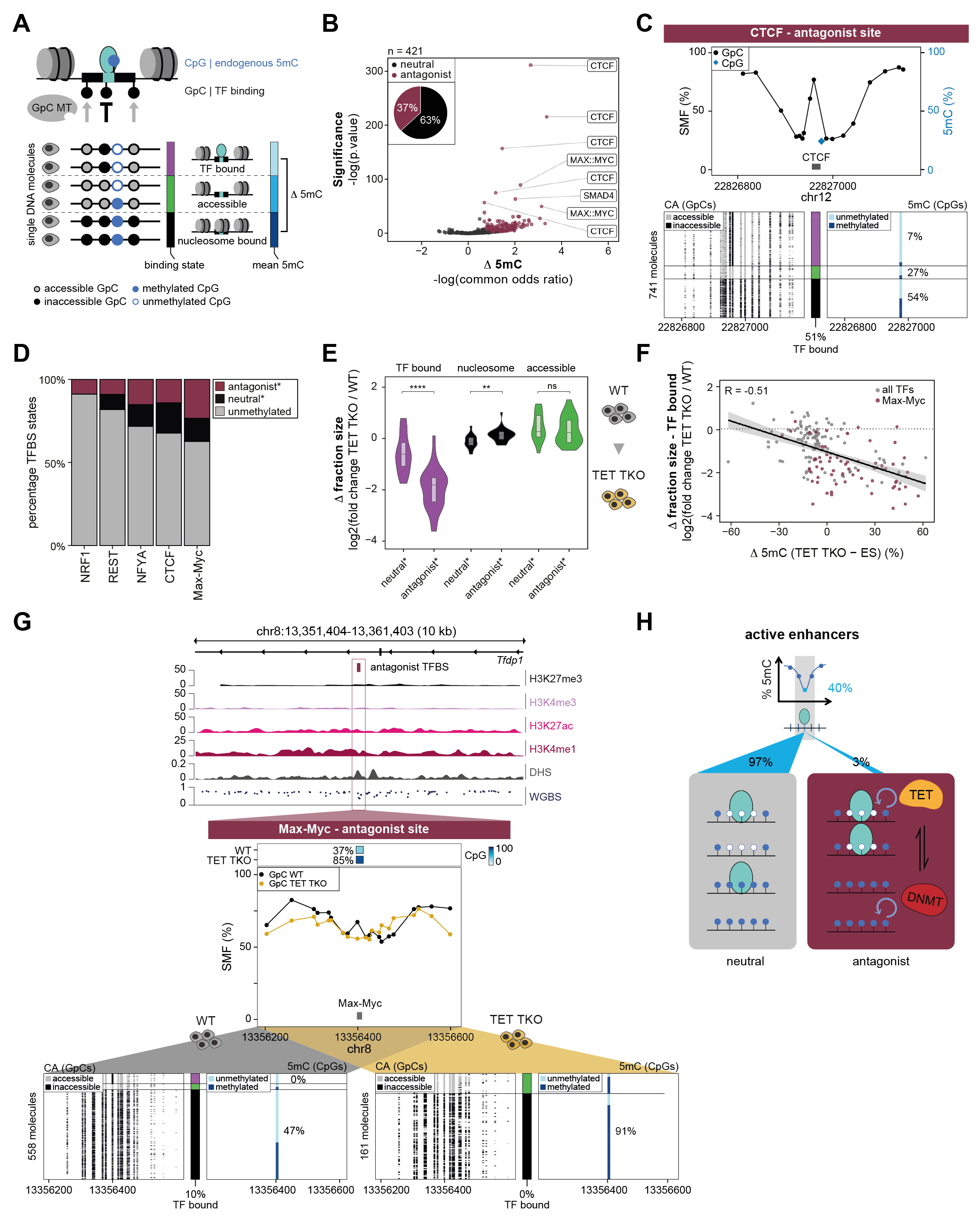

### Figure_S1.png

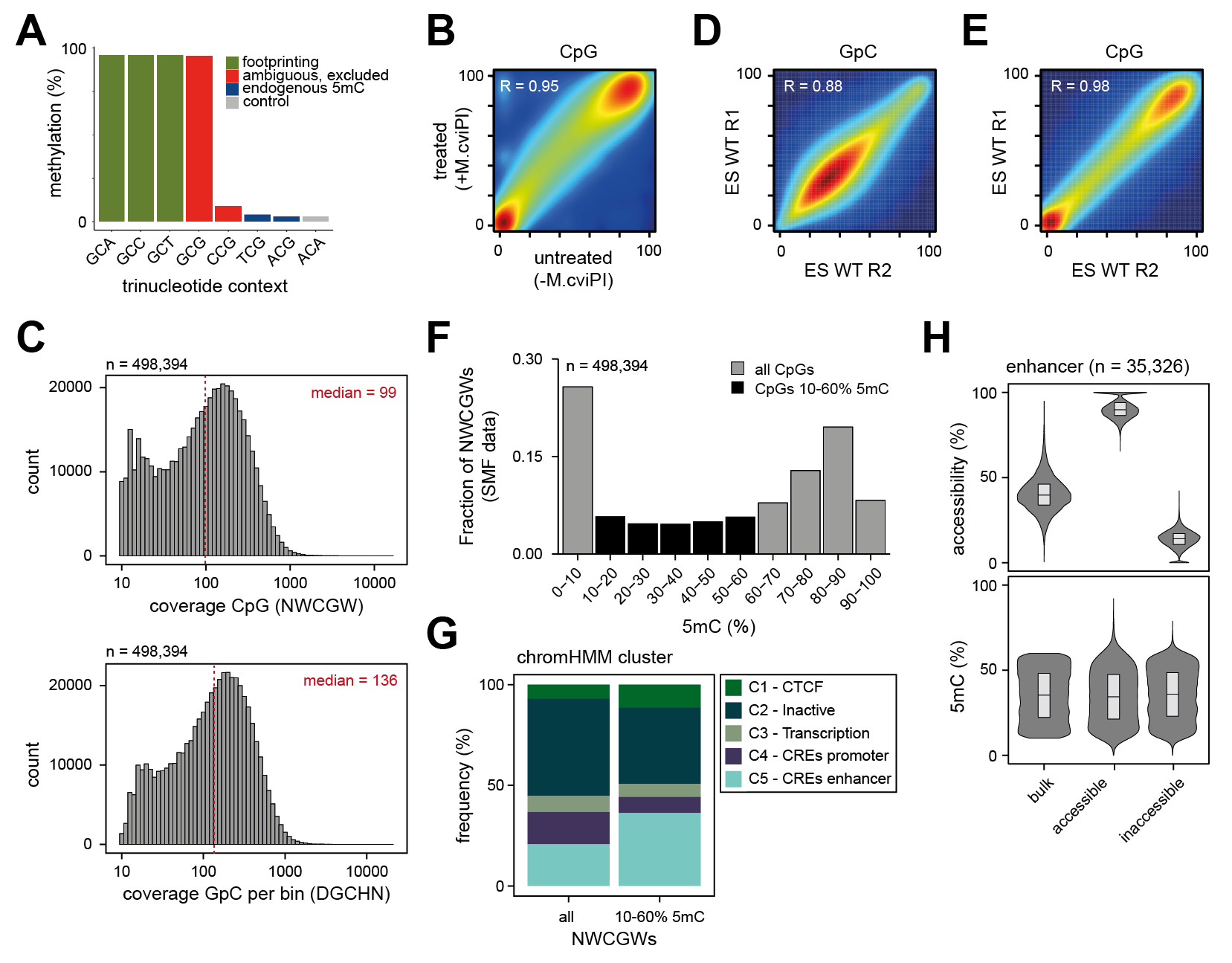

### Figure_S2.png

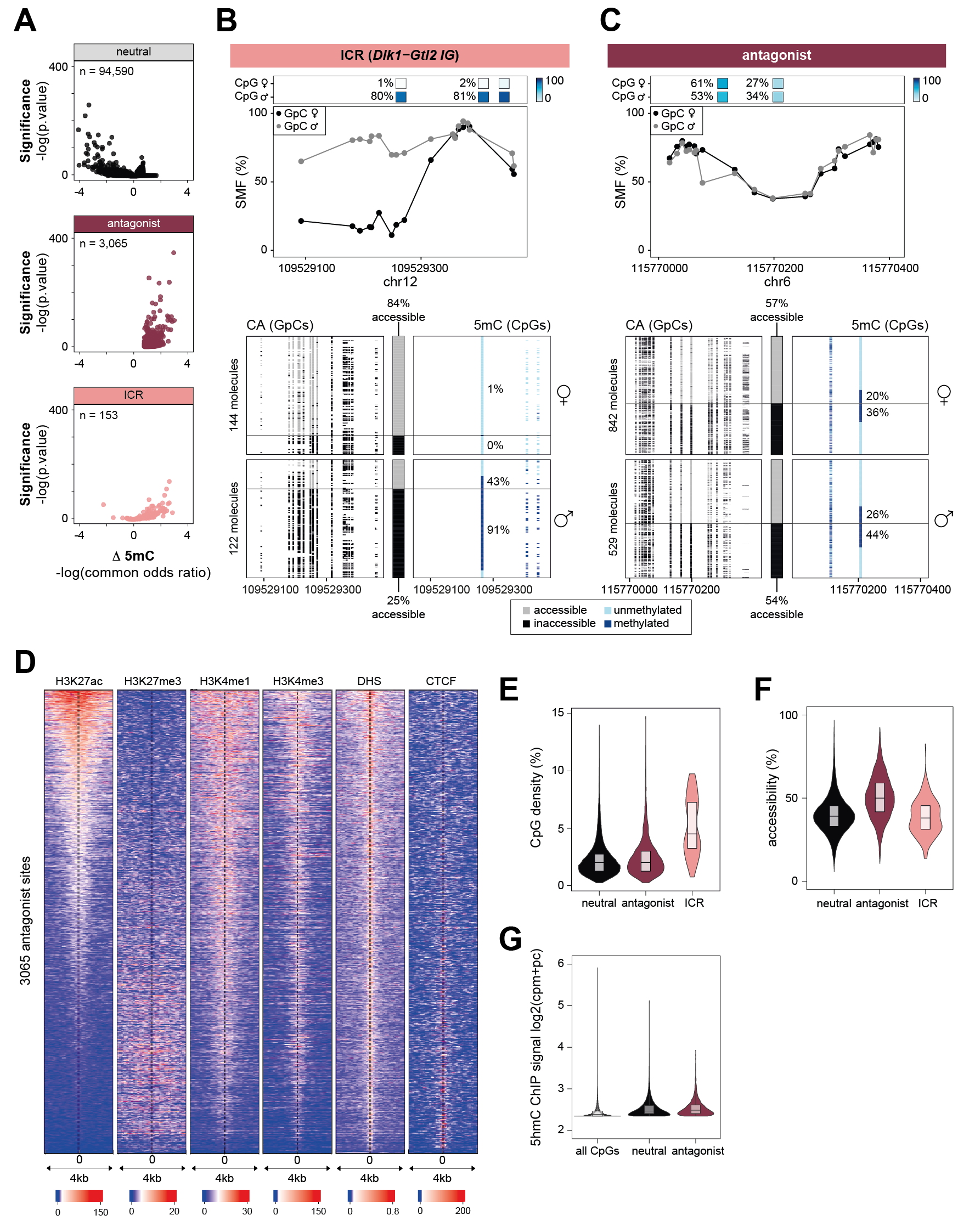

### Figure_S3.png

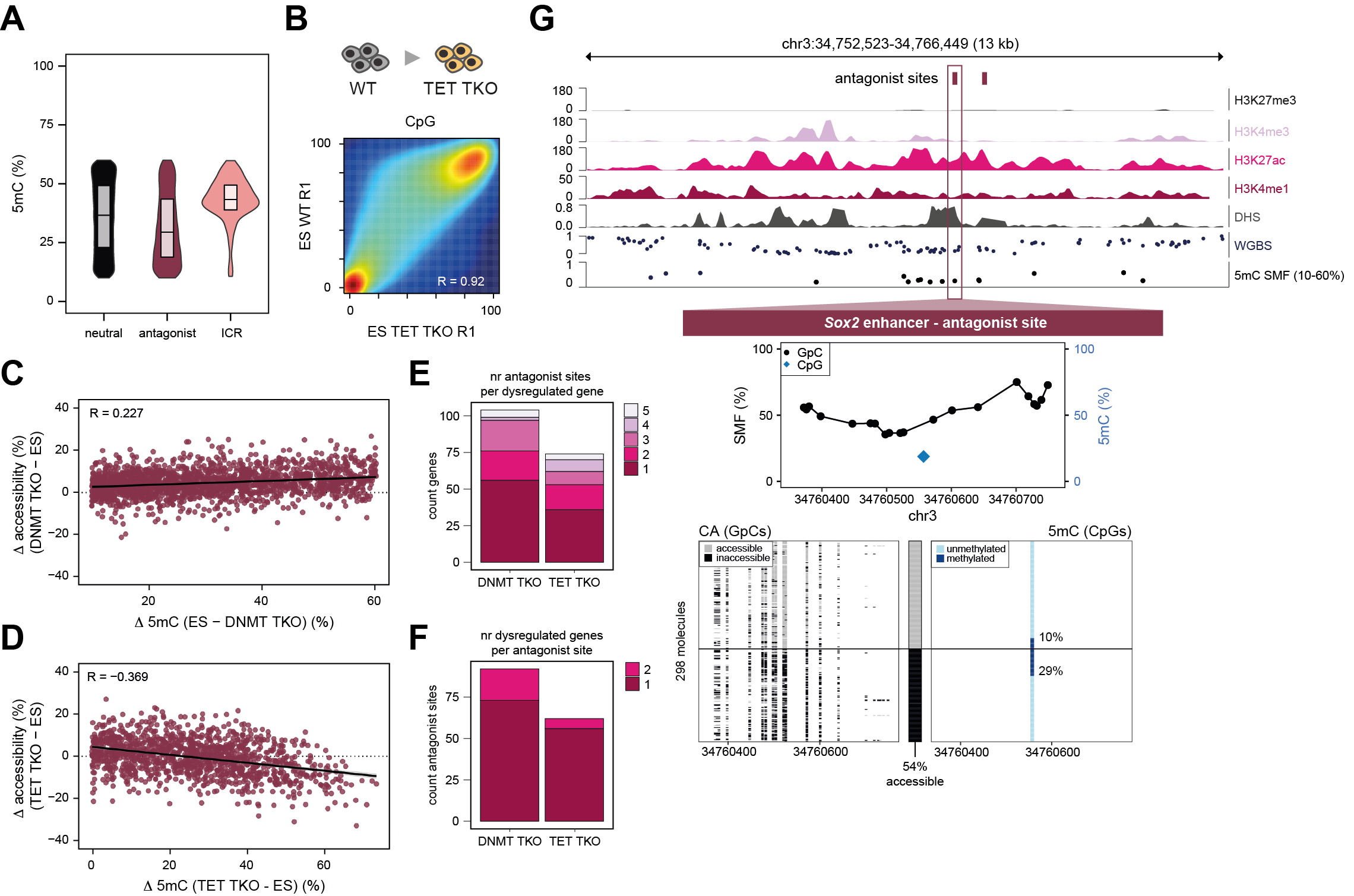

### Figure_S4.png

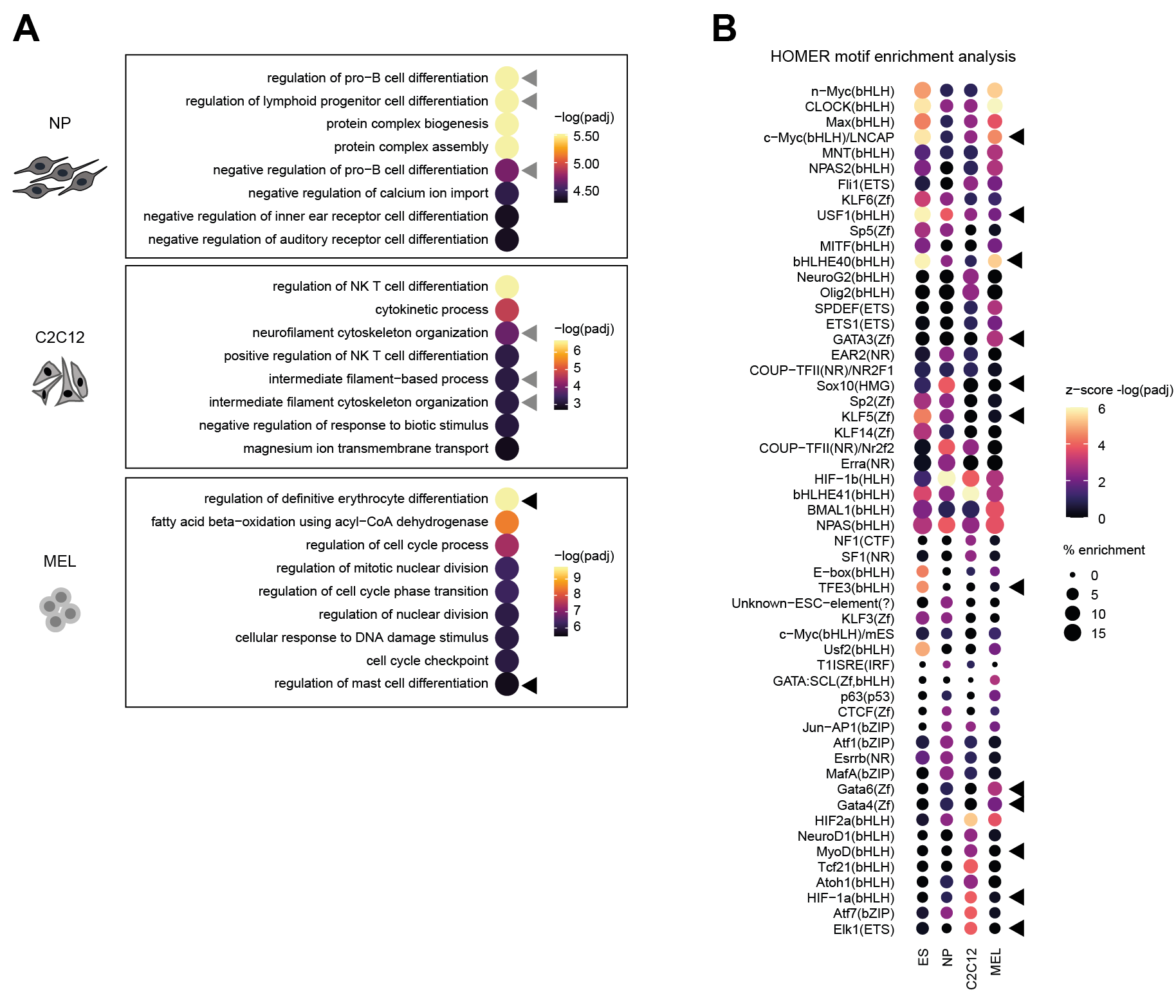

### Figure_S5.png

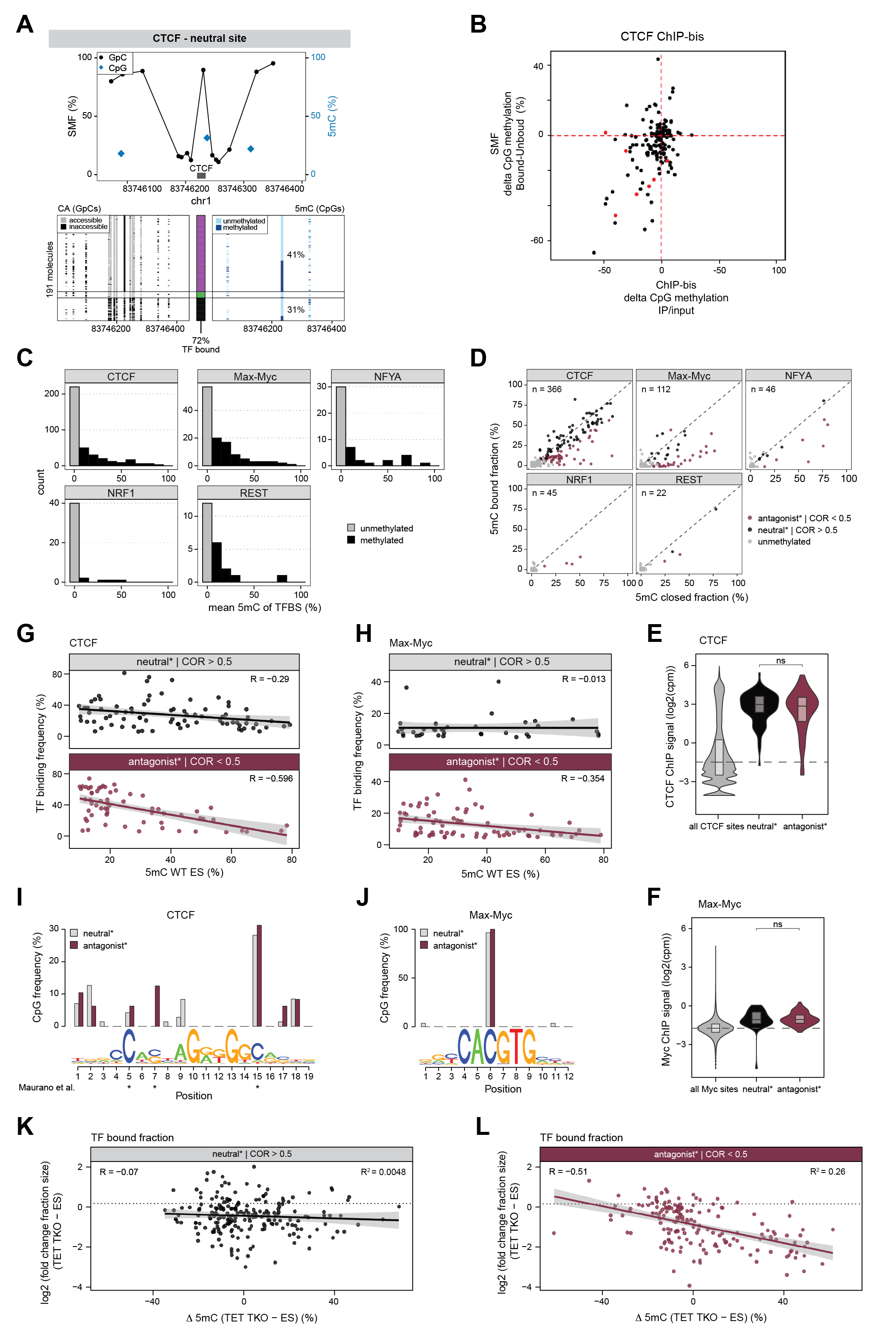
